# Microbial community of recently discovered Auka vent field sheds light on vent biogeography and evolutionary history of thermophily

**DOI:** 10.1101/2021.08.02.454472

**Authors:** Daan R. Speth, Feiqiao B. Yu, Stephanie A. Connon, Sujung Lim, John S. Magyar, Manet E. Peña, Stephen R. Quake, Victoria J. Orphan

**Author notes:** Max Planck Institute for Marine Microbiology, Bremen, Germany (Daan R. Speth).

## Abstract

Hydrothermal vents have been key to our understanding of the limits of life, and the metabolic and phylogenetic diversity of thermophilic organisms. Here we used environmental metagenomics combined with analysis of physico-chemical data and 16S rRNA amplicons to characterize the diversity, temperature optima, and biogeographic distribution of sediment-hosted microorganisms at the recently discovered Auka vents in the Gulf of California, the deepest known hydrothermal vent field in the Pacific Ocean. We recovered 325 metagenome assembled genomes (MAGs) representing 54 phyla, over 1/3 of the currently known phylum diversity, showing the microbial community in Auka hydrothermal sediments is highly diverse. Large scale 16S rRNA amplicon screening of 227 sediment samples across the vent field indicates that the MAGs are largely representative of the microbial community. Metabolic reconstruction of a vent-specific, deeply branching clade within the Desulfobacterota (Tharpobacteria) suggests these organisms metabolize sulfur using novel octaheme cytochrome-c proteins related to hydroxylamine oxidoreductase. Community-wide comparison of the average nucleotide identity of the Auka MAGs with MAGs from the Guaymas Basin vent field, found 400 km to the Northwest, revealed a remarkable 20% species-level overlap between vent sites, suggestive of long-distance species transfer and sediment colonization. An adapted version of a recently developed model for predicting optimal growth temperature to the Auka and Guaymas MAGs indicates several of these uncultured microorganisms could grow at temperatures exceeding the currently known upper limit of life. Extending this analysis to reference data shows that thermophily is a trait that has evolved frequently among Bacteria and Archaea. Combined, our results show that Auka vent field offers new perspectives on our understanding of hydrothermal vent microbiology.

## Introduction

Microbial communities at hydrothermal vents have long been of interest for their impact on localized productivity and nutrient cycling in the deep ocean, as surface expressions of the subsurface biosphere, and as potential analogs for ocean life on icy moons. These chemosynthetic communities differ strongly from the communities inhabiting the surrounding seafloor, but the variation between different hydrothermal areas is not well understood. Based on host lithology, hydrothermal areas can be classified into three groups: basalt-hosted, ultramafic, and sediment-hosted. The majority of well-studied high temperature vents are basalt-hosted, with hydrothermal fluid directly discharged from fissures into the overlying seawater (Dick 2019). In contrast, sediment-hosted hydrothermal vents such as those found in Guaymas Basin are distinctive for the interaction of the superheated fluid with overlying sediment. This interaction further alters the fluid composition through incorporating thermally-degraded organic compounds during advection to the seafloor, resulting in steep temperature gradients in the sediments and near surface oil production (Procesi et al. 2019).

This additional complexity makes sediment-hosted vent fields attractive study sites for microbial ecology (Teske 2020). Indeed, Guaymas Basin has proven to be a particularly rich source for discovery of novel metabolic capabilities of thermophilic microorganisms. Examples include thermophilic anaerobic oxidation of methane coupled to sulfate reduction by consortia of *Desulfofervidus sp.* bacteria and ANME-1 archaea (Holler et al. 2011; Schouten et al. 2003), anaerobic butane degradation by the sulfate-reducing bacterium *Desulfosarcina* BuS5 (Kniemeyer et al. 2007), and anaerobic butane oxidation by consortia of *Synthrophoarchaeum sp.* and *Desulfofervidus sp*. Bacteria (Laso-Pérez et al. 2016). In addition, some of the most extreme hyperthermophiles, *Methanopyrus kandleri* and *Pyrodictium abyssi*, with maximum measured growth temperatures of 122°C and 110°C, respectively, have been isolated from Guaymas Basin sediments and chimneys (Pley et al. 1991; Kurr et al. 1991; Takai et al. 2008). The outsized role of Guaymas Basin in discovery of microbial processes makes the recent discovery of the Auka vent field, a second sediment hosted hydrothermal vent system along the same fault in the Gulf of California (Goffredi et al. 2017; Paduan et al. 2018; Espinosa-Asuar et al. 2020), especially exciting, as it provides a unique opportunity for comparative analyses of sediment hosted hydrothermal vent systems.

Auka is located at >3650m water depth in the Southern Pescadero Basin, a pull-apart basin at the southern tip of the Gulf of California and 400 kilometers southeast of Guaymas Basin. The composition of the hydrothermal fluids at both sites is similar. The fluids are slightly acidic (pH 6), with high concentrations of methane (81 / 16 mmol kg^-1^), hydrogen sulfide (10.8 / 6 mmol kg^-1^), and carbon dioxide (49.2 / 43 mmol kg^-1^) at Auka and Guaymas respectively, and comparatively low hydrogen gas concentration (2 mmol kg^-1^ at Auka) (Von Damm et al. 1985; Welhan 1988; Paduan et al. 2018). The temperature of the fluids measured at chimney orifices is close to 300°C at both locations. Due to these high temperatures, fluids advecting through the sediments at both sites contain thermogenic hydrocarbons, originating from the catagenesis of sediment organic matter.

While the similarities between both sites are striking, there are stark differences as well. At 3650m Auka is the deepest known hydrothermal vent system in the Pacific Ocean, and more than twice as deep as Guaymas Basin. The thicker sediment cover (700 - 1000m, (Lizarralde et al. 2007; Procesi et al. 2019), results in higher load of thermogenic hydrocarbons than observed at Pescadero Basin, where the sediment thickness is estimated to be less than 50 m in the areas directly adjacent to the hydrothermal mounds (Paduan et al. 2018). This combination of overlapping and contrasting conditions between the two sites makes them prime targets for comparative analysis of their microbial communities to elucidate the factors shaping microbial populations at either site.

To characterize the sediment microbial community at the Auka vent field, we analyzed 325 MAGs recovered from metagenomic sequencing combined with a 16S rRNA gene-based diversity survey. The diversity and genomic similarity was then compared between Auka MAGs and those recently reported from Guaymas Basin sediments (Dombrowski et al. 2017; Dombrowski, Teske, and Baker 2018; Seitz et al. 2019), providing important insights into shared microbial species and patterns in vent biogeography. Focusing on one of the novel lineages shared between Auka and Guaymas, a deep-branching lineage of Desulfobacterota, named Tharpobacteria, we conducted a detailed analysis of the metabolic potential within this novel vent-associated clade. Finally, by adapting a genome-based optimal growth temperature prediction model (Sauer and Wang 2019) we determined the distribution of mesophily-hyperthermophily across these diverse lineages, exploring the role temperature plays in shaping microbial communities in Gulf of California hydrothermal vent sediments and, more broadly, evolutionary patterns of thermophily across the tree of life.

## Results

The Auka vent field is located at the western edge of the Southern Pescadero Basin and occupies an area of approximately 200 by 600 meters (Figure 1). There are five prominent sites of vigorous fluid venting referred to as P-vent, C-vent, Z-vent, Diane’s vent, and Matterhorn, each characterized by prominent calcite chimneys (Figure 1; (Paduan et al. 2018). Between these vents lie extensive carbonate platforms dotted with centimeter scale sites of hydrothermal fluid discharge supporting localized clumps of chemosynthetic *Oasisia* tubeworms (Goffredi et al. 2017). These hydrothermal features are sediment-hosted, with sediment thickness in the area directly surrounding the chimneys and carbonate platforms estimated to be less than 50 m (Paduan et al. 2018). Throughout the vent field, but primarily near Diane’s vent and south of Z-vent, dispersed microbial mats covering the sediment surface indicate widespread advective hydrothermal discharge through the sediment (Figure 1). The microbial mats exhibited a range of colors: pink, gray/white, yellow, and contain localized bright white spots surrounding focused flow of shimmering hydrothermal fluid (Supplemental figure 1). Sediment temperatures at 30cm depth were elevated relative to the bottom seawater 2.4°C, ranging from 10°C to 177°C, with the highest temperatures measured below yellow-colored mats and beneath the bright white zones associated with visible hydrothermal fluid discharge. The release of oil droplets was observed during sediment coring to the south of Z-vent within a microbial mat, consistent with near-surface petroleum production within the sediment.

**Figure 1.**
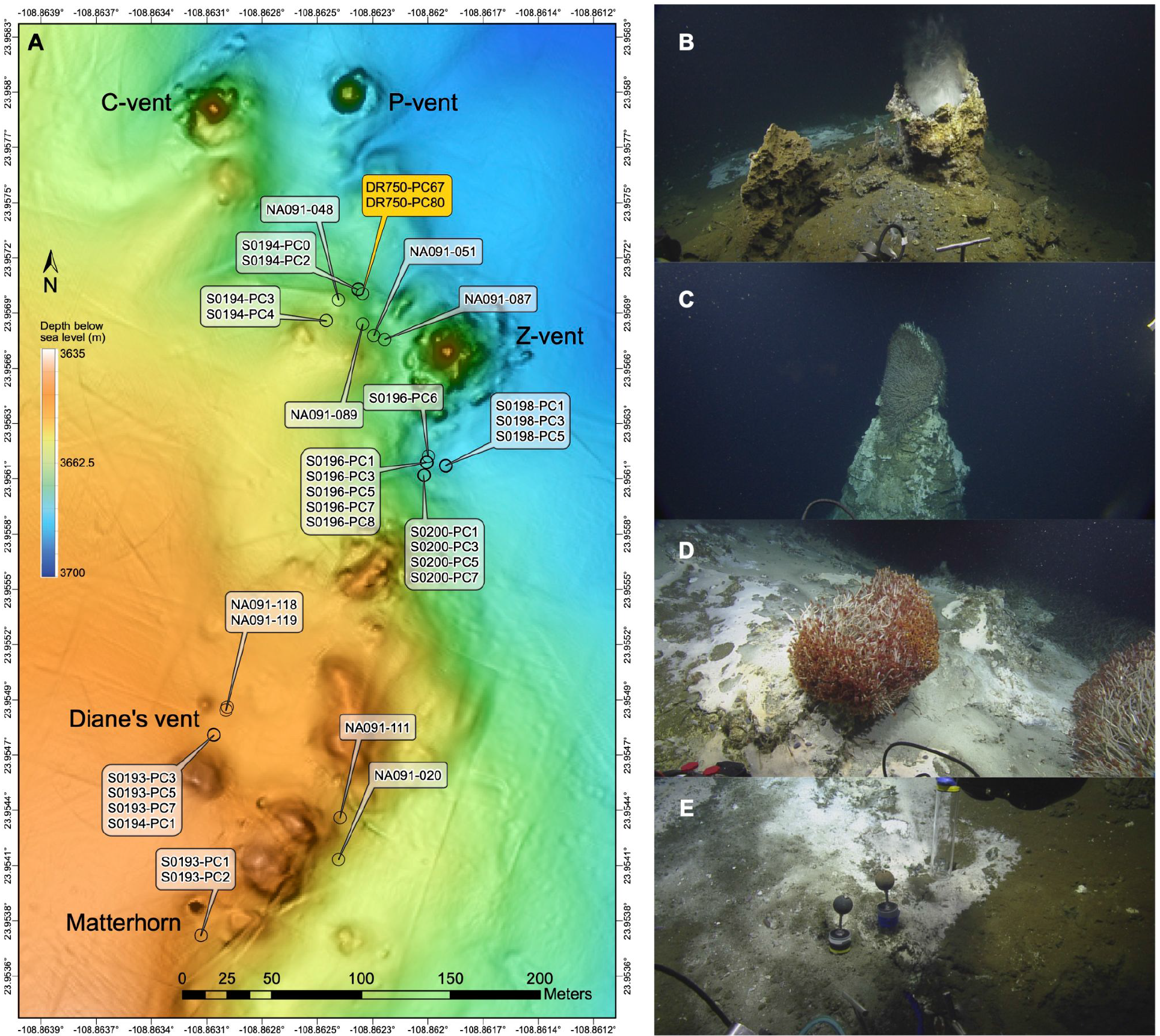
Overview of sampling area and impressions of Auka vent field. **A)** Bathymetric map of Auka vent field with the most prominent sites of focused hydrothermal venting labeled by name. The sampling locations of push cores analyzed in this study are indicated, with the cores used for metagenomic sequencing highlighted in orange. **B)** Diane’s vent, a chimney with distinctive clear hydrothermal fluid discharge clearly visible. **C)** Top of the Matterhorn, a ∼10m high free standing chimney, with the area around the central orifice fully covered in *Oasisia sp.* tubeworms. Microbial mat and shallow flange structures are visible on the sides of the chimney, indicative of diffuse fluid discharge through the chimney wall. **D)** Carbonate platform covered in white and grey microbial mat with *Oasisia sp.* tubeworms (∼20cm tall) clustered around a localized spot of hydrothermal fluid discharge. This is a representative example of the unlabeled elevated mounds shown in panel A. **E)** Sediment covered in microbial mat (at Diane’s vent) with heterogeneity of colors and textures indicating centimeter scale spatial heterogeneity in fluid diffusion through the sediment. Pushcore locations in panel A correspond to areas with prevalent microbial mats.

To characterize the microbial community of Auka vent field, we performed shotgun metagenomic sequencing on sediments from two paired cores, DR750-PC67 and DR750-PC80, collected approximately 50 m from the main Z vent chimney (Figure 1). The cores were collected within sediment covered by patchy yellow microbial mat, 5 cm from visible discharge of hydrothermal fluid and associated bright white spot on the sediment. DR750-PC80 was inserted closest to the fluid discharge, with DR750-PC67 directly adjacent (<2 cm apart), in a straight line away from the white spot (Supplemental Figure 1). Each core was sectioned into two 7 cm horizons (0-7cm and 7-14cm), resulting in four samples. To assess whether there were differences in the microbial community attached to, or forming, larger aggregates in the sediments, <10 µm filtrate of a subsample of each horizon was processed for DNA extraction; doubling the total number of metagenomic samples to eight.

After sequencing, assembly, and binning, we retrieved 331 metagenome assembled genomes (MAGs) with estimated completeness over 50%. Recovered MAGs were highly diverse, including 212 Bacteria and 119 Archaea and representing 54 different phyla based on the genome taxonomy database (GTDB) assignment (Chaumeil et al. 2019) (Figure 2, Supplemental Data S5). 105 of these MAGs were estimated to be >90% complete with less than 5% contamination, and 111 MAGs contained (fragments of) a 16S rRNA gene (Supplemental Data S5) allowing direct comparison with the more extensive 16S rRNA amplicon survey data from Auka.

**Figure 2.**
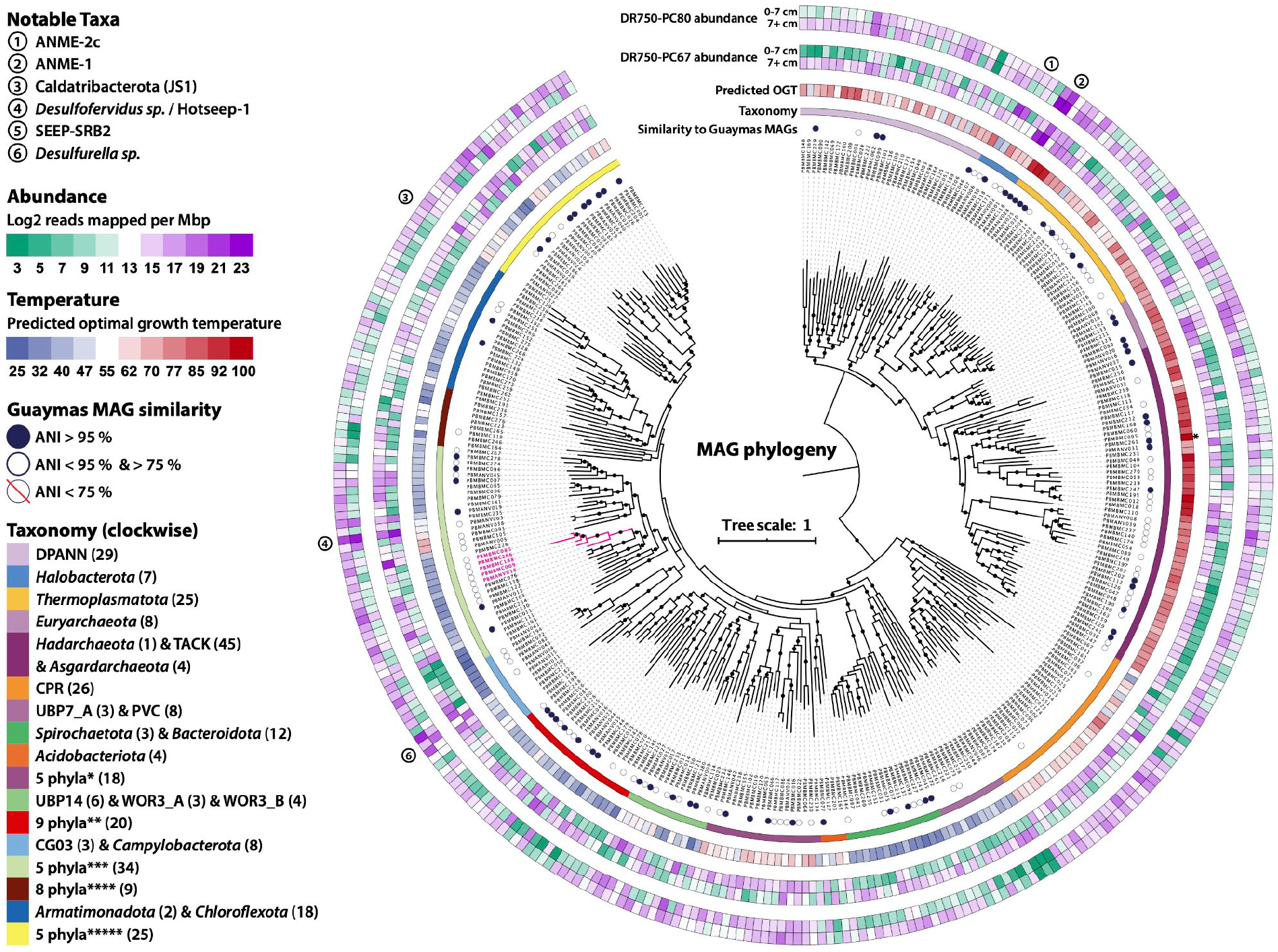
Concatenated marker gene phylogeny of the 325 Auka MAGs. Phylogeny of the 325 MAGs recovered from Auka vent field sediments, based on 25 concatenated marker genes. The scale bar indicates 1 substitution per site. From inside to outside, the concentric circles around the phylogeny indicate: the MAG ID, the average nucleotide identity (ANI) with MAGs previously retrieved from Guaymas Basin, phylum level taxonomy, MAG predicted optimal growth temperature (OGT), MAG abundance in the surface (0-7cm) and deep (7+cm) section of core DR750-PC67, and MAG abundance in the the abundance in the surface (0-7cm) and deep (7+cm) section of core DR750-PC80. Numbers in parentheses indicate the number of MAGs belonging to that lineage in the dataset (All MAGs are shown in the figure). The Tharpobacteria branch (see text) is highlighted in pink. The predicted OGT of PBMBMC_261 (111 °C) was outside the scale, and is indicated with an asterisk. Phyla corresponding to abbreviated groups in the taxonomy legend: *1 UBP7 (1), Ratteibacteria (2), Omnitrophota (6), Calescibacterota (2), Aerophobota (7); **Fermentibacterota (1), Krumholzibacteriota (1), Cloacimonadota (4), Latescibacterota (3), Zixibacteria (2) KSB1 (1) SM23-31 (1), Calditrichota (1), Marinisomatota (6); ***Proteobacteria (2), Myxococcota (1), Desulfuromonadota (3), Desulfobacterota_A (5), Desulfobacterota (23); ****UBP3 (1), Sumerlaeota (1), RBG-13-66-14 (1), Poribacteria (1), Hydrogenedentota (1), Eremiobacterota (1), Firmicutes (1), Firmicutes_A (2); *****Caldatribacteriota (2), Synergistota (3), Caldisericota (3), Bipolaricaulota (6), Thermotogota (11).

15 MAGs were more than twofold depleted in the <10 μm filtered fraction (Supplemental Data S5). The most depleted MAG in this fraction was a dominant ANME-2c archaeon, which is consistent with members of this methanotrophic clade frequently forming multi-celled consortia >10 μm in association with syntrophic sulfate-reducing bacteria (SRB). Notably, the putative sulfate-reducing partner of ANME-2c (e.g. *Desulfobacterota* SEEP-SRB2), was also depleted (1.88 fold) in the filtered fraction. Another clade depleted in the filtered fraction were the heterotrophic bacteria *Izimaplasma* originally described from methane cold seeps (Skennerton et al. 2016; Zheng et al. 2021). While anaerobic enrichments of this clade consisted of free-living organisms (Skennerton et al. 2016), their depletion in the filtered fraction could be due to association with larger organic particles for heterotrophic growth on DNA (Zheng et al. 2021). Notably, all four recovered MAGs belonging to the *Odinarchaeota* clade of Asgard Archaea were also strongly depleted in the filtered fraction. The morphology and eco-physiology of *Odinarchaeota* is poorly characterized and these findings may suggest an association with sediment particles or other microorganisms, or perhaps linked to an increased effective cell size due to unusual morphology, as observed for *Prometheoarchaeum* MK-D1, the only cultured member of the Asgard Archaea (Imachi et al. 2020). None of the four ANME-1 or three *Desulfofervidales* MAGs were strongly depleted in the filtered sample, consistent with previous observations that sediment-hosted ANME-1 often occur as single cells and tend to form less well-structured aggregates with SRB relative to ANME-2 consortia (Orphan et al. 2002; Holler et al. 2011).

Conversely, 14 MAGs were >2 fold enriched in the filtered fraction, suggesting limited association with the sediment matrix relative to other community members. These 14 MAGs represented 12 bacteria (seven Proteobacteria, two Synergistota, one Desulfobacterota, one Poribacteria, and one Thermotogota), and two Thermoplasmatota Archaea (Supplemental Data S5). Six of these MAGs (three Alphaproteobacteria and three Gammaproteobacteria) were absent from the unfiltered data and were therefore considered contamination and removed from further analyses, leaving 325 MAGs representing 54 phyla (Figure 2).

The 325 MAGs recovered from 2 sediment cores describe the microbial community at Auka vent field at a single location. To better understand the sediment microbial community structure and distribution patterns in the total vent area in relation to the physicochemical parameters, we conducted an area-wide survey using 16S rRNA gene amplicon sequencing of 29 additional sediment push cores, representing 227 total samples, collected by remotely operated vehicle during expeditions in 2017 and 2018 (Figure 1). Porewater geochemical profiles for 22 of these cores showed a variable degree of mixing of hydrothermal fluids with seawater, as evidenced by a steep drop in magnesium concentration with depth (Von Damm et al. 1985), and concurrent increases in calcium and potassium concentration (Supplemental figures S2 and S3, Supplemental Data S1). Consistent with a variable degree of mixing, temperature measured at 30 cm depth in the sediments varied greatly, ranging from 10°C to 177°C, a range similar to reports from Guaymas basin sediments (Biddle et al. 2012; McKay et al. 2016; Dowell et al. 2016; Teske et al. 2016). Concentrations of oxidized nitrogen species (nitrate and nitrite) were below the detection limit in porefluids, while ammonium reached concentrations as high as 16 mM, likely as a result of thermal degradation of organic matter (Haberstroh and Karl 1989). Porewater sulfide profiles frequently showed maxima up to 12 mM between 5 and 15 cm below the sediment surface. Sulfate concentrations dropped below the detection limit in the top 20 cm below the seabed in 14 cores, attributed to a combination of microbial sulfate reduction and seawater mixing with sulfate-depleted hydrothermal fluid. In cores with the highest inferred flux of hydrothermal fluid (e.g NA091-119 and S200-PC1), porewater sulfate concentrations were consistently below 10 mM at all depths (Supplemental figures S2 and S3).

**Figure 3.**
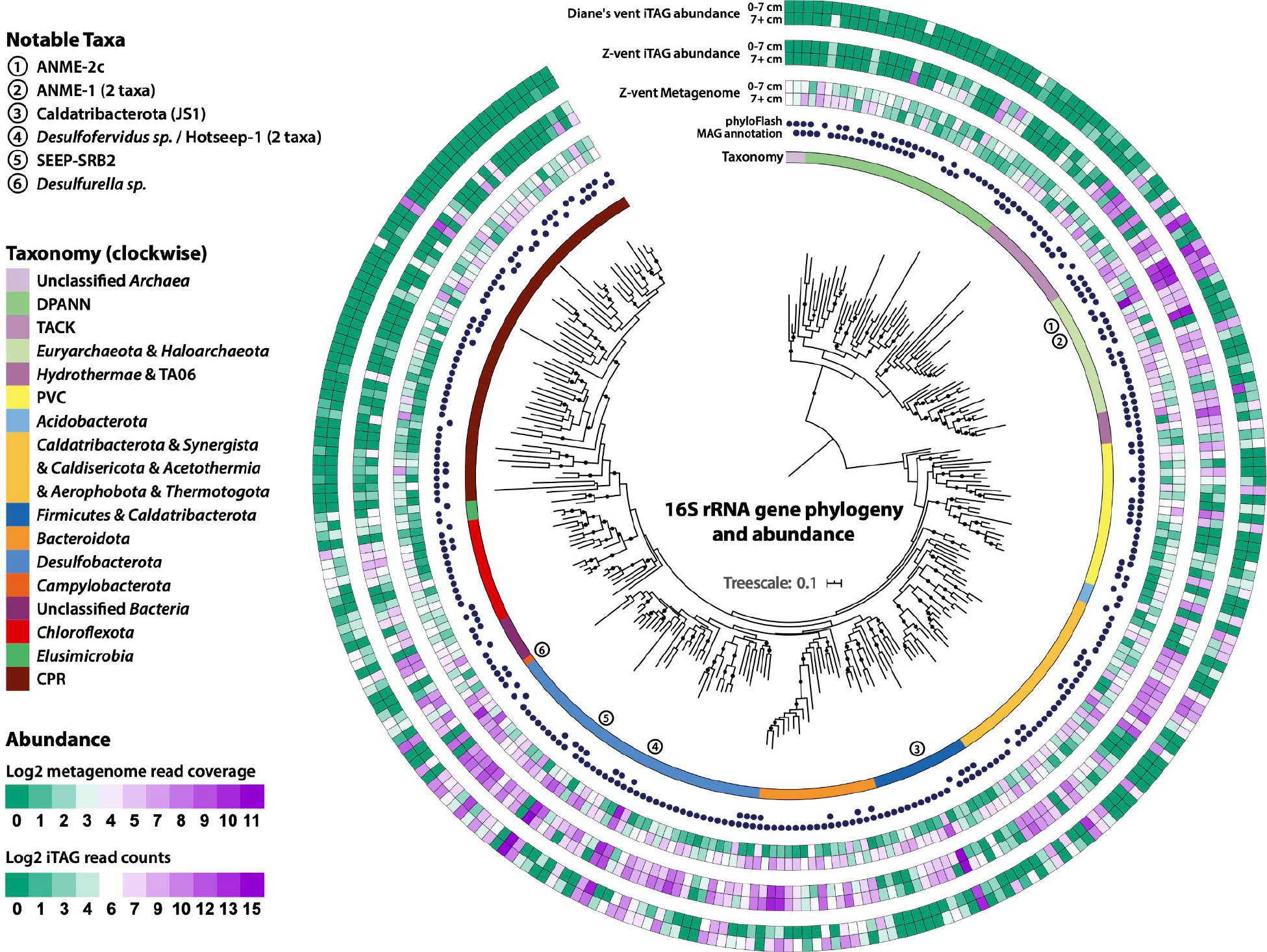
16 rRNA gene phylogeny and abundance in metagenome and amplicon data. Phylogeny of 284 16S rRNA genes reconstructed from the metagenomic data. From inside to outside the concentric circles around the phylogenetic tree indicate: the taxonomy of the major clades in the phylogeny, as assigned by Silva138, the recovery of the sequence through phyloFlash and/or annotation of the retrieved MAGs, the abundance in the surface (0-7cm) and deep (7+cm) section of the cores used for metagenomic sequencing, the abundance in the surface (0-7cm) and deep (7+cm) sections of the cores retrieved from Z-vent through amplicon sequencing, and the abundance in the shallow (0-7cm) and deep (7+cm) sections of the cores retrieved from Diane’s vent through amplicon sequencing. Circled numbers highlight abundant taxa discussed in the main text.

The taxa detected in our 16S rRNA gene amplicon survey were broadly similar in the 29 cores (Supplemental Figures S4 - S32). A comparison of the amplicon sequences with 16S rRNA genes retrieved from the metagenome using PhyloFlash (Gruber-Vodicka, Seah, and Pruesse 2020), and assembled 16S rRNA genes from select MAGs, showed congruence between the most abundant taxa in all 29 cores and dominant taxa recovered from metagenome sequencing (Figure 3). This shows that the MAGs assembled from a single location (2 adjacent cores) provides representative insight of the microbial community within the greater vent field. The overall distribution pattern in Auka sediments indicates widespread distribution of sulfide/sulfur-oxidizing Campylobacterota (formerly Epsilonproteobacteria) and putatively heterotrophic Bacteroidota as part of the surface microbial mat assemblage, with limited recovery of sulfur-oxidizing *Beggiatoa* (Supplemental text). Within the underlying sediment, ANME-2c archaea and Seep-SRB2 sulfate-reducing bacteria (SRB) of the *Dissulfuribacteraceae,* linked to the sulfate-dependent anaerobic oxidation of methane were dominant near the sediment surface, with consortia of ANME-1 Archaea and *Desulfofervidus sp.* SRB were highly abundant in deeper sediment horizons. This ANME niche separation along temperature and geochemical gradients has been reported previously from Guaymas Basin, with ANME-2c observed in lower temperature cores, and ANME-1 in higher temperature cores (Teske et al. 2002; Biddle et al. 2012; Holler et al. 2011). Besides temperature, sulfate concentration may also contribute to niche separation at Auka, with ANME-1 phylotypes dominant in horizons corresponding to low sulfate concentrations, a pattern that has also been described from cold seeps. In several cores (e.g. NA091-118 & 119) amplicon sequence variant (ASV) distribution showed further depth stratification of ANME-1 phylotypes, with ANME-1a ASVs co-occurring with Seep-SRB2 phylotypes in shallower horizons, again consistent with observations from methane cold seeps (Metcalfe et al. 2021) (Supplemental figures S10 & S11). Other clades present in high abundance are *Desulfobacteraceae* in the shallow sediments, with Caldatribacterota (JS1), Thermoplasmatota (DHVEG-2), and Thermotogota ASV’s common in the deeper horizons (Figure 3). Consistent with the variation in porewater ion concentrations from hydrothermal fluid advection, the depth distribution of taxa varied between sediment cores, with higher inferred hydrothermal fluid input corresponding to limited detection of ANME-2c, higher abundance of ANME-1 in the shallow horizons, and appearance of hyperthermophilic lineages (e.g. Methanopyraceae, Thermoprotei) in the deeper horizons. A full summary of sediment 16S rRNA diversity from the 29 core survey is provided in Supplemental Figures S4-S32 and Supplemental Data S2-S4.

Both the 16S rRNA diversity analyses and the metagenomic sequencing indicate community stratification by depth in the sediment. We expect temperature to play a major role in shaping community structure, because of the high degree of hydrothermal fluid mixing and observed steep thermal gradients (up to 7°C cm^-1^). While broad trends in relative abundance of sediment-hosted microbial community members along a temperature gradient have been shown for Guaymas Basin and other hydrothermal sites (Lutz et al. 2008; Anderson et al. 2013; Ding et al. 2017), the relationship of these trends to optimal growth temperature (OGT) is lacking for the majority of taxa due to the lack of cultured representatives. Genomic data is shedding new light on physiological characteristics like OGT, with strong correlations observed between select genomic features and OGT for cultured bacteria and archaea ((Sauer and Wang 2019) and references therein). This knowledge was recently synthesized and incorporated into an OGT prediction model and used to accurately predict OGTs of phylogenetically diverse cultured microorganisms (Sauer and Wang 2019). Here we apply a modified version of this model (see methods) and used this to estimate OGT for the environmental MAGs recovered from the Auka vents, Guaymas Basin, and a selected set of over 4000 archaeal and bacterial genomes from the GTDB (Figure 2, Supplemental Data S5). For several microorganisms with known OGT or enrichment temperature, these genome-based predictions were largely consistent with reported values. For example, the predicted OGTs of the three Auka *Desulfofervidales* MAGs were 52°C, 53°C, and 60°C, close to the experimentally determined OGT of 60°C for *Desulfofervidus auxili* in pure culture (Krukenberg et al. 2016) and the predicted OGT of the two ANME-1 MAGs in the GB60 clade were 63°C and 65°C, also close to the 60°C enrichment temperature of the GB60 enrichment culture (Holler et al. 2011) (Figure 2, Supplemental Data S5). Predicted OGT of the MAGs correlated with their abundance (approximated by read coverage) as a function of sediment depth, with most MAG’s with OGTs over 50°C showing higher abundance in the >7 cm horizon in both cores. Notably, this OGT depth trend was less pronounced in core DR750-PC80, collected directly adjacent to localized hydrothermal fluid discharge, compared to core DR750-PC67 which was collected on the far side of DR750-PC80, with the closest edge approximately 10 cm further away from the fluid source (Figure 4, Supplemental Figure 1). While sediment temperatures were not measured for these cores, the visible fluid discharge at the seabed indicates that core DR750-PC80 likely was exposed to a greater flux of hydrothermal fluids and steeper temperature gradients relative to PC67, potentially explaining the higher abundance of MAGs with predicted OGT values over 50°C in the 0-7 cm horizon.

**Figure 4.**
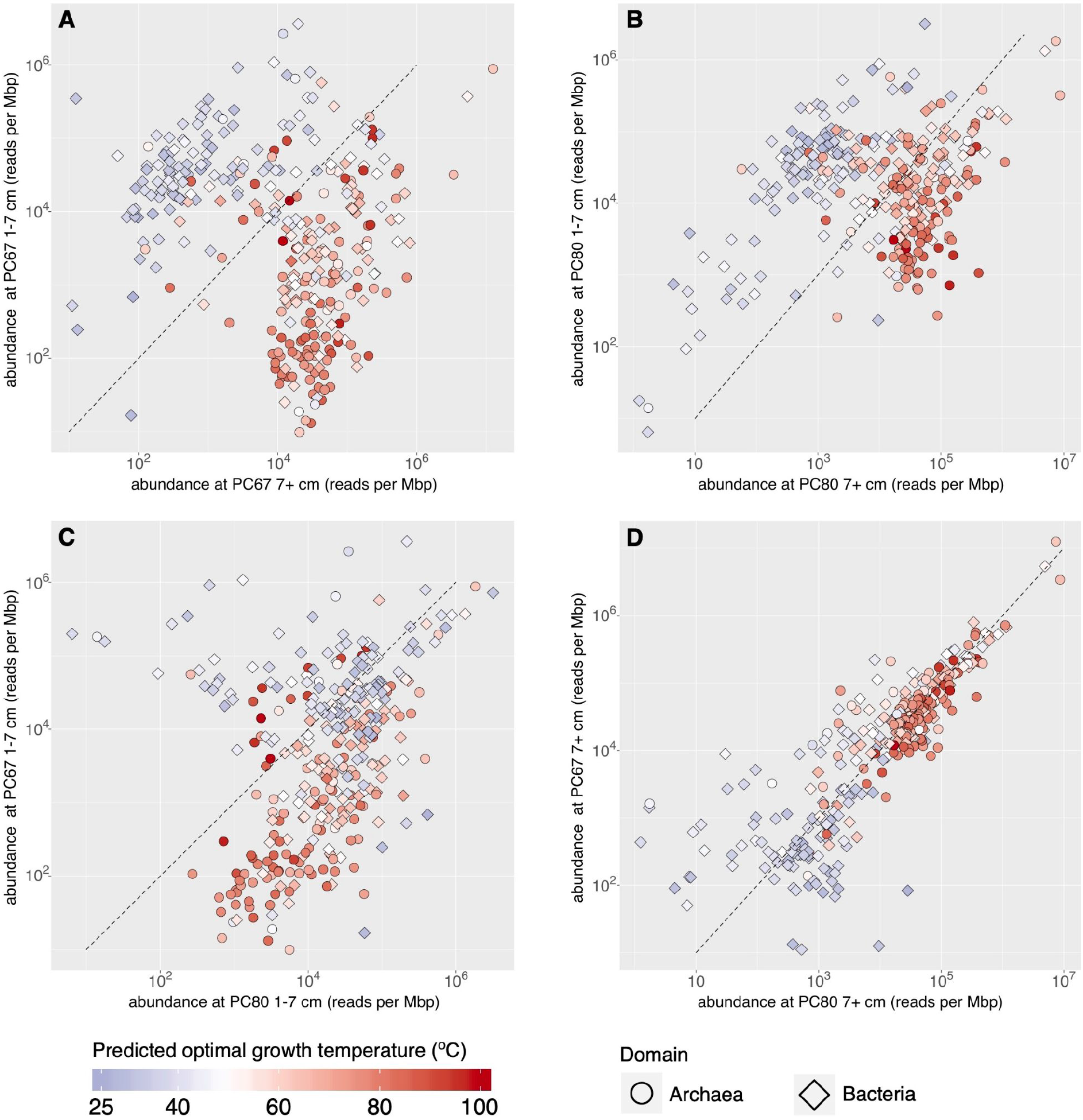
Abundance and predicted optimal growth temperature of the 325 Auka MAGs. Scatter plots showing reads per million base pairs of the Auka MAGs, as a proxy for organism abundance, in the sampled cores with point color corresponding to predicted optimal growth temperature (OGT) of each MAG. The dashed line represents equal abundance between both samples. **A)** Comparing MAG abundance between the DR750-PC67 surface horizon (0-7 cm) and DR750-PC67 deep horizon (7+ cm). **B)** Comparing MAG abundance between the DR750-PC80 surface horizon (0-7 cm) and DR750-PC80 deep horizon (7+ cm). **C)** Comparing MAG abundance between the surface horizons of both cores. **D)** Comparing MAG abundance between the deep horizons of both cores.

In addition to the correlation between predicted OGT and increasing abundance in lower horizons, there also is a clear correlation between predicted OGT and phylogeny (Figure 2). Using our MAGs and reference data from the genome taxonomy database (GTDB) (Parks et al. 2020), we assessed the phylogenetic signal of predicted OGT in more detail. After taxonomic assignment (Chaumeil et al. 2019), Auka MAGs, recent Guaymas MAGs (Dombrowski et al. 2017; Dombrowski, Teske, and Baker 2018; Seitz et al. 2019), and reference genomes from the GTDB (v89) were used to construct clade specific phylogenies with corresponding OGT predictions, representing a total of 5111 genomes, including all archaea and the majority of bacterial phyla included in GTDB v89. Within these 14 phylogenies, organisms with predicted (hyper)thermophilic OGT values grouped together in distinct lineages, often emerging from lineages with predicted mesophilic OGT values, suggesting thermophilic adaptation is common throughout both the bacterial and archaeal domains of the tree of life, and frequently persists in a lineage once acquired (Supplemental Figures S33 - S46). In addition, there are several examples of mesophilic lineages evolving from hyperthermophilic ancestors, such as the Nitrososphaeria (formerly Thaumarchaea), showing that thermophily is a reversible trait (Supplemental figure S33). The OGTs predicted for MAGs retrieved from Auka and for previously published MAGs from Guaymas Basin indicate the occurrence of hyperthermophilic microorganisms across many phyla at these sites, with one MAG from an uncultured member of the Thermoprotei from Guaymas Basin having the highest predicted OGT of all organisms analyzed, at 117°C. Among the Auka MAGs, a member from the same Thermoprotei clade had a predicted OGT of 111°C (Figure 3, Supplemental Figure S33, Supplemental Data S5). Surprisingly, the four Asgard Archaea MAGs recovered from Auka are the only predicted (hyper)thermophiles in this clade, despite several other Asgard MAGs being retrieved from hydrothermal sites (Supplemental Figure S33). Consistent with surveys of cultured organisms (Sauer and Wang 2019), the maximum predicted OGT in the 5111 genomes is considerably higher for Archaea (117°C) than for Bacteria (89°C);(Figure 2), although we note that not all Bacteria in the GTDB were included in this analysis.

In addition to clade-specific adaptation to high temperature, the phylogenetic analysis showed that Guaymas Basin and Auka vent fields have substantial community overlap. To further quantify this overlap in microbial communities between sites, we calculated average nucleotide identity (ANI) between all MAGs for both the Auka and Guaymas datasets. This analysis revealed that 68 Auka MAGs, representing 23 bacterial and archaeal phyla, have ANI values that are >95% to MAGs from Guaymas Basin sediments (Dombrowski et al. 2017; Dombrowski, Teske, and Baker 2018; Seitz et al. 2019), corresponding to nearly 20% species overlap between these geographically distant vent fields in the Gulf of California (Jain et al. 2018; Olm et al. 2020) (Figure 2 & Figure 5A, Supplemental Data S5). Another 93 Auka MAGs had ANI values between 75% and 95% with MAGs from Guaymas, further showing broad community similarity. Previous work has shown that the community at Guaymas Basin is distinct from those at basalt-hosted and ultramafic systems (Reveillaud et al. 2016). Indeed, a comparison with the MAGs retrieved from sampling deep hydrothermal fluids at Juan de Fuca ridge in the north Pacific (Jungbluth, Amend, and Rappé 2017), and hydrothermal fluids from mafic and ultramafic vent sites Cayman rise in the Caribbean (Anderson et al. 2017) showed only 9 and 12 MAGs with ANI above the 75% threshold value to the MAGs from Auka respectively, and no MAGs with >85% ANI in either dataset (Figure 5A).

**Figure 5.**
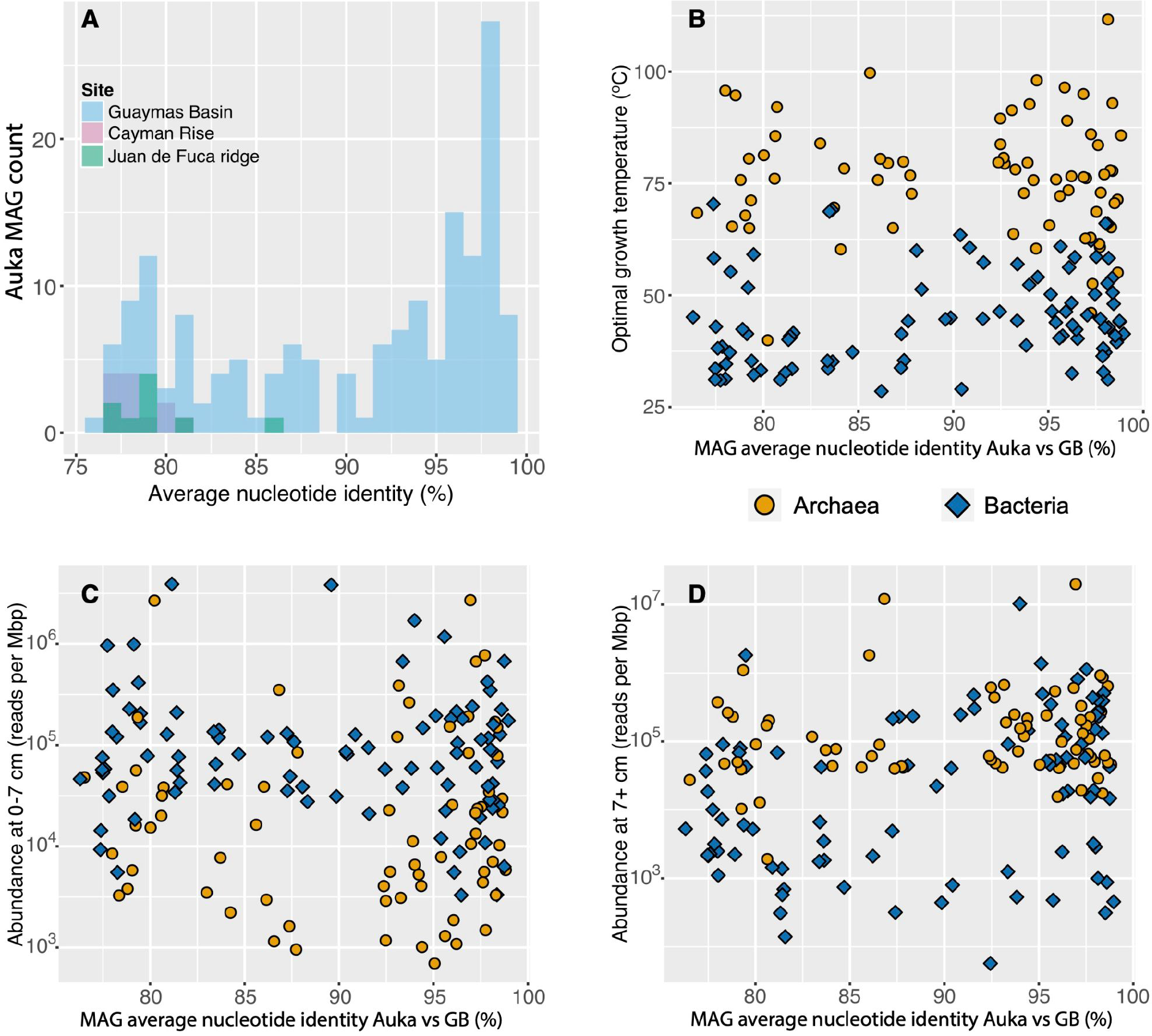
Community similarity between Pescadero Basin and other hydrothermal sites. **A)** Histograms of the Auka MAGs with average nucleotide identity (ANI) greater than 75% with MAGs obtained from Guaymas Basin (161), Cayman Rise (12), and Juan de Fuca ridge (9), indicating a high community similarity between Auka and Guaymas Basin. **B)** Scatterplot of Auka vs Guaymas Basin (GB) MAG ANI and predicted optimal growth temperature (OGT), showing no correlation between OGT and ANI. **C)** Scatterplot of Auka vs GB MAG ANI and MAG abundance in the surface horizons, indicating no correlation between surface abundance and ANI. D) Scatterplot of Auka vs GB MAG ANI and MAG abundance in the deep horizons, indicating no correlation between abundance in deeper horizons and ANI.

We hypothesize that the observed 20% species overlap is due to continuous transfer of microbial populations between Auka and Guaymas Basin. The exchange over the ∼400 km between both sites may be facilitated by hydrothermal plumes, which have been shown to rise at least 900 m upwards from the vents at Guaymas Basin (Merewether, Olsson, and Lonsdale 1985) and harbor distinct microbial communities from the surrounding seawater (Dick and Tebo 2010; Dick 2019; Anantharaman, Breier, and Dick 2016). In addition, long distance transfer of thermophilic microorganisms through ocean currents has been documented (Hubert et al. 2009; Müller et al. 2014; Resing et al. 2015), and the open ocean has been shown to contain a “seed bank” of hydrothermal vent taxa (Gonnella et al. 2016). Several organisms in this “seed bank” were also shown to be culturable from seawater particulates (Stetter et al. 1993), indicating that plume mediated microbial population transfer throughout the Gulf of California is highly likely.

Considering that hydrothermal plumes likely facilitate a constant flux of organisms between both basins, the species overlap of 20% indicates selection at dispersal, transfer, colonization, or a combination of these. As there is a strong correlation between temperature and community composition at Auka, we investigated whether the predicted OGT of MAGs correlated with ANI values between Auka and Guaymas Basin, but found no correlation, with both mesophilic and thermophilic groups represented among the 20% (Figure 5B). In addition, there was no correlation with ANI values and MAG abundance in surface sediments (Figure 5C), or MAG abundance in deeper horizons, with both abundant and rare taxa represented (Figure 5D). The latter observation contrasts previous work showing that abundant taxa were more likely to be cosmopolitan (Anderson, Sogin, and Baross 2015). The sediment MAGs with species-level overlap between the Gulf of California vent sites are phylogenetically and physiologically diverse, suggesting there is not a single determinant of transfer and colonization success.

Based on these results, we hypothesize that the species overlap between Auka and Guaymas is primarily driven by the niches created from similar environmental conditions in the hydrothermal vent sediments and deeply sourced fluids (Paduan et al. 2018), selecting for colonization by a subset of the transferred microbial population. In addition, the distribution of ANI values may indicate ongoing speciation between populations within the communities at the two sites. Previous studies have shown a gap in pairwise ANI values in the range 85% - 95% which was used to support 95% ANI as the species cutoff (Jain et al. 2018; Olm et al. 2020). While there is a clear peak of ANI values above 95% between Auka and Guaymas MAGs, an additional 25 MAGs (8% of the community) share ANI values between 90% - 95% showing a less pronounced gap in ANI values than previously observed (Figure 5A). These MAGs may represent lineages undergoing speciation after immigration and colonization of the new vent site.

Thus far, the genomic determinants for colonization success have not been established, potentially because such factors are likely to be lineage specific. For example, it is striking that strains of the same archaeal ANME-1 species are the most abundant organism in sediments from both the Auka vent field and the Guaymas basin, but that the other six ANME-1 MAGs (three at each site) belong to distinct lineages, suggesting niche differentiation (Supplemental Figure S36). 16S rRNA gene amplicon sequencing, which was done at higher sediment depth resolution, showed differential distribution of ANME-1 lineages in multiple sediment cores at Auka, supporting niche differentiation between them (Supplemental Figure S4 - S32).

Beyond implications for biogeography, the lineages largely or fully consisting of organisms found at Auka and Guaymas are of interest for future comparative genomics work. For example, several Aerophobota with OGT’s between 50°C and 68°C were detected in both Auka and Guaymas Basin (Supplemental fig. S38). MAGs from this phylum were recently also retrieved from other hydrocarbon-rich environments, including methane cold seeps in the South China Sea (Huang et al. 2019) and areas of petroleum seepage in the Gulf of Mexico (Dong et al. 2019), making this clade a promising target for genome guided metabolic predictions and enrichment cultivation focused on anaerobic hydrocarbon degradation. Another example is a deep branching clade within the Desulfobacterota phylum (Supplemental figure S43), likely representing a novel class, that was originally discovered in Guaymas Basin (Dombrowski, Teske, and Baker 2018) and is well represented in the surface layers of Auka vent field sediments. The Guaymas basin MAGs representing this group (indicated as DQWO01) were also included in a recent large scale analysis of Desulfobacterota metabolic potential (Langwig et al. 2021). We recovered five MAGs from this clade, ranging from 55 - 92% estimated completeness, with estimated contamination ranging from 0 - 4.3%. Combined with three MAGs from Guaymas Basin (completeness 55 - 70%, contamination 1.6 - 7.1%) these eight MAGs formed a deep branching monophyletic clade within the Desulfobacterota which we selected for further analysis. The size of the Desulfobacterota MAGs ranged from 1.3 - 3.1 Mbp and, using read mapping as a proxy for abundance, these organisms are enriched in the surface horizons of cores taken at both Auka and Guaymas. Their predicted OGT values (35 - 41°C) is consistent with a niche in a mesophilic environment. In accordance with the recent proposal to use genomic information as type material for naming microbial taxa (Chuvochina et al. 2019), we propose “*Candidatus Tharpobacterium aukensis”* for the organism represented by the PB_MBMC_085 MAG (82% estimated completeness, 0% estimated contamination), and will refer to the clade containing these eight genomes as Tharpobacteria (Supplemental Data S5). The genus name *Tharpobacterium* was chosen in honor of Marie Tharp, for her work on ocean floor mapping and plate tectonics that is key to our understanding of hydrothermal vents, where these organisms are found. The species name *aukensis* represents Auka, the location the MAG for which the name is proposed was recovered.

To investigate the metabolic potential of the Tharpobacteria, we performed functional enrichment analysis (Shaiber et al. 2020) on the 427 genomes within the Desulfobacterota obtained from Auka, Guaymas Basin and GTDB v89 (Supplemental Figure S43). This analysis indicated the Tharpobacteria MAGs encode the potential for beta oxidation of long chain fatty acids and degradation of benzoyl-CoA, indicating a possible role in aromatic hydrocarbon degradation (Boll and Fuchs 1995). Furthermore, Tharpobacteria are likely capable of degradation of butyrate, as recently reported for *Ca. Phosphitivorax sp.* in the UBA1062 order (Hao et al. 2020), another deep branching group within the Desulfobacterota. Unlike the UBA1062 genomes, Tharpobacteria MAGs encode NADH dehydrogenase (complex I), bc_1_-complex (complex III), and a heme biosynthesis pathway. We propose that Tharpobacteria MAGs use the electrons obtained from the oxidation of fatty acids and other hydrocarbons, and subsequently directed into the quinone pool via complex I to reduce a periplasmic electron acceptor (Figure 6A). However, the nature of this electron acceptor was not directly obvious from the genomic potential of the Tharpobacteria MAGs. Unlike many Desulfobacterota, the Tharpobacteria do not encode the capability to use sulfate as electron acceptor, and neither the functional enrichment analysis nor a follow up manual investigation of the genomes revealed terminal reductases for utilization of common electron acceptors.

**Figure 6.**
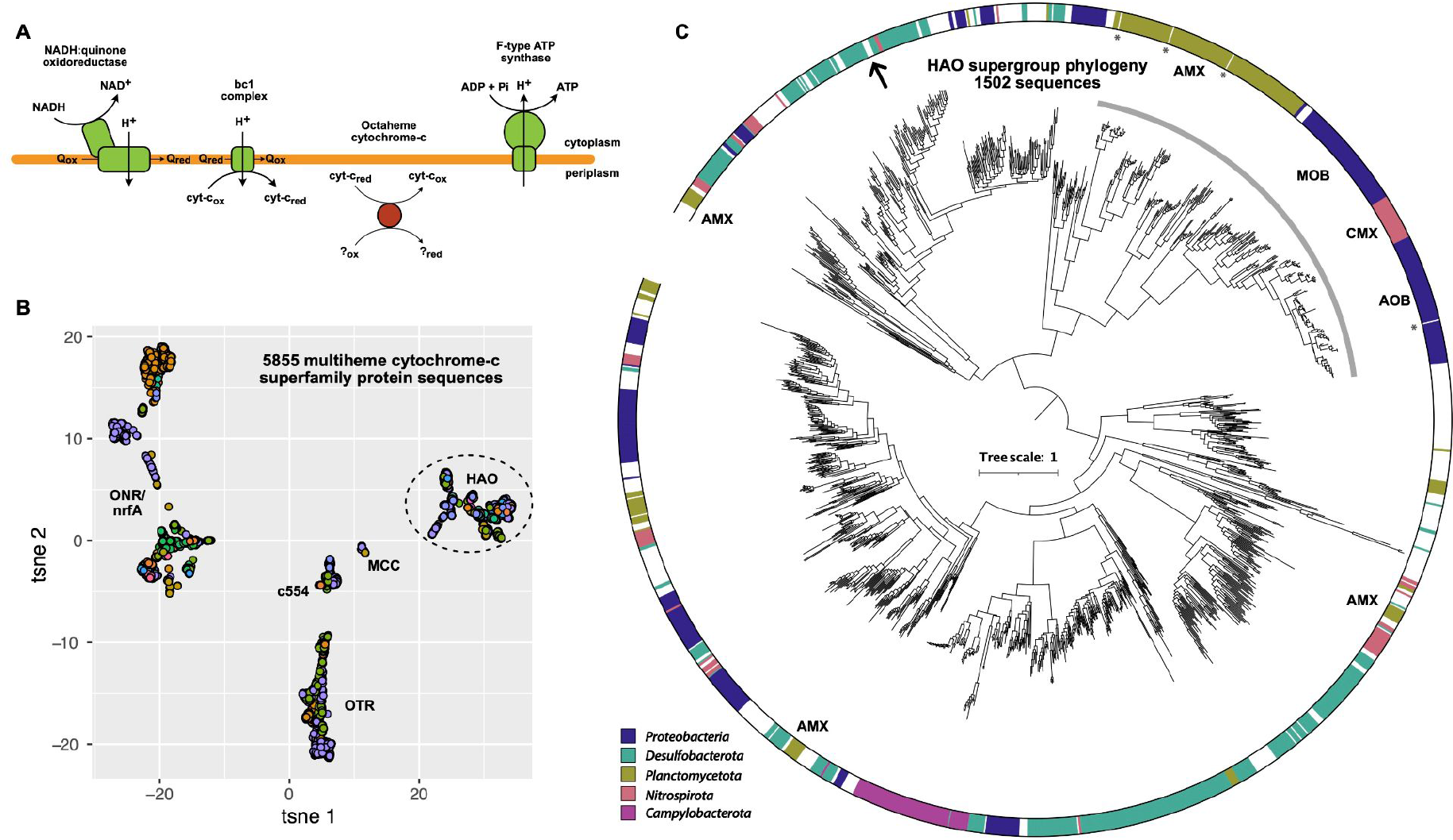
HAO family protein as proposed candidates for reduction of the terminal electron acceptor in Tharpobacteria clade. **A)** Schematic overview of components of the electron transport chain detected in Tharpobacteria genomes, with the octaheme cytochrome c proposed to be involved in terminal electron reduction in Tharpobacteria indicated in red. **B)** t-distributed stochastic neighborhood embedding (tSNE) representation of alignment score matrix of 5855 proteins of the HAO/OTR/ONR/nrfA/MCC/c554 structural fold family. Each point represents a protein sequence, colored by taxonomic affiliation at the phylum level. Dashed ellipse indicates sequences included in the phylogeny in panel C. HAO: hydroxylamine oxidoreductase, OTR: octaheme tetrathionate reductase, ONR: octaheme nitrite reductase, nrfA: pentaheme nitrite reductase, MCC: octaheme sulfite reductase, c554: tetraheme cytochrome c554. **C)** Approximate maximum likelihood phylogeny of 1502 HAO family protein sequences. The outer ring indicates taxonomic affiliation of the 5 most represented phyla, white space indicates other phyla. Sequences known to be involved in nitrogen cycling are indicated with a grey arc. Discussed Tharpobacteria clade sequences are indicated with an arrow, sequences of structures included in the promals3D alignment are indicated with asterisks, known clades involved in nitrogen cycling are highlighted with 3 letter abbreviations. The function of several HAO family members in anammox *Planctomycetota* is unknown. AOB: ammonia oxidizing bacteria, MOB: methane oxidizing bacteria, CMX: comammox *Nitrospirota*, AMX: anammox *Planctomycetota*.

We screened for other potential terminal reductases among the Tharpobacteria genomes by analyzing genes containing heme binding motifs (CxxCH), as hemes can be involved in both electron transfer and catalytic centers. A protein affiliated with the multiheme cytochrome-*c* (MCC) fold family, harboring well-characterized proteins involved in nitrogen and sulfur cycling (Simon et al. 2011), was present in 7 of the 8 MAGs. An exploratory analysis of all 5855 MCC family proteins retrieved from the GTDB genomes (v95) indicated this protein of interest was a member of the hydroxylamine oxidoreductase (HAO) family (Figure 6B). Proteins of the HAO family are known to play key roles in the nitrogen cycle, first identified as an essential protein in aerobic ammonia oxidizing bacteria (AOB) (Igarashi et al. 1997), and later shown to be essential for anaerobic ammonium oxidation (anammox) as well (Kartal and Keltjens 2016). HAO family proteins from Campylobacterota have been shown to catalyze nitrite reduction to ammonium *in vitro*, although their physiological role remains unclear (Haase et al. 2017). Notably, none of the HAO family proteins have suggested roles outside of the nitrogen cycle. However, our analysis found members of the HAO protein family in the genomes of many organisms not associated with nitrogen cycling, spanning 65 phyla (GTDB v95), and most commonly in the Desulfobacterota phylum. Phylogenetic analysis of the 1502 HAO family proteins showed that those with a known function in aerobic and anaerobic ammonium oxidation form a monophyletic clade. The Campylobacterota HAOs form a separate clade, while proteins of unknown function form several other clades within the family, with the Tharpobacteria proteins clustering within a clade with proteins from other Desulfobacterota (Figure 6C). Based on this distribution, and the absence of detectable oxidized nitrogen species from the Auka sediments, we hypothesize this protein catalyzes the reduction of sulfur species, rather than nitrogen. There is precedence for this as sulfur cycling has been shown for other members of the MCC fold family such as sulfite reductase mccA (Hermann et al. 2015), and octaheme tetrathionate reductase (Mowat et al. 2004). We propose that this HAO family protein catalyzes the reduction of the terminal electron acceptor in the Tharpobacteria clade, based on its prevalence across the MAGs.

It is also important to consider the possibility that individual Tharpobacteria MAGs use different electron acceptors, as there are several other protein complexes representing potential candidates as the site of the final reduction in the electron flow. Most prominently, several Tharpobacteria MAGs encode molybdopterin oxidoreductases, key complexes of many anaerobic metabolisms (Grimaldi et al. 2013). However, none of these molybdopterin oxidoreductase complexes have a known function, or are conserved in more than three of the eight genomes. In addition, there are several other cytochrome-containing proteins that could be candidate sites for terminal electron acceptor reduction in specific MAGs. Further research is needed to resolve the metabolism of this deep branching Desulfobacterota clade, but this metagenomic analysis offers an entry point for further characterization of the closely related HAO family proteins from isolated organisms as well as designing targeted enrichments, transcriptomic analyses, or stable isotope probing experiments using vent samples harboring Tharpobacteria.

## Discussion

Genome-resolved metagenomics is rapidly yielding genomic information of organisms across the tree of life, which is valuable for hypotheses about the ecology and physiology of organisms in uncultured lineages, and to provide targets for subsequent experimental study. Our analyses of the Auka sediments revealed a highly diverse microbial community, comprising many understudied lineages affiliated with sediment hosted hydrothermal vents. An in-depth look at one of these lineages, which we designated Tharpobacteria, revealed respiratory fatty acid degradation coupled to an unidentified electron acceptor, for which we propose reduction of a sulfur compound catalyzed by a MCC fold family protein. This provides a clear target for experimental verification, and confirmation of our hypothesis would have implications for interpretation of the biological sulfur cycle well beyond the Tharpobacteria.

In addition to gene function, investigating the genomic factors involved in colonization success at Auka and Guaymas is an exciting direction for future work, and a distinct advantage of using MAGs over the 16S rRNA gene amplicon analyses more frequently used for biogeography studies (Nemergut et al. 2011; Anderson, Sogin, and Baross 2015; Ruff et al. 2015). The geography and geological setting of the Gulf of California makes this region exceptionally well suited as a model system for hydrothermal vent biogeography. The narrow gulf constrains the currents and thus the direction of hydrothermal plumes and the increasing sediment thickness with distance from the mouth of the gulf differentiates conditions between the two known vent sites. The spreading centers on the Carmen and Farallon segments, located between the Guaymas and Pescadero segments (Lizarralde et al. 2007), could also harbor as yet undiscovered hydrothermal vents that act as stepping stones between the two sites.

Furthermore, the sediment-hosted hydrothermal vents of the Gulf of California provide an excellent study site for further discoveries of (hyper)thermophilic organisms. The maximum predicted OGT for organisms at Auka (111°C) and Guaymas (117°C) were substantially higher than the highest experimentally determined OGT values of 106°C for *Pyrolobus fumarii (Blöchl et al. 1997)*, and 105°C for *Methanopyrus kandleri* grown at high pressure (Takai et al. 2008). Interestingly, the predicted OGT for *M. kandleri* is 98°C, identical to its observed optimum at ambient pressure (Kurr et al. 1991), while the OGT prediction for *Pyrolobus fumarii* is 102°C, slightly lower than the reported OGT. Strain 121, with a predicted optimum of 103*°*C (Kashefi and Lovley 2003), does not have a publicly available genome sequence; but the predicted OGT of 98°C for *Pyrodictium abyssii,* the most closely related organism with a sequenced genome, is consistent with experimental observation (Pley et al. 1991). The MAGs with predicted OGTs higher than the highest experimentally observed OGTs are not restricted to a single lineage, but rather distributed over several clades in the Thermoproteota (formerly Crenarchaeota). The Thermoproteota are severely undersampled, with the MAGs from Auka and Guaymas combined representing 40% of the Thermoproteota genomes analysed in this study. This indicates further sampling will likely yield organisms with even higher predicted OGTs than those predicted here, and suggests the upper temperature limit of life could be considerably higher than currently known. Both *in situ* and *in vitro* experiments on the sediments of the Gulf of California hydrothermal vent sites could push our knowledge of the upper temperature boundary of life. Finally, OGT prediction and genome-resolved metagenomics provide a powerful combination to examine the evolutionary history of thermophily. Our analysis shows that thermophily has evolved frequently in both the Bacteria and Archaea (Supplemental figures S33-S46), suggesting adaptation of mesophiles to the colonization of high temperature environments. The rapidly increasing genomic representation of lineages in the tree of life makes phylogenomics combined with optimal growth temperature predictions an exciting angle on the debate about the conditions in which life on earth arose and diversified.

## Methods

### Sample collection & shipboard processing

Samples were collected from Auka vent field in Pescadero Basin (Gulf of California, Mexico, 23.954, -108.863) on R/V Western Flyer in 2015 (MBARI2015), E/V Nautilus in 2017 (NA091) and R/V Falkor in 2018 (FK181031); information for the required sampling permits are provided in the funding acknowledgement section. Pushcore samples were collected using remotely operated vehicle (ROV) Doc Ricketts (MBARI2015), ROV Hercules (NA091), and ROV SuBastian (FK181031) from sediment covered areas with microbial mat cover and/or visible hydrothermal fluid flow (Figure 1). Sites were deemed suitable for sampling if a 28 cm core could be fully inserted into the sediment. During MBARI2015, two sediment cores were collected ∼10cm and ∼15cm distance from a site of focused hydrothermal fluid discharge close to Z-vent (Figure 1, Supplementary Figure 1). Cores were split in surface (0-7 cm) and deep (7+ cm) horizons, and stored in heat sealed mylar at 4 °C, under nitrogen gas. During NA091, eight sediment cores were collected from locations across Auka vent field (Figure 1, Supplemental Figure 1). Cores were sectioned in 1-3 cm thick horizons (Supplemental Data S1 & S2). 2 mL subsamples of each horizon were frozen at -80 °C for DNA extraction, and 6 mL sediment was centrifuged at 16000g for 2 minutes to collect sediment porewater. 0.25 mL filtered porewater was preserved in 0.25 mL 0.5M zinc acetate solution for later sulfide analysis. 0.25 mL filtered porewater was stored at -20 °C for subsequent ion chromatography. During FK181031, 22 sediment cores were collected from locations across Auka vent field (Figure 1, Supplemental Figure 1). Cores were sectioned in 1 or 3 cm horizons (Supplemental Data S1 & S2). 2 mL subsamples of each horizon were frozen at -80 °C for DNA extraction, and porewater was extracted from ∼15 mL (1 cm horizons) or ∼50 mL (3 cm horizons) sediment under nitrogen gas using a pneumatic sediment squeezer (KC Denmark A/S, Silkeborg, Denmark). 0.25 mL filtered porewater was preserved in 0.25 mL 0.5M zinc acetate solution for sulfide analysis. 0.25 mL filtered porewater was stored at -20 °C for ion chromatography.

### Geochemistry

All filtered water samples were stored at -20°C until analysis. Major ions were measured with a Dionex ICS-2000 (Dionex, Sunnyvale, CA, USA) ion chromatography system (Environmental Analysis Center, Caltech) with anion and cation columns running in parallel. An autosampler loads samples diluted 1:50 in 18 MΩ water run through an LC-Pak polisher (MilliporeSigma, Burlington, MA, USA) serially to a 10 µL sample loop on the anions channel, then a 10 µL sample loop on the cations channel. Both columns and detectors are maintained at 30°C. The ion chromatography system was run as described previously (Green-Saxena et al. 2014) with the following modifications. Anions were resolved by a 2mm Dionex IonPac AS19 analytical column protected by a 2mm Dionex IonPac AG19 guard column (ThermoFisher, Waltham, MA, USA). A potassium hydroxide eluent generator cartridge generated a hydroxide gradient that was pumped at 0.25 mL/min. The gradient was constant at 10 mM for 5 minutes, increased linearly to 48.5 mM at 27 minutes, then increased linearly to 50 mM at 40 minutes. A Dionex AERS 500 suppressor provided suppressed conductivity detection running on recycle mode with an applied current of 30 mA. Cations were resolved by a 4mm Dionex IonPac CS16 analytical column protected by a 4mm Dionex IonPac CG16 guard column. A methanesulfonic acid eluent generator cartridge generated a methanesulfonic acid gradient that was pumped at 0.36 mL/min. The gradient was constant at 10 mM for 5 minutes, nonlinearly increased to 20 mM at 20 min (Chromeleon curve 7, concave up), and nonlinearly increased to 40 mM at 40 min (Chromeleon curve 1, concave down). A Dionex CERS 2mm suppressor provided suppressed conductivity detection with an applied current of 32 mA. Chromatographic peaks were integrated by Chromeleon 7.2 using the Cobra algorithm, and were correlated to concentration by running known standards. The threshold of detection was approximately 10 µM for bromide and thiosulfate, 50 µM for ammonium, 100 µM for calcium, potassium, and sulfate, and 400 µM for magnesium.

Sulfide was determined colorimetrically using a protocol based on (Cline 1969). Briefly, 22 μL of a 1:1 mixture of sample and 0.5 M zinc acetate was added to 198 μL milliQ water in a 96 well plate and mixed thoroughly. Immediately after mixing, 20 μL of the diluted sample was transferred to a second 96 well plate containing 180 μL milliQ water, resulting in one 96 well plate containing 10-fold diluted samples and one 96 well plate containing 100-fold diluted samples with 200 μL final volume in each well. 20 μL of 1:1 mixture of Cline reagents (30 mM FeCl_3_ · 6H_2_O in 6N HCl, and 11.5 mM N,N-dimethylphenylenediamine dihydrochloride (DPDD) in 6N HCl) was added to each well and allowed to react for 1hr before measuring on a Tecan Sunrise 4.2 plate reader, using the Magellan software (version 7.3). Duplicates of all samples were run on the same 96-well plate. Sample concentration was determined by comparison to a 12 point standard curve added to each plate in duplicate, and treated identically to the samples..

### DNA extraction and metagenome sequencing

The core horizons (0-7 cm and 7+ cm) from both cores obtained during MBARI2015, DR750-PC67 and DR750-PC80, were split in two subsamples each. Of these 8 samples, half was filtered through a 10 μm mesh to deplete sediment-attached organisms or those forming large aggregates, and the other half was untreated. DNA was extracted from 0.25 g wet sediment using a CTAB/phenol/chloroform organic solvent extraction protocol modified from Zhou et al. (Zhou, Bruns, and Tiedje 1996) as previously described (Speth et al. 2016). Cell lysis was achieved in a CTAB extraction buffer (10 g L^−1^ CTAB, 100 mM Tris, 100 mM EDTA, 100 mM sodium phosphate, 1.5 M NaCl pH 8) with serial addition of, and incubation with, lysozyme, proteinase K, and sodium dodecyl sulfate SDS. After cell lysis, DNA was recovered using phenol/chloroform/isoamylalcohol (25:24:1) at 65°C, followed by an additional chloroform/isoamylalcohol (24:1) extraction to remove traces of phenol. After recovery of the aqueous phase, DNA was precipitated using isopropanol at room temperature, and washed with ice-cold (-20 °C) 70 % ethanol and resuspended in nuclease-free water. Barcoded Nextera XT V2 libraries (Illumina) were made with dual sequencing indices, pooled, and purified with 0.75 volumes of AMpure beads (Agencourt). The resulting libraries were sequenced on a HiSeq2500 with 2x125 protocol in high output mode. (Elim Biopharmaceuticals, Hayward, CA, USA), resulting in 415 million paired reads.

### Assembly and Binning

To recover high quality metagenome assembled genomes (MAGs), an iterative assembly and binning workflow was adopted. Each single dataset was subsampled to 10 million reads, assembled using SPAdes v3.14.1 (Bankevich et al. 2012) and manually binned using Anvi’o (Murat Eren et al. 2015). Reads were mapped to the manual bins, using BWA (version 0.7.12-r1039) (H. Li and Durbin 2009), filtered at >95 % identity over >80 % of the read length using BamM, (http://ecogenomics.github.io/BamM/) with matching reads removed from the dataset, and the process was repeated for all datasets until no more manual bins could be obtained, resulting in 50 manual bins representing abundant community members (Supplemental Data S5). Subsequently, all unmapped reads of all eight datasets, 194 million paired reads total, were pooled, co-assembled using Megahit (version 1.2.9) (D. Li et al. 2015), and binned using Metabat2 (version 2.12.2) (Kang et al. 2019), and then manually inspected and corrected using Anvi’o (version 6.2) (Murat Eren et al. 2015). The automated correction removed >1 % of bases in 166 metabat bins and >20 % of bases from 54 metabat bins, highlighting the value of manual inspection. Bin quality was assessed using checkM (version 1.1.2), and the single copy marker gene sets included in Anvi’o. Bins with >50 % completeness and <10% redundancy as determined by CheckM were retained. 8 additional bins with >10% redundancy were included after a second manual inspection (Supplemental Data S5). This workflow resulted in 331 metagenome assembled genomes (MAGs), six of which were not detected in the unfiltered datasets. Those six MAGs were considered contamination and removed from all subsequent analyses. Taxonomic assignment of the MAGs was done using GTDB-tk (version 1.3.0) (Chaumeil et al. 2019).

### MAG phylogeny and annotation

The two-domain phylogenetic tree of all 325 MAGs was generated using the single copy marker gene sets “Archaea_76” and “Bacteria_71’’ included in Anvi’o, using the 25 genes shared between these two single copy marker sets. Genes matching the 25 selected markers were extracted, aligned using muscle (version 3.8.1551) (Edgar 2004), and the alignment concatenated using “anvi-get-sequences-for-hmm-hits”. The resulting concatenated alignment was converted to PHYLIP format using the ElConcatenero.py script (https://github.com/ODiogoSilva/ElConcatenero), and a phylogeny was calculated using RAxML (v8.2.12) (Stamatakis 2014), with the PROTGAMMALG4X model (Le, Dang, and Gascuel 2012) and the autoMRE bootstopping criterion resulting in 100 bootstrap replicates (Pattengale et al. 2009).

Gene calling on the MAGs was done using Prodigal (version 2.6.3) (Hyatt et al. 2010), and predicted genes were annotated using the DIAMOND (Buchfink, Xie, and Huson 2015) against the NCBI-NR database, PFAM (El-Gebali et al. 2019), KEGG (Kanehisa and Goto 2000), COG (Tatusov et al. 2003), CDD (Marchler-Bauer et al. 2015), EGGNOG (Huerta-Cepas et al. 2019, 2017), and CATH (Sillitoe et al. 2019; Lewis et al. 2018) (Supplemental Data S6). KEGGDecoder (Graham, Heidelberg, and Tully 2018) was used to assess broad metabolic capabilities of the MAGs.

A clade we refer to as Tharpobacteria, comprising eight genomes (five obtained from Auka and three from Guaymas), was chosen for further analysis. The metabolic capabilities of this clade were compared to 419 other *Desulfobacterota* genomes (Supplemental figure S43) by functional enrichment analysis with anvi-compute-functional-enrichment using KEGG module assignments from anvi-estimate-metabolism (Anvi’o version 7). Multiheme cytochrome c fold family proteins were obtained from the GTDB using two iterations of sequence recruitment and filtering using a bit score ratio (Rasko, Myers, and Ravel 2005) for each of the five constituent protein families. The resulting sequence sets were merged and dereplicated, and the resulting 5855 proteins sequences were classified using ASM-clust with t-distributed stochastic neighborhood embedding (tSNE) perplexity value set to 500 (Speth and Orphan 2019).

### MAG reference phylogenies and optimal growth temperature prediction

Species overlap between the Auka vent field MAGs and other hydrothermal vent field MAGs was assessed using FastANI (version 1.3). 666 Guaymas basin MAGs (Dombrowski et al. 2017; Dombrowski, Teske, and Baker 2018; Seitz et al. 2019) and 99 MAGs obtained from Juan de Fuca ridge hydrothermal fluid (Jungbluth, Amend, and Rappé 2017) were downloaded from NCBI. 348 bins from the Cayman Rise hydrothermal vent field (Anderson et al. 2017) were retrieved as Anvi’o databases (version 2.1.0) from Figshare (https://figshare.com/projects/Mid-Cayman_Rise_Metagenome_Assembled_Genomes/20783).

Contig fasta files were exported from the Anvi’o databases and 131 bins over 50% completeness were used in ANI analysis.

Detailed phylogenetic placement of the Auka MAGs was obtained by downloading the genomes comprising the identified phyla, as well as sister phyla, from the genome taxonomy database (GTDB, version 89) (Parks et al. 2020). In addition 666 MAGs recently obtained from Guaymas basin were included in the reference set (Dombrowski et al. 2017; Dombrowski, Teske, and Baker 2018; Seitz et al. 2019). The genome set was split in 14 subsets, based on taxonomy and the GTDB reference tree, for alignment and tree calculation. Anvi’o databases were generated for all genomes, and the “Archaea_76” and “Bacteria_71” gene sets included in Anvi’o were used to generate concatenated alignments for Archaeal and Bacterial subsets, respectively, using muscle (version 3.8.1551) (Edgar 2004). Phylogenies were calculated using FastTree (version 2.1.7) (Price, Dehal, and Arkin 2010).

Optimal growth temperatures (OGT) were predicted for the 325 MAGs and the reference genomes using the method described by Sauer and Wang (Sauer and Wang 2019) (https://github.com/DavidBSauer/OGT_prediction). Regression models modified by David Sauer to exclude 16S rRNA and genome size, to account for absence of 16S rRNA and genome incompleteness, were downloaded from https://github.com/DavidBSauer/OGT_prediction/tree/master/data/calculations/prediction/regres sion_models. The regression models for “Superkingdom Archaea” and “Superkingdom Bacteria” were chosen because of the diversity of the organisms in the dataset. The prediction_pipeline.py script was run as described in the documentation, with a custom “genomes_retrieved.txt” file (two tab-delimited columns, no header, columns: filename of gzip compressed contig fasta <tab> genome ID) and “species_taxonomic.txt” file (two tab-delimited columns, headers: “species <tab> superkingdom”, columns: “Genome ID <tab> Archaea|Bacteria”).

### 16S rRNA gene analyses

To gain insight in the microbial diversity of the Auka vent field, 216 samples derived from 30 sediment cores, and 8 samples from microbial mats or biofilms, were used for DNA extraction using the Qiagen Dneasy PowerSoil kit (Valencia, CA, USA) following the manufacturer’s protocol, with the exception that cells were lysed using MP Biomedicals FastPrep-24 (Irvine, CA, USA) for 45s, at 5.5 m s^-1^. The V4-V5 region of the 16S rRNA gene was PCR amplified from the resulting 224 DNA extracts using the 515f/926r primer set (Walters et al. 2016) modified with Illumina (San Diego, CA, USA) adapters on 5’ end (515F 5’-TCGTCGGCAGCGTCAGATGTGTATAAGAGACAG-GTGYCAGCMGCCGCGGTAA-3’and 926R 5’-GTCTCGTGGGCTCGGAGATGTGTATAAGAGACAG-CCGYCAATTYMTTTRAGTTT-3’).

Duplicate PCR reactions were set up for each sample with Q5 Hot Start High-Fidelity 2x Master Mix (New England Biolabs, Ipswich, MA, USA) in a 15 μL reaction volume, with annealing at 54°C, for 28 cycles. The number of cycles was increased if no product was obtained, as detailed in the sample metadata (Supplemental Data S2). Duplicate PCR samples were then pooled and barcoded with Illumina Nextera XT index 2 primers that include unique 8-bp barcodes (P5 5’-AATGATACGGCGACCACCGAGATCTACAC-XXXXXXXX-TCGTCGGCAGCGTC-3’ and P7 5’-CAAGCAGAAGACGGCATACGAGAT-XXXXXXXX-GTCTCGTGGGCTCGG-3’). Amplification with barcoded primers used Q5 Hot Start PCR mixture but used 2.5 μL of product in 25 μL of total reaction volume, annealed at 66°C, and cycled 10 times. Products were purified using Millipore-Sigma (St. Louis, MO, USA) MultiScreen Plate MSNU03010 with vacuum manifold and quantified using ThermoFisher Scientific (Waltham, MA, USA) QuantIT PicoGreen dsDNA Assay Kit P11496 on the BioRad CFX96 Touch Real-Time PCR Detection System. Barcoded samples were combined in equimolar amounts into single tubes and purified with Qiagen PCR Purification Kit 28104 before sequencing on Illumina MiSeq with the addition of 15-20% PhiX and with either a 2x250 or a 2x300 protocol (Laragen Inc., Culver City, CA, USA). Demultiplexed sequencing data was processed using QIIME2 (v2020.2) (Bolyen et al. 2019), with DADA2 for amplicon sequence variant (ASV) calling (Callahan et al. 2016) and Cutadapt for sequence quality trimming and primer removal (Martin 2011). resulted in 18777 ASVs representing 6,244,164 trimmed and filtered reads.

16S rRNA genes were recovered from the MAGs using the HMMs included in Anvi’o (Murat Eren et al. 2015), and by *de novo* assembly of the combined metagenomic sequencing reads using phyloFlash (Gruber-Vodicka, Seah, and Pruesse 2020). After combining and dereplication of the two sets of sequences at 97% identity using usearch (v11.0.667) (Edgar 2010), the resulting 284 sequences were aligned using muscle (v3.8.1551) (Edgar 2004), and a phylogenetic tree was calculated using RAxML (v8.2.12) (Stamatakis 2014), with the GTRGAMMA model and the autoMRE bootstopping criterion resulting in 400 bootstrap replicates (Pattengale et al. 2009). Metagenome abundance of the 284 dereplicated 16S rRNA gene sequences was determined by mapping the combined reads of the shallow (0-7cm) and deep (7+ cm) horizons of both cores onto the 16S sequences using bbmap, with a 95% identity filter (Bushnell 2014). Approximately 0.06% of the reads mapped on the genes, which is consistent with expectation based on the 16S rRNA gene, and average genome length. Abundance of the 284 dereplicated 16S rRNA gene sequences in the amplicon sequencing (ITag) data was determined by aligning all 18,777 ASVs to the 16S rRNA genes using BLAST version 2.10.1 (Altschul et al. 1990), and summing the number of reads represented by all ASVs with >97% identity to the assembled 16S rRNA genes.

## Acknowledgments

We thank the pilots, crew, and participants on the cruises to the southern Gulf of California: R/V Western Flyer operated by the Monterey Bay Aquarium Research Institute (MBARI), E/V Nautlius operated by the Ocean Exploration Trust, with cruise NA091 supported by the Dalio Foundation and Woods Hole Oceanographic Institute, and R/V Falkor operated by the Schmidt Ocean Institute. We appreciate the support and opportunity to sail with chief scientists Scott Wankel and Anna Michel on NA091. We also thank David W. Caress and Jennifer B. Paduan (MBARI) for providing the high resolution version of the bathymetric map in Figure 1, David Sauer for recalculating the optimal growth temperature prediction model to be appropriate for metagenomics, and Haley Sapers for providing a framework for the 16S rRNA gene amplicon processing. This research used samples provided by the Ocean Exploration Trust’s Nautilus Exploration Program, cruise NA091.

## Data availability

Raw metagenome reads, assembled metagenome bins, and 16S rRNA gene amplicon sequencing data are available in GenBank under BioProject accession number

## Funding

This work was supported by the Center for Dark Energy Biosphere investigations (C-DEBI), Canadian Institute for Advanced Science (CIFAR), the US Department of Energy, Office of Science, Office of Biological and Environmental Research under award number DE-SC0016469 to Victoria J. Orphan. Daan R. Speth was supported by the Netherlands Organisation for Scientific Research, Rubicon award 019.153LW.039. Feiqiao B. Yu and Stephen R. Quake were supported by the John Templeton Foundation grant 51250 and the Chan Zuckerberg Biohub.Victoria J. Orphan is a CIFAR fellow in the Earth 4D program.

Sample collection permits were granted by la Dirección General de Ordenamiento Pesquero y Acuícola, Comisión Nacional de Acuacultura y Pesca (CONAPESCA: Permiso de Pesca de Fomento No. PPFE/DGOPA-200/18) and l a Dirección G eneral de G eografía y Medio A mbiente, Instituto Nacional de Estadística y Geografía (INEGI: Autorización EG0122018), with t he associated Diplomatic Note number 18-2083 (CTC/07345/18) f rom l a Secretaría d e R elaciones Exteriores - Agencia Mexicana de Cooperación Internacional para el Desarrollo / Dirección General de Cooperación Técnica y Científica. Sample collection permit f or cruise NA091 was obtained b y t he O cean E xploration T rust under permit number EG0072017.

