## Supplemental figures S1-S46 and supplemental text for "Microbial community of recently discovered Auka vent field sheds light on vent biogeography and evolutionary history of thermophily"

**Supplemental information**

This file contains supplemental information to “Microbial community of recently discovered Auka vent field sheds light on vent biogeography and evolutionary history of thermophily” by Speth *et al.* It contains a list of supplemental data files, a list of supplemental figures, and Supplemental Text.

**List of Supplemental data files:**

Supplemental Data S1 - Processed major ion data for the Auka sediment cores
Supplemental Data S2 - Metadata for the 16S rRNA gene amplicon sequencing
Supplemental Data S3 - ASV abundance data in Auka sediment cores
Supplemental Data S4 - Fasta file with sequences for the 18777 ASVs
Supplemental Data S5 - Table with MAG metadata
Supplemental Data S6 - Annotations for all proteins in all MAGs
Supplemental Data S7 - Fasta file with all proteins in all MAGs


**List of Supplemental Figures:**

Supplemental Figure S1 - Environmental context of sediment pushcore samples
Supplemental Figure S2 - Major ions of eight sediment cores retrieved on NA091
Supplemental Figure S3 - Major ions of fourteen sediment cores retrieved on FK181031
Supplemental Figure S4 - ASV abundance by depth in core020
Supplemental Figure S5 - ASV abundance by depth in core048
Supplemental Figure S6 - ASV abundance by depth in core051
Supplemental Figure S7 - ASV abundance by depth in core087
Supplemental Figure S8 - ASV abundance by depth in core089
Supplemental Figure S9 - ASV abundance by depth in core111
Supplemental Figure S10 - ASV abundance by depth in core118
Supplemental Figure S11 - ASV abundance by depth in core119
Supplemental Figure S12 - ASV abundance by depth in S0193 PC1
Supplemental Figure S13 - ASV abundance by depth in S0193 PC2
Supplemental Figure S14 - ASV abundance by depth in S0193 PC3
Supplemental Figure S15 - ASV abundance by depth in S0193 PC5
Supplemental Figure S16 - ASV abundance by depth in S0193 PC7
Supplemental Figure S17 - ASV abundance by depth in S0194 PC0
Supplemental Figure S18 - ASV abundance by depth in S0194 PC1
Supplemental Figure S19 - ASV abundance by depth in S0194 PC2
Supplemental Figure S20 - ASV abundance by depth in S0194 PC3
Supplemental Figure S21 - ASV abundance by depth in S0194 PC4
Supplemental Figure S22 - ASV abundance by depth in S0196 PC1
Supplemental Figure S23 - ASV abundance by depth in S0196 PC5
Supplemental Figure S24 - ASV abundance by depth in S0196 PC6
Supplemental Figure S25 - ASV abundance by depth in S0196 PC7
Supplemental Figure S26 - ASV abundance by depth in S0196 PC8
Supplemental Figure S27 - ASV abundance by depth in S0198 PC1

Supplemental Figure S28 - ASV abundance by depth in S0198 PC3
Supplemental Figure S29 - ASV abundance by depth in S0198 PC5
Supplemental Figure S30 - ASV abundance by depth in S0200 PC1
Supplemental Figure S31 - ASV abundance by depth in S0200 PC5
Supplemental Figure S32 - ASV abundance by depth in S0200 PC7
Supplemental Figure S33 - Phylogeny and OGT prediction of Crenarchaeota
Supplemental Figure S34 - Phylogeny and OGT prediction of DPANN superphylum
Supplemental Figure S35 - Phylogeny and OGT prediction of Euryarchaeota
Supplemental Figure S35 - Phylogeny and OGT prediction of Euryarchaeota
Supplemental Figure S36 - Phylogeny and OGT prediction of Halobacterota
Supplemental Figure S37 - Phylogeny and OGT prediction of Thermoplasmatota
Supplemental Figure S38 - Phylogeny and OGT prediction of Acidobacteriota
Supplemental Figure S39 - Phylogeny and OGT prediction of phyla basal to Bacteroidota
Supplemental Figure S40 - Phylogeny and OGT prediction of Campylobacterota
Supplemental Figure S41 - Phylogeny and OGT prediction of Chloroflexota
Supplemental Figure S42 - Phylogeny and OGT prediction of CPR superphylum
Supplemental Figure S43 - Phylogeny and OGT prediction of Desulfobacterota
Supplemental Figure S44 - Phylogeny and OGT prediction of PVC superphylum
Supplemental Figure S45 - Phylogeny and OGT prediction of Spirochaetota
Supplemental Figure S46 - Phylogeny and OGT prediction of Thermoplasmatota

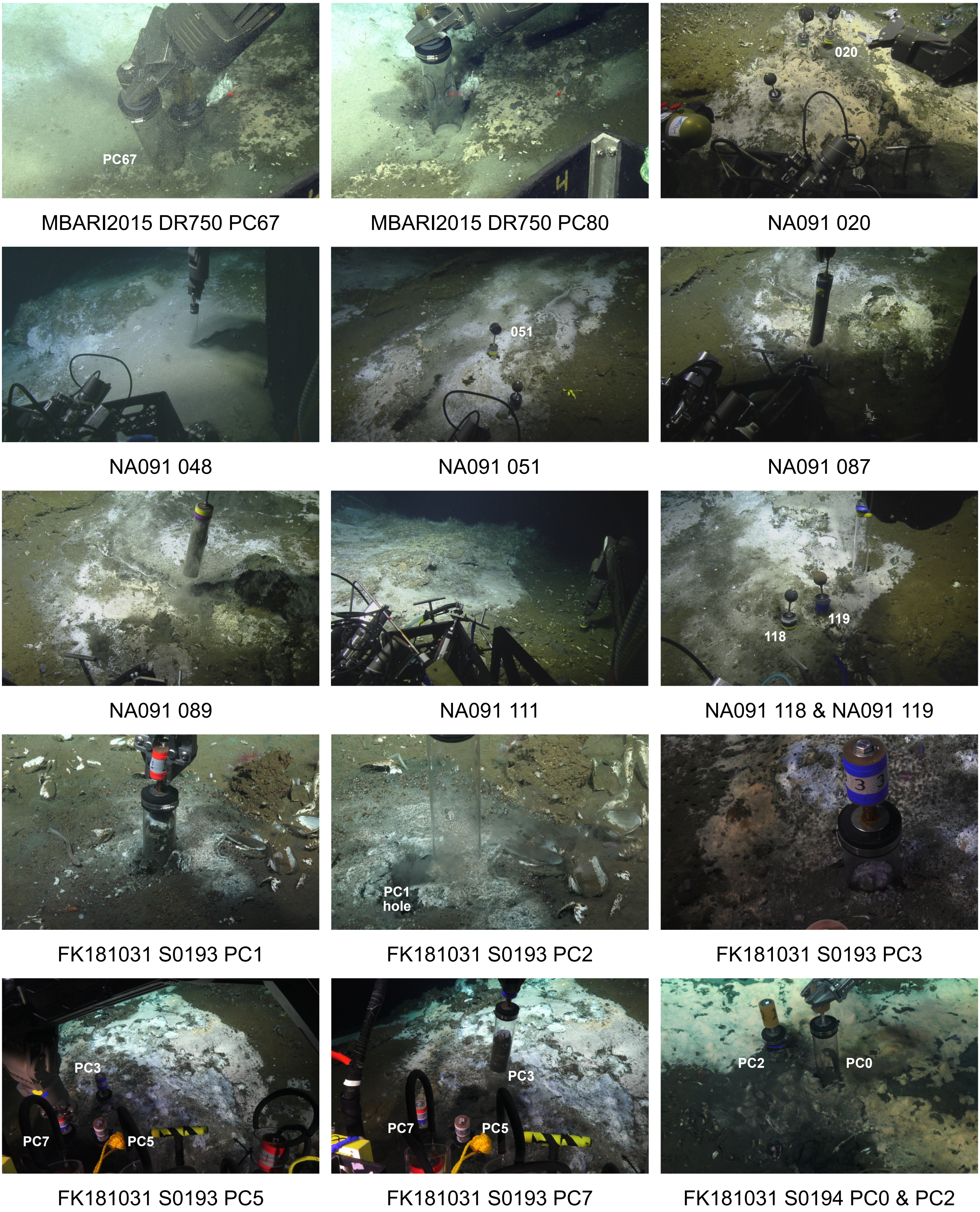


**Supplemental figure S1. Overview of sediment environments sampled using push cores**
Photos of sampling of the sediment push cores used in this study, sampled on RV Western Flyer using ROV Doc Ricketts (MBARI2015), EV Nautilus using ROV Hercules (NA091), and RV Falkor using ROV SuBastian (FK181031). In photos showing multiple cores, cores are labeled. Figure continues on the next page.

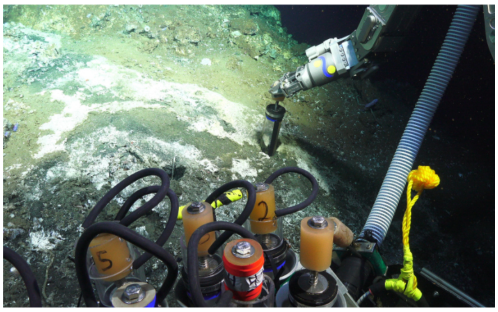

FK181031 S0194 PC1

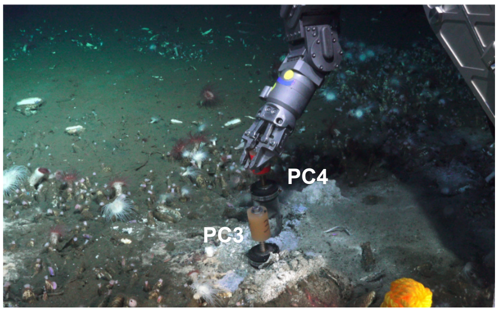


FK181031 S0194 PC3 & PC4

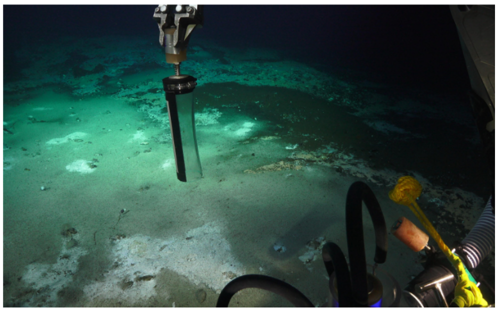

FK181031 S0196 PC1

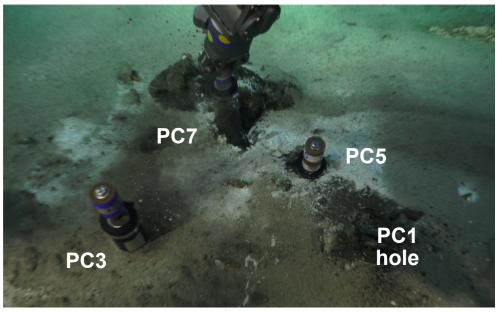


FK181031 S0196 PC5 & PC7

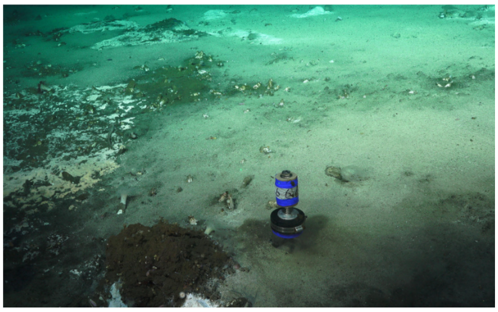

FK181031 S0196 PC6

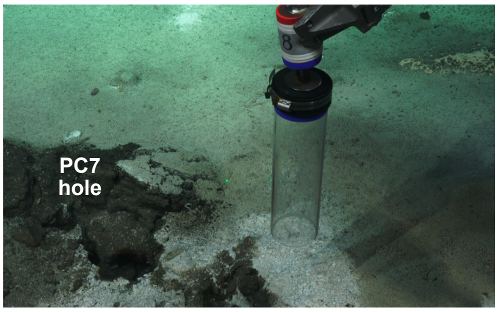


FK181031 S0196 PC8

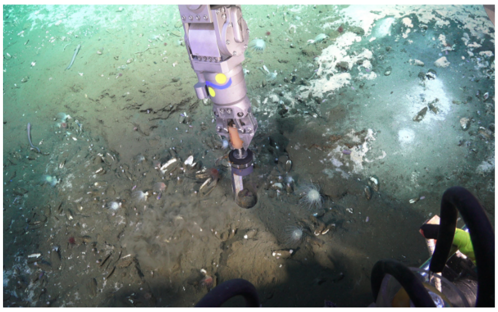

FK181031 S0198 PC1

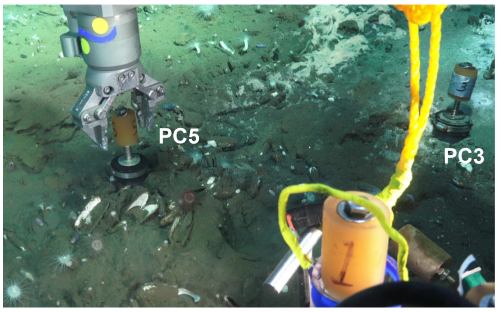


FK181031 S0198 PC3 & PC5

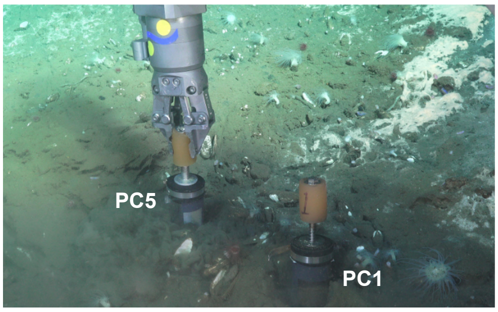


FK181031 S0198 PC1 & PC5

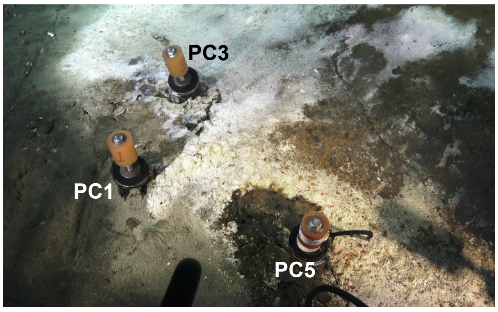


FK181031 S0200 PC1, PC3, PC5

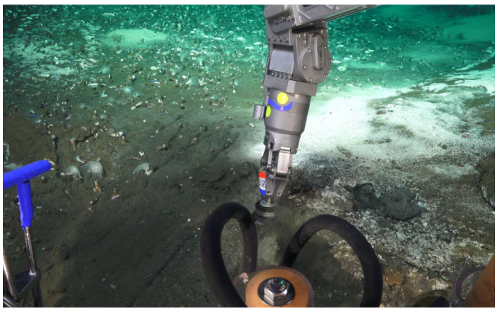

FK181031 S0200 PC7

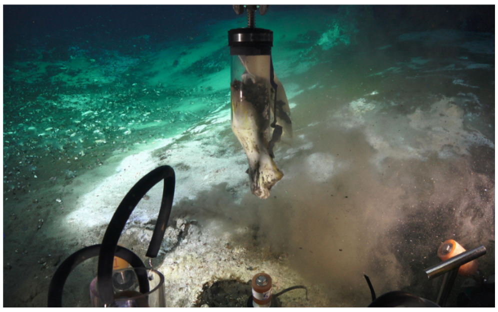

FK181031 S0200 PC3

**Supplemental figure S1. Overview of sediment environments sampled using push cores (cont.)** Photos of sampling of the sediment push cores used in this study, sampled on RV Western Flyer using ROV Doc Ricketts (MBARI2015), EV Nautilus using ROV Hercules (NA091), and RV Falkor using ROV SuBastian (FK181031). In photos showing multiple cores, cores are labeled. Page 2 of 2.

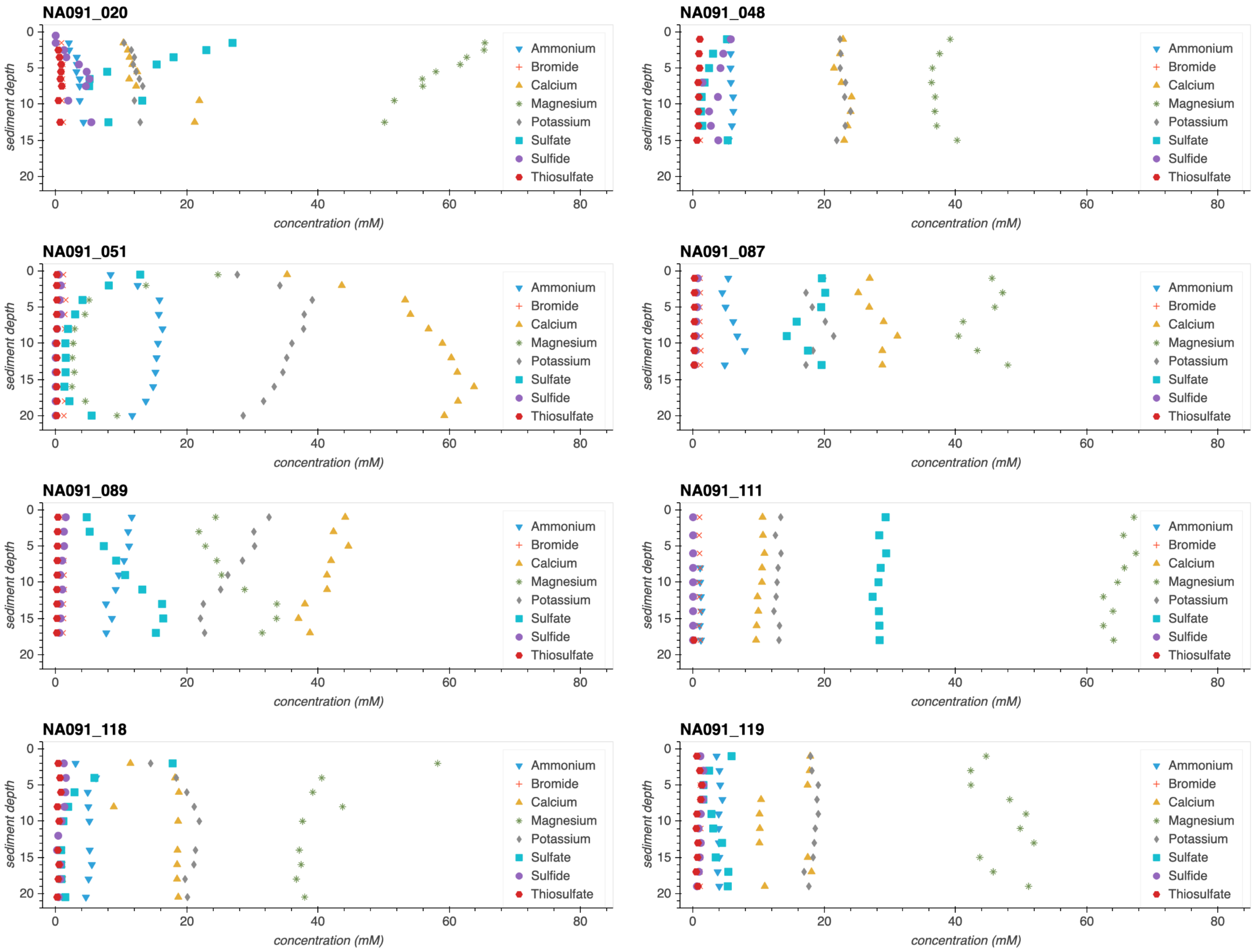


**Supplemental figure S2. Major ions in the pushcore samples from cruise NA091**
Concentrations of major ions as a function of depth in the sediments. Each point represents the porewater extracted from a sediment core horizon, plotted at the average depth of the horizon depth interval. All ions except for sulfide were measured using ion chromatography, and sulfide was measured colorimetrically.

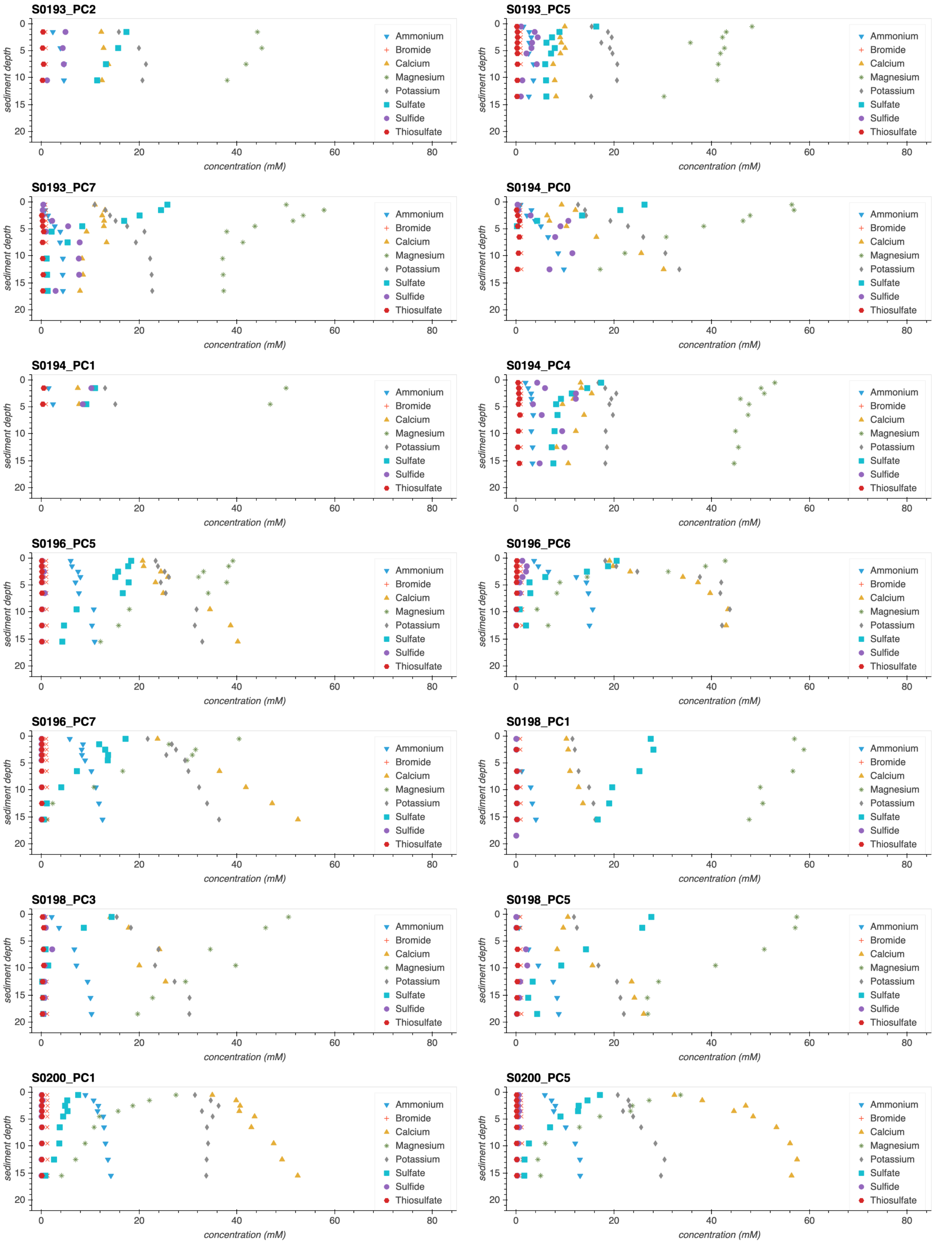


**Supplemental figure S3. Major ions in the pushcore samples from cruise FK181031**
Concentrations of major ions as a function of depth in the sediments. Each point represents the porewater extracted from a sediment core horizon, plotted at the average depth of the horizon depth interval. All ions except for sulfide were measured using ion chromatography, and sulfide was measured colorimetrically.

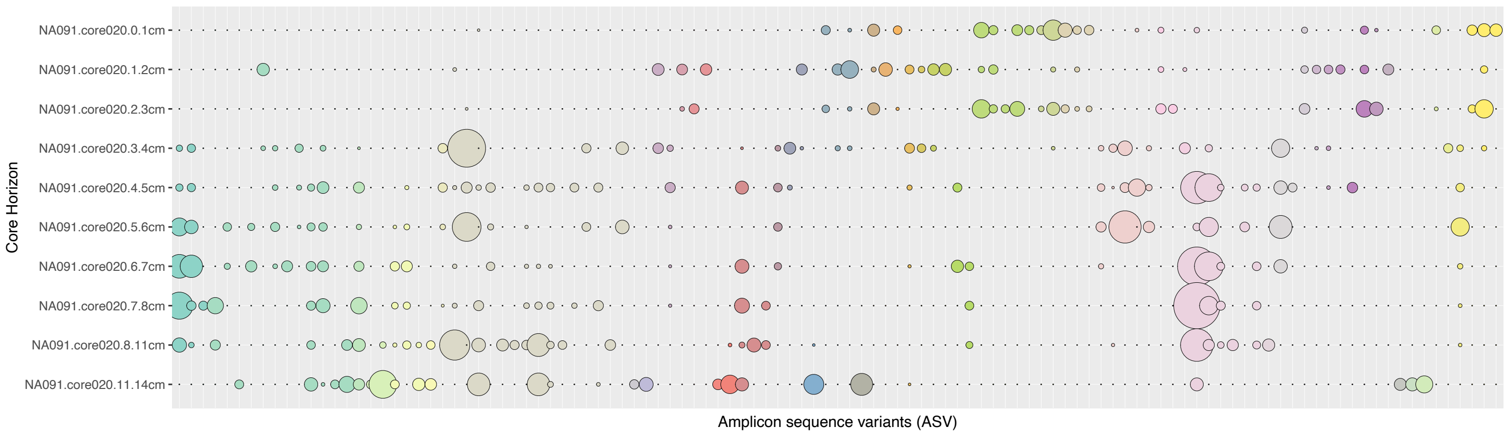


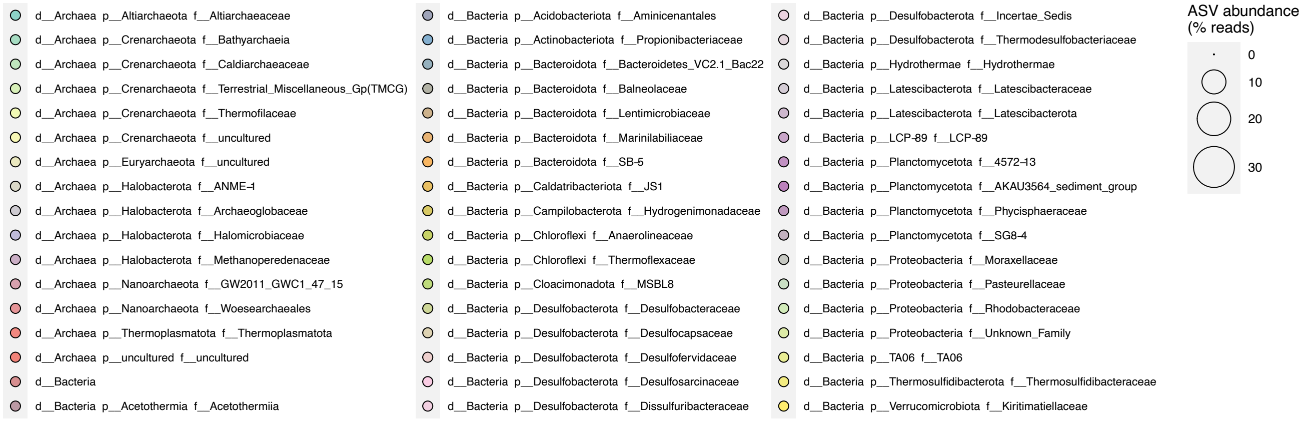


**Supplemental figure S4.**
NA091 020
ASV abundance

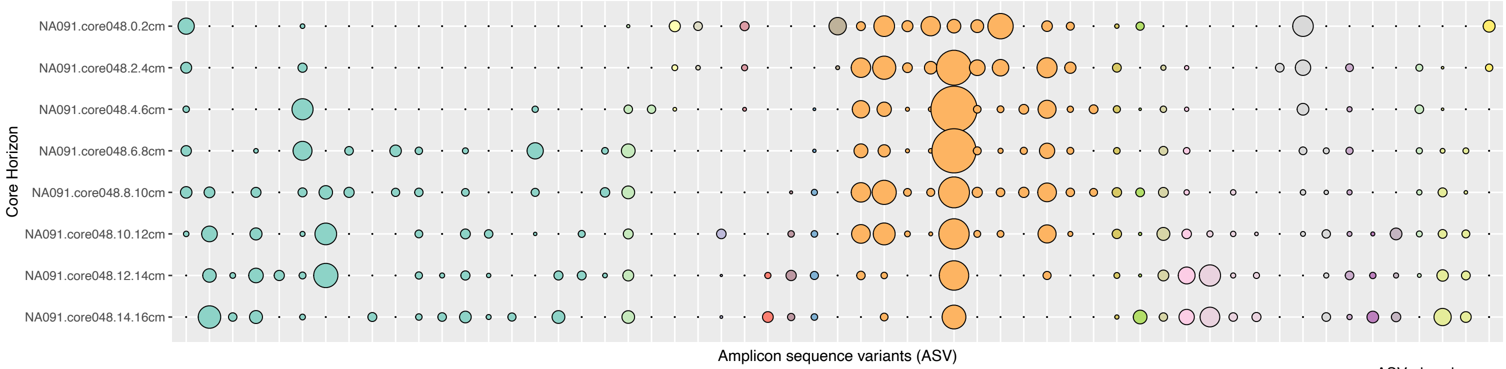


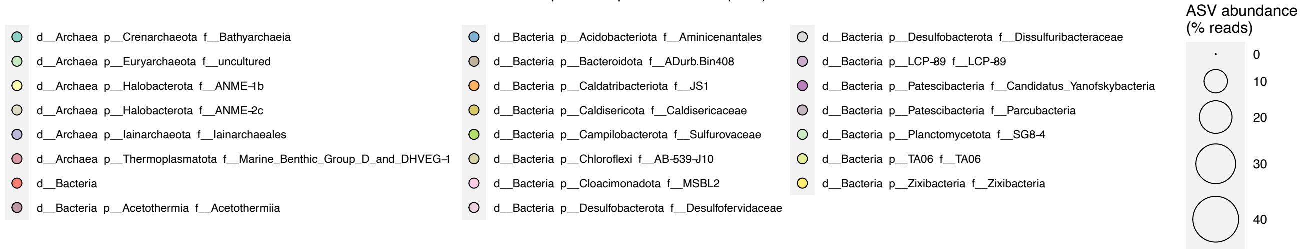


**Supplemental figure S5.**
NA091 048
ASV abundance

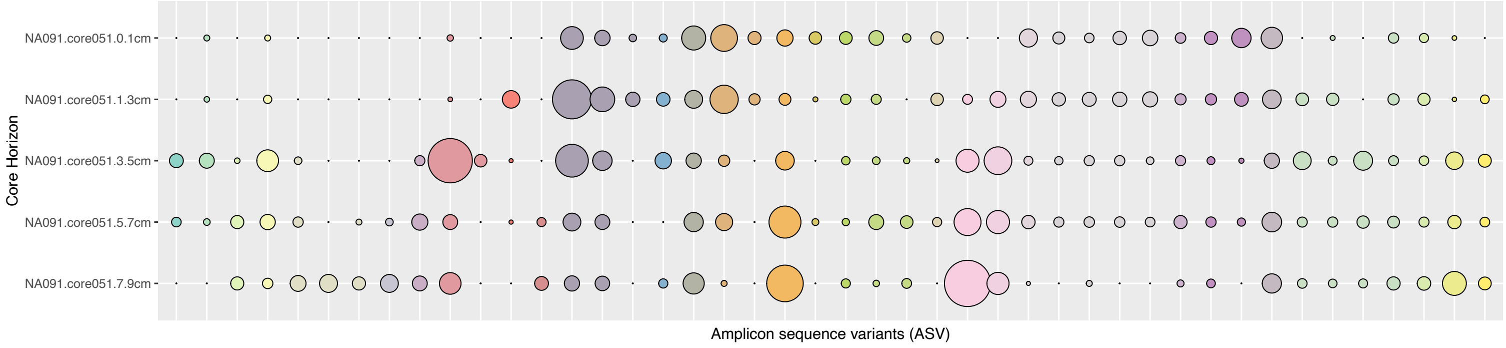


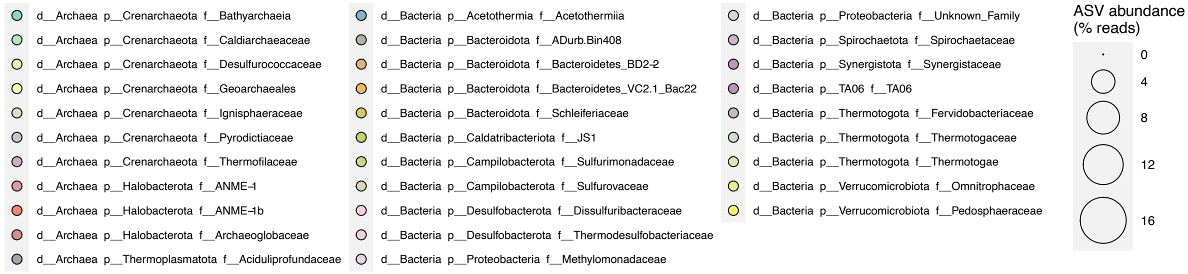


**Supplemental figure S6.**
NA091 051
ASV abundance

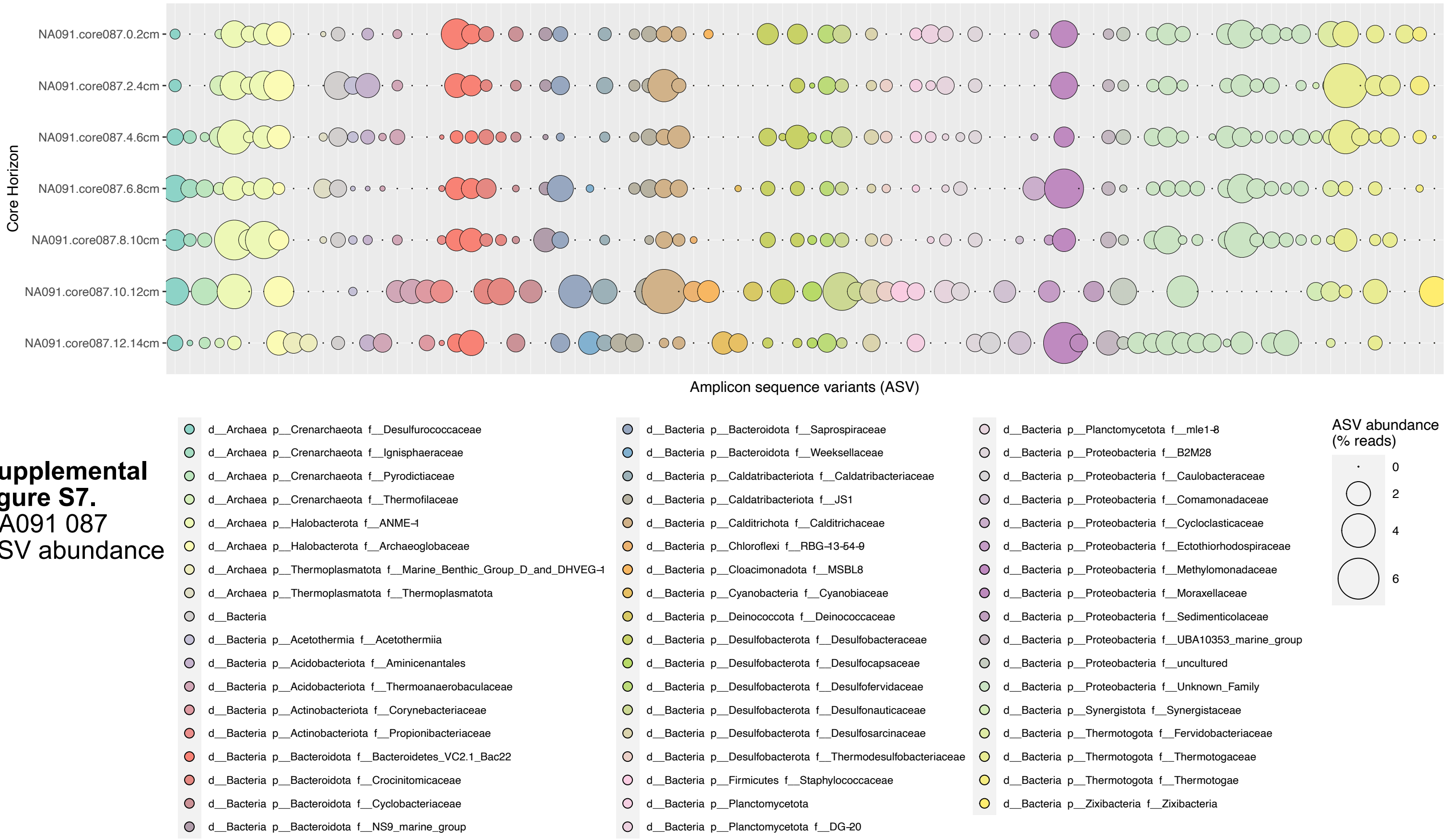


**Supplemental figure S7.** NA091 087 ASV abundance

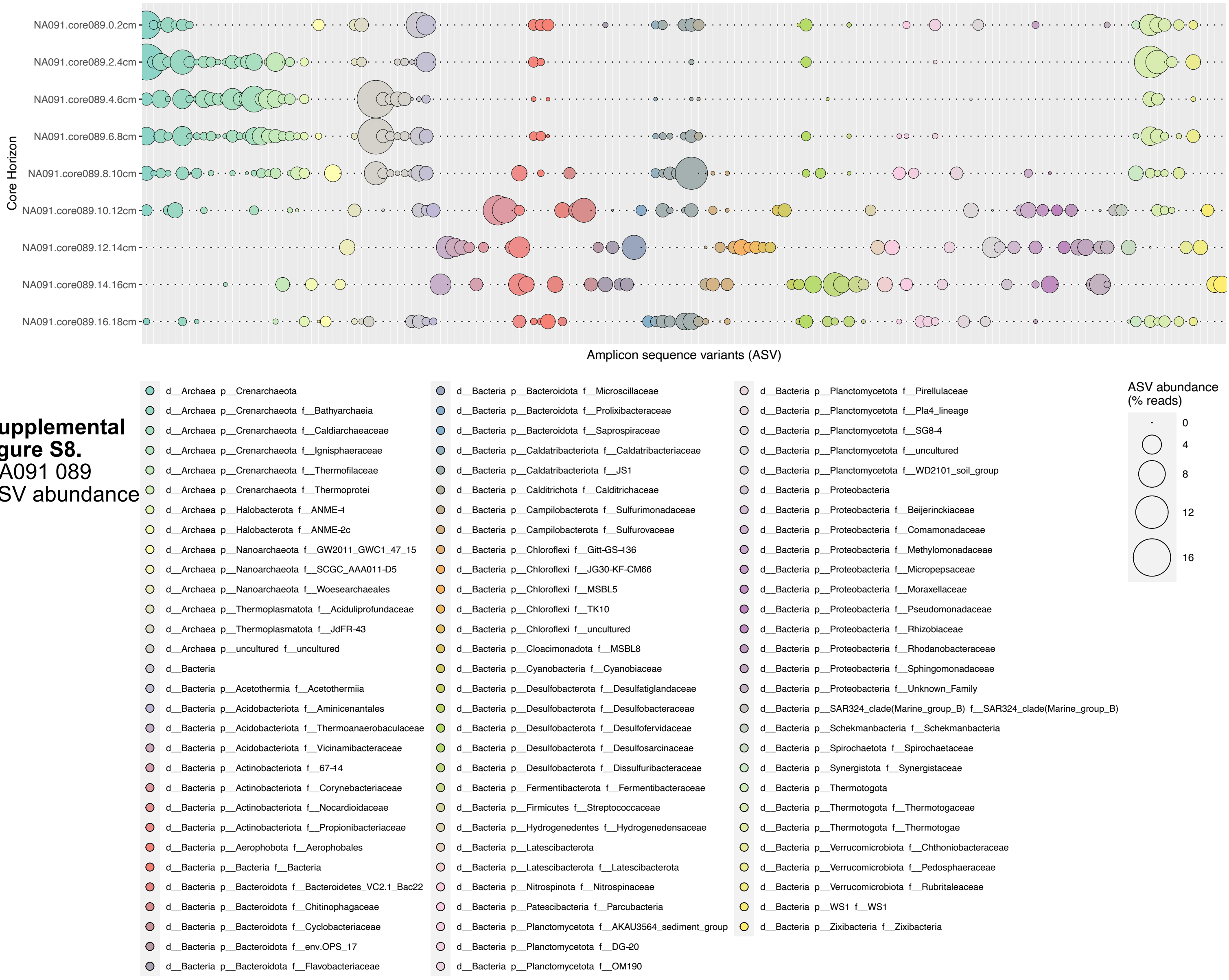


**Supplemental figure S8.** NA091 089 ASV abundance

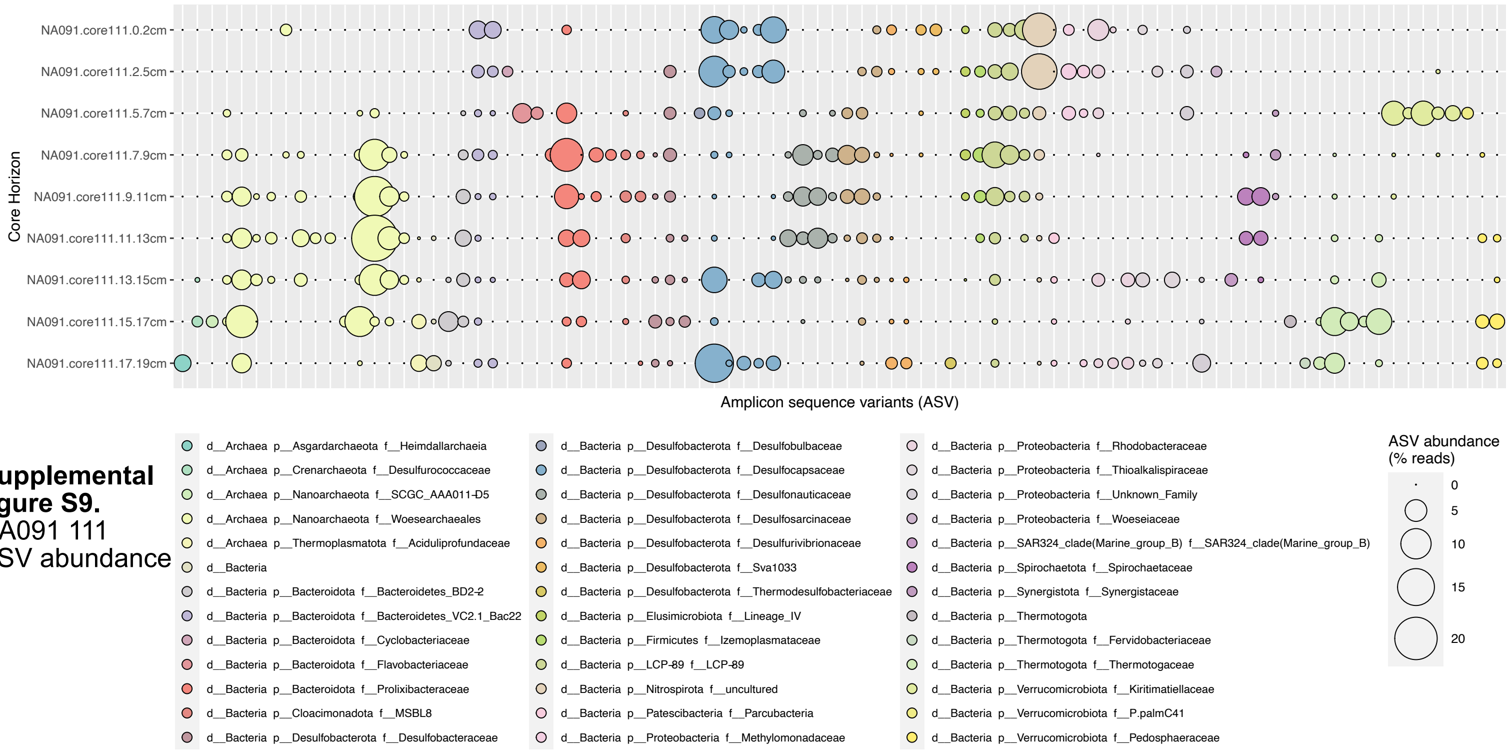


**Supplemental figure S9.**
NA091 111
ASV abundance

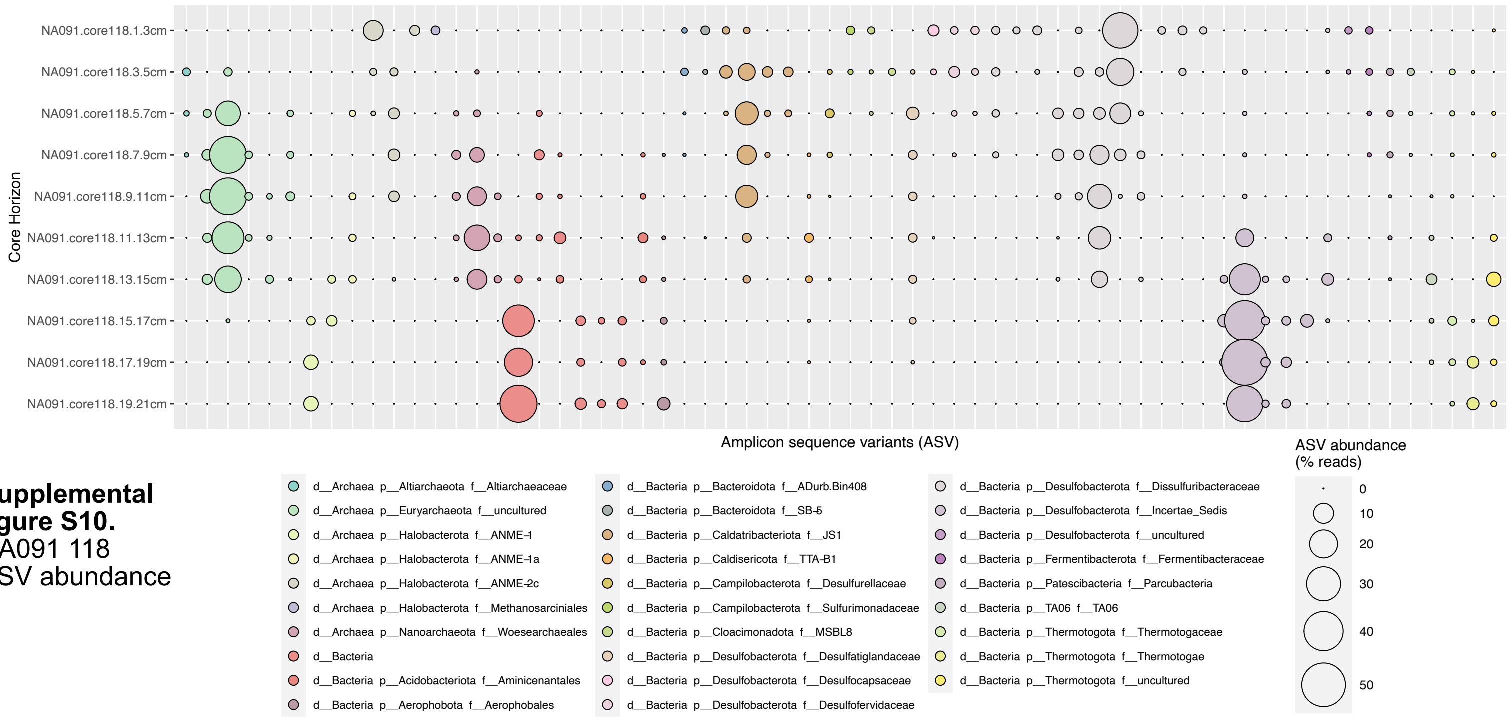


**Supplemental figure S10.**
NA091 118
ASV abundance

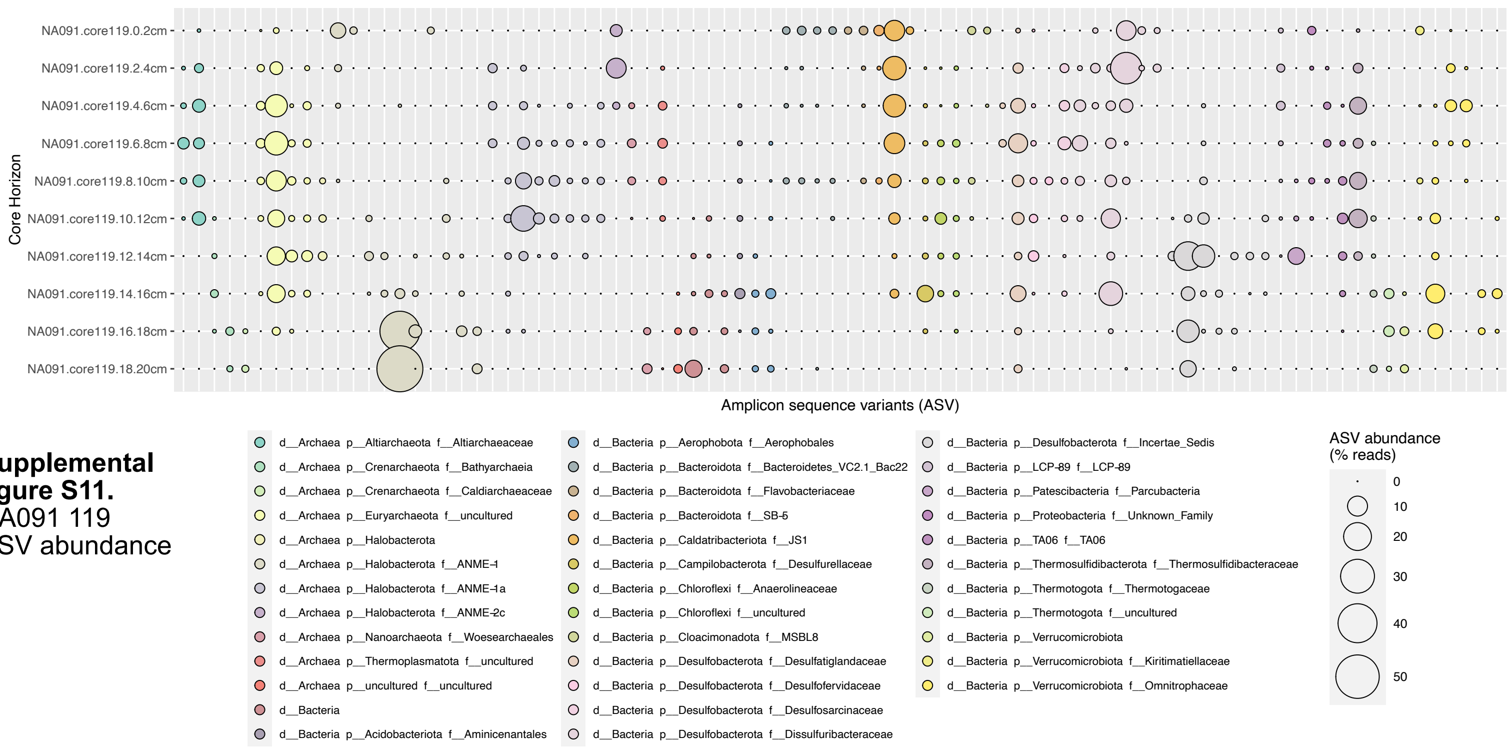


**Supplemental figure S11.**
NA091 119
ASV abundance

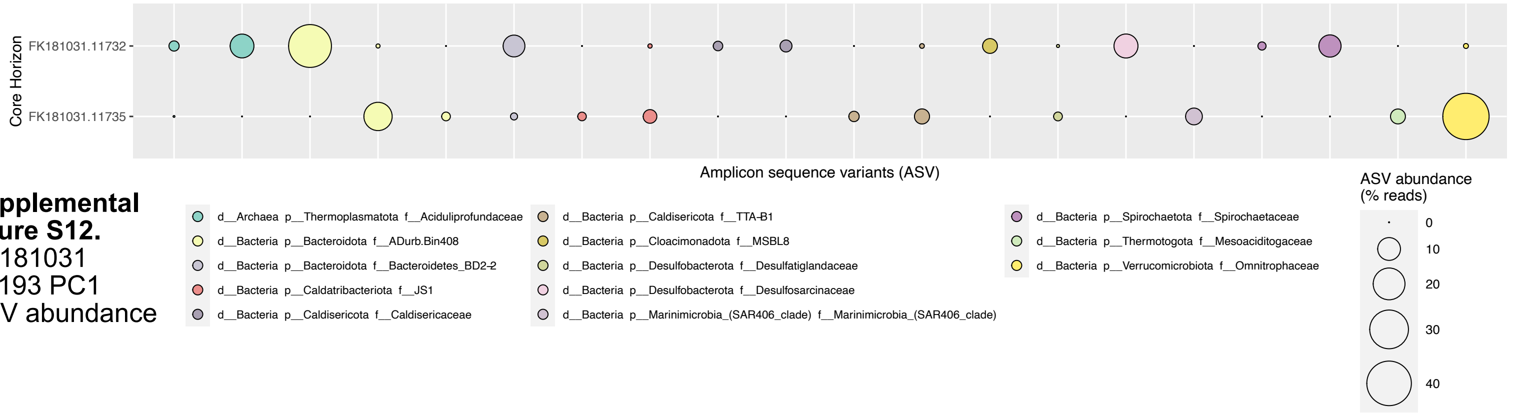


**Supplemental figure S12.**
FK181031
S0193 PC1
ASV abundance

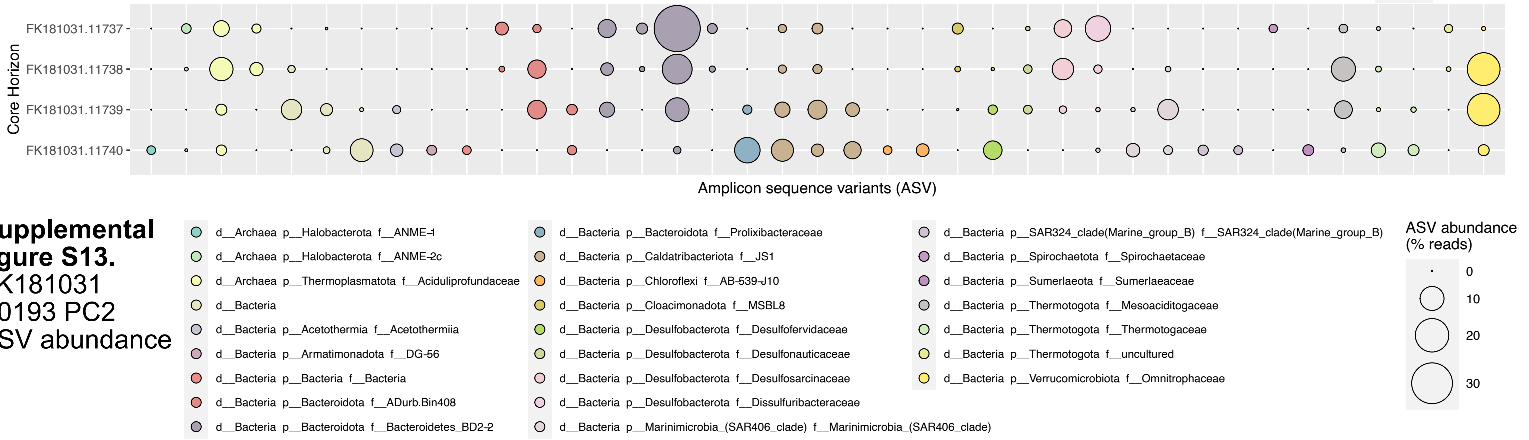


**Supplemental figure S13.**
FK181031
S0193 PC2
ASV abundance

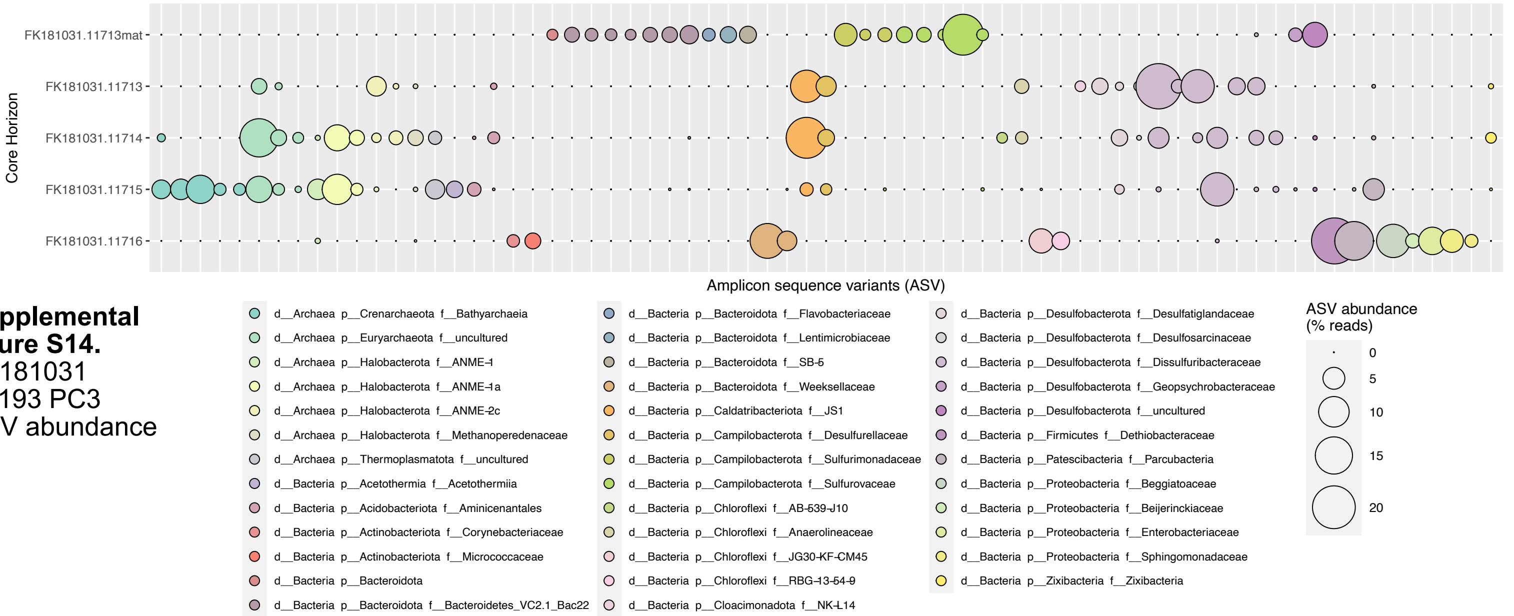


**Supplemental figure S14.**
FK181031
S0193 PC3
ASV abundance

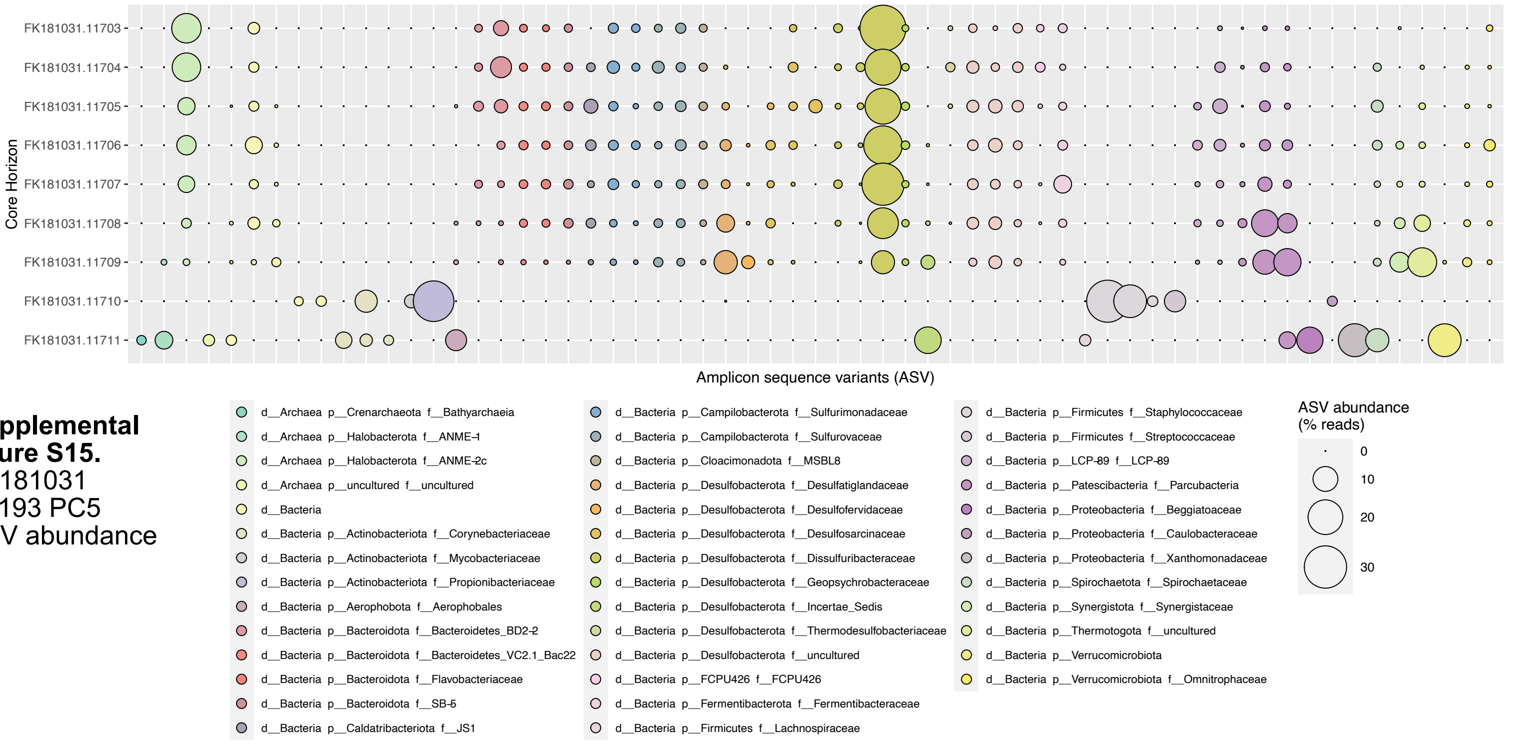


**Supplemental figure S15.**
FK181031
S0193 PC5
ASV abundance

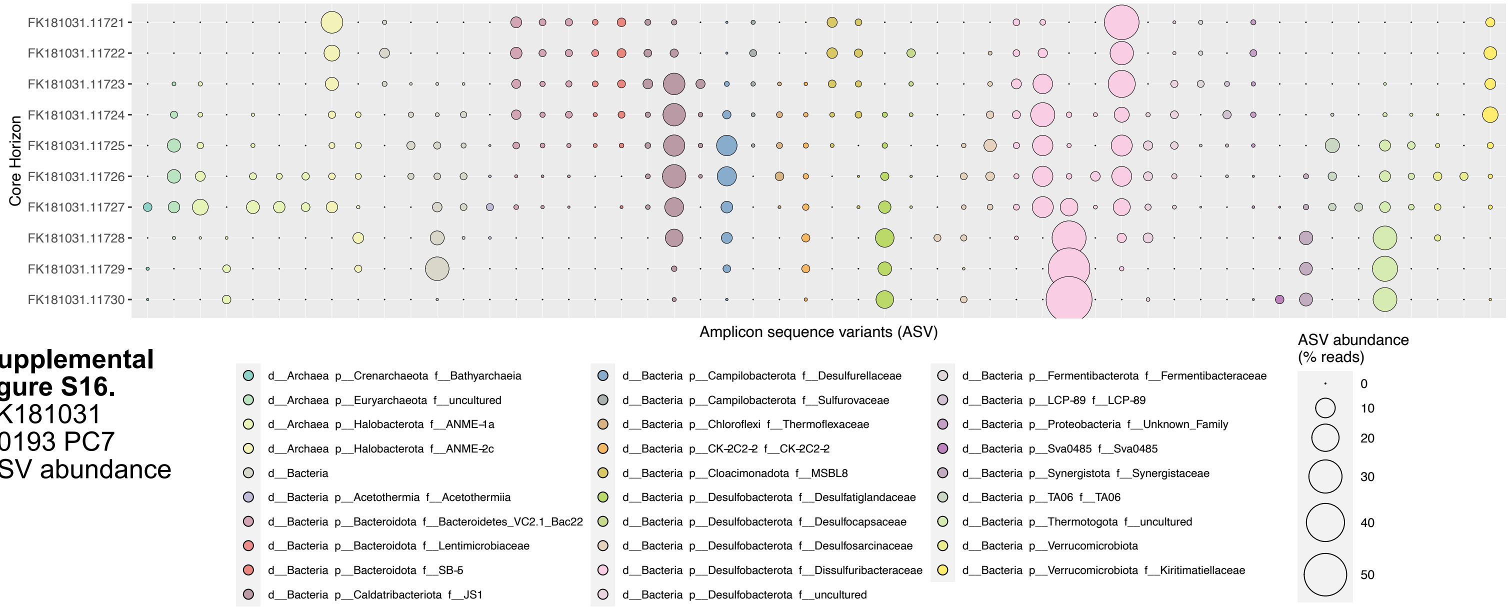


**Supplemental figure S16.**
FK181031
S0193 PC7
ASV abundance

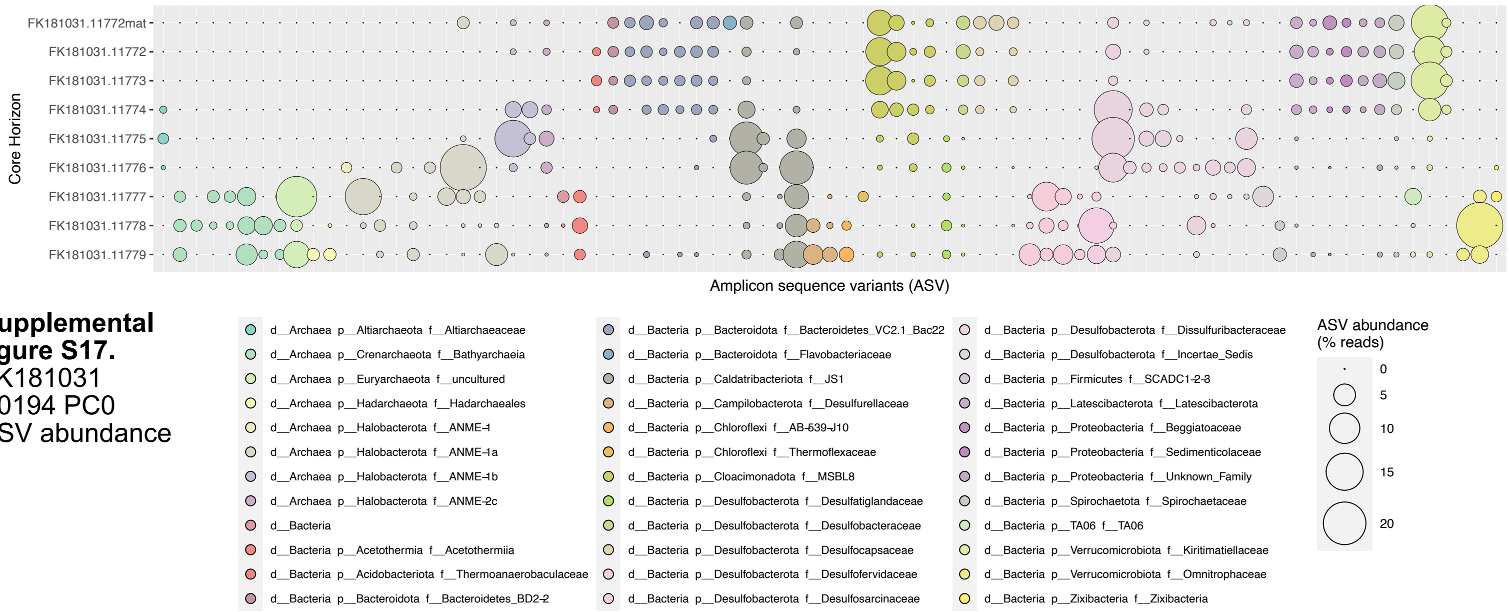


**Supplemental figure S17.**
FK181031
S0194 PC0
ASV abundance

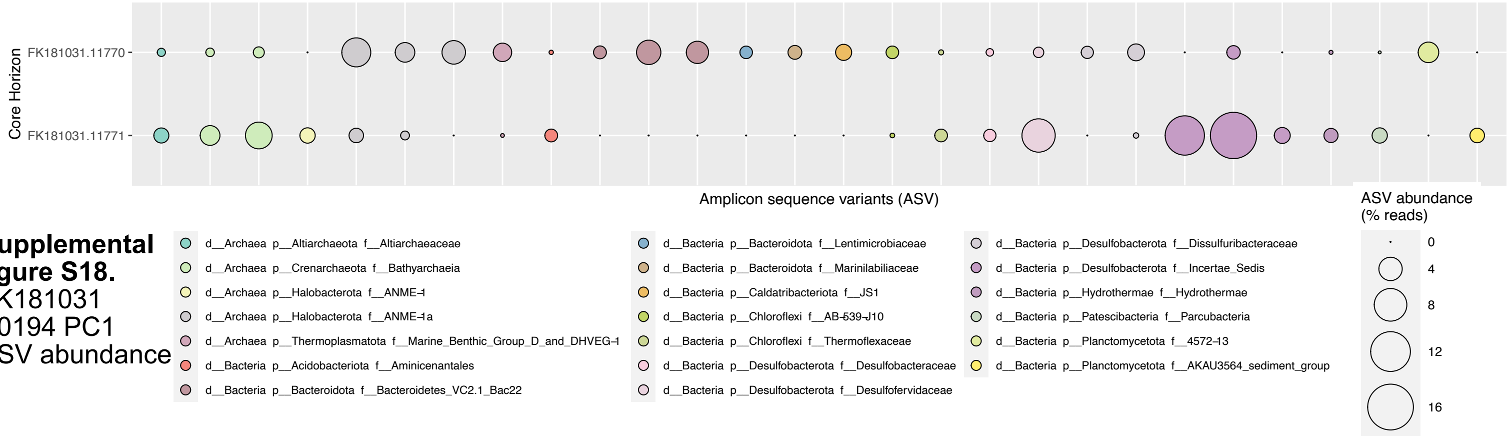


**Supplemental figure S18.**
FK181031
S0194 PC1
ASV abundance

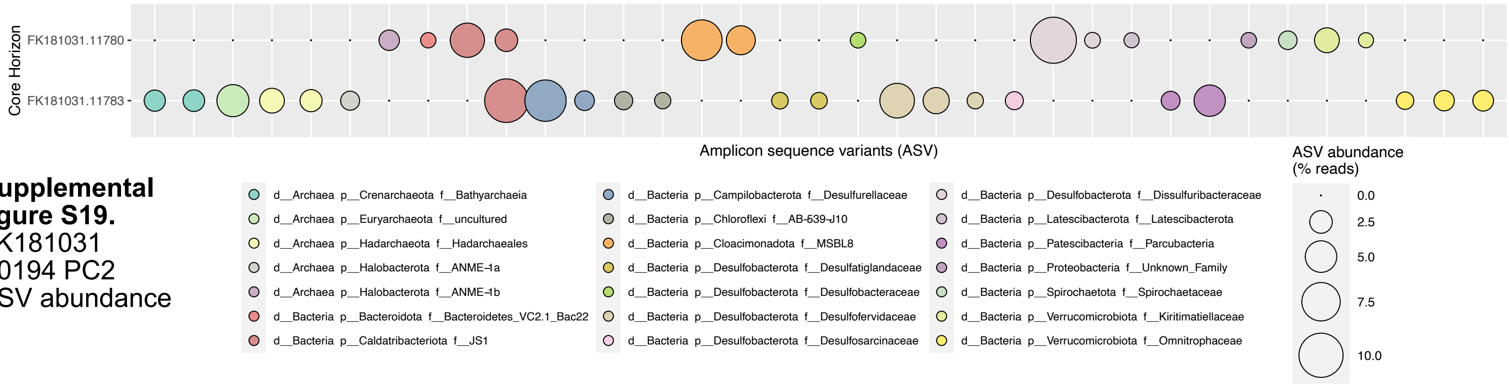


**Supplemental figure S19.**
FK181031
S0194 PC2
ASV abundance

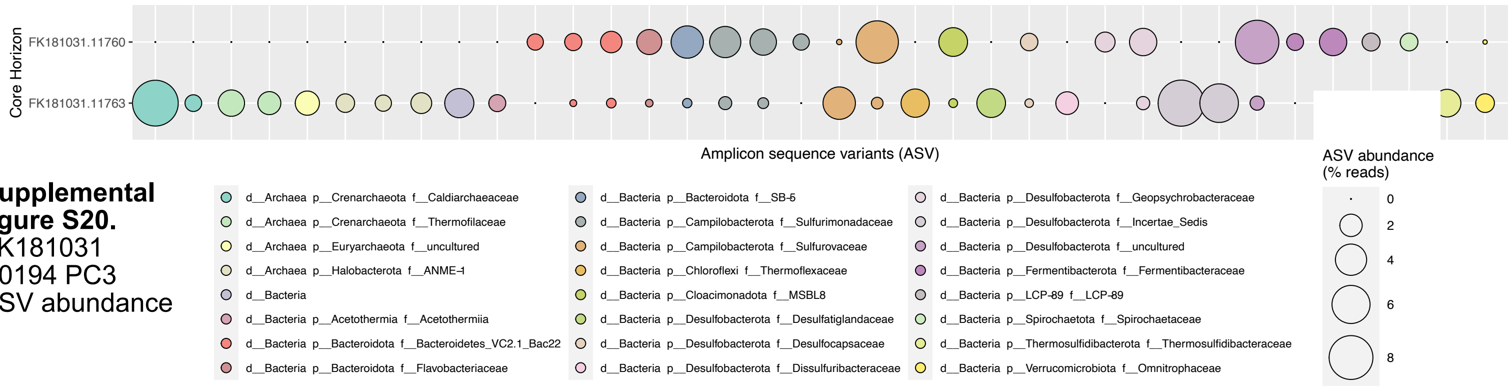


**Supplemental figure S20.**
FK181031
S0194 PC3
ASV abundance

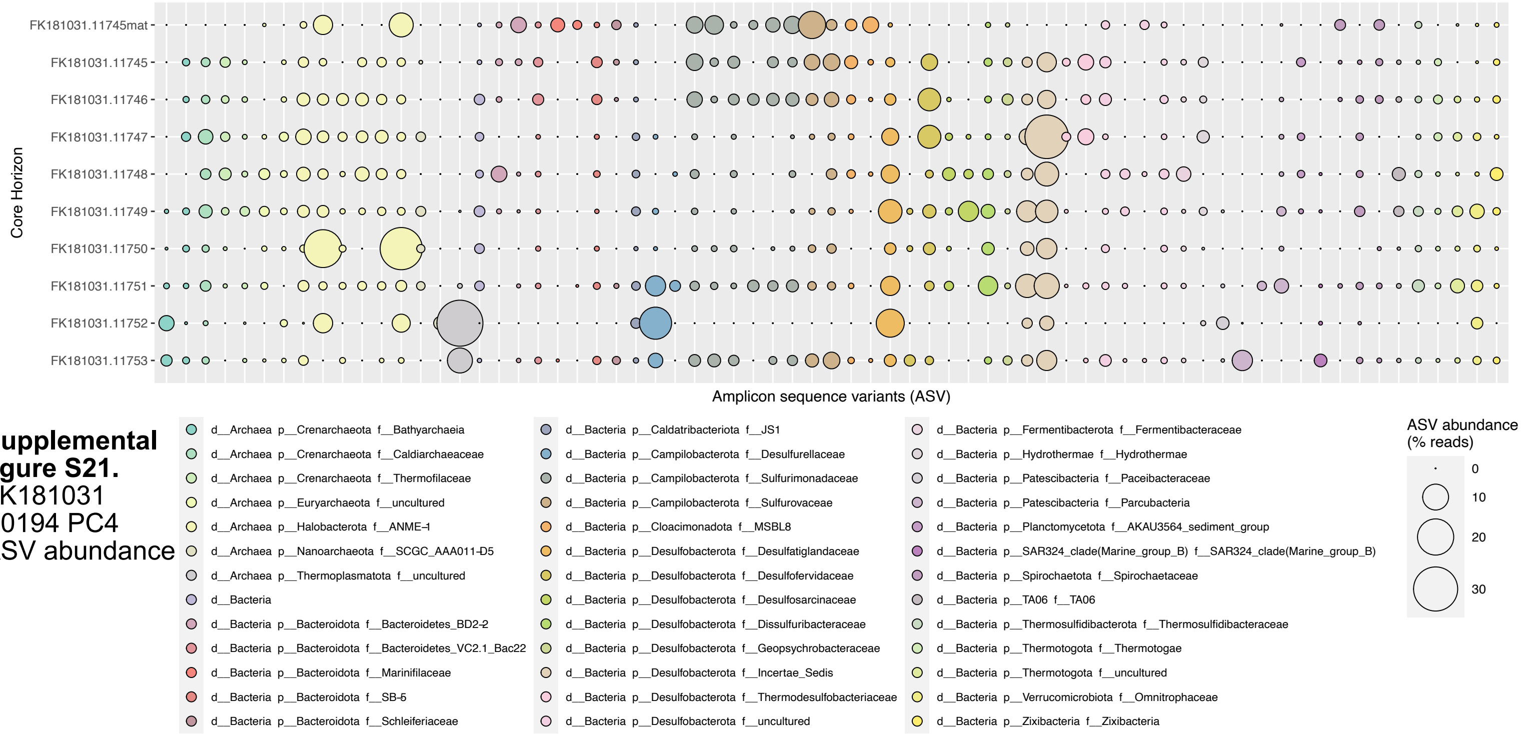


**Supplemental figure S21.**
FK181031
S0194 PC4
ASV abundance

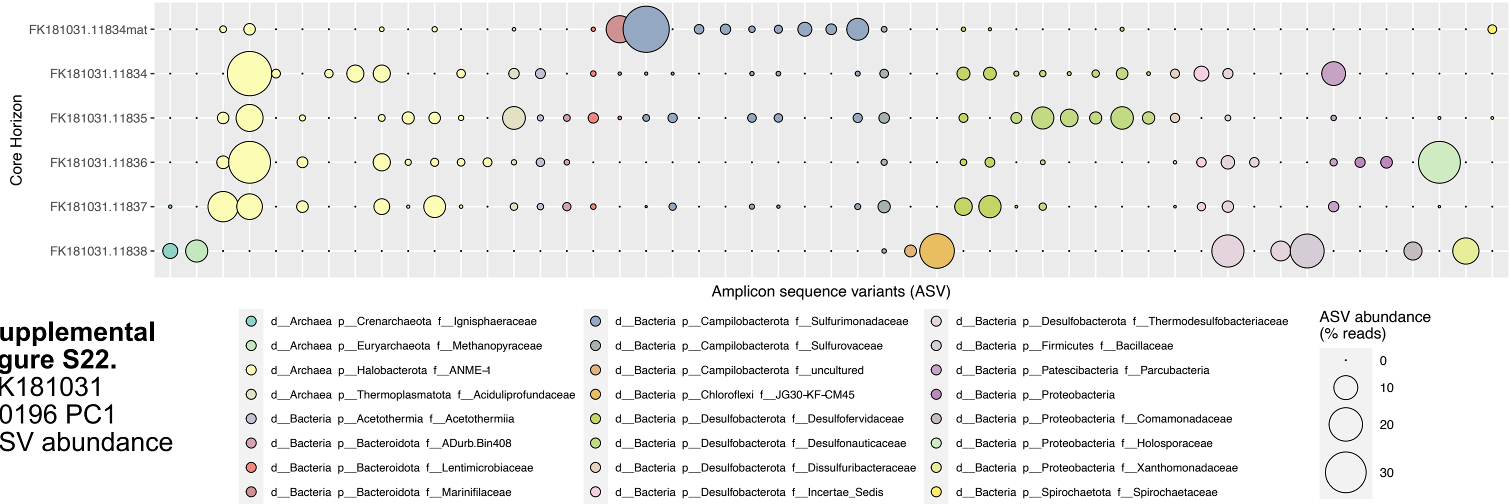


**Supplemental figure S22.**
FK181031
S0196 PC1
ASV abundance

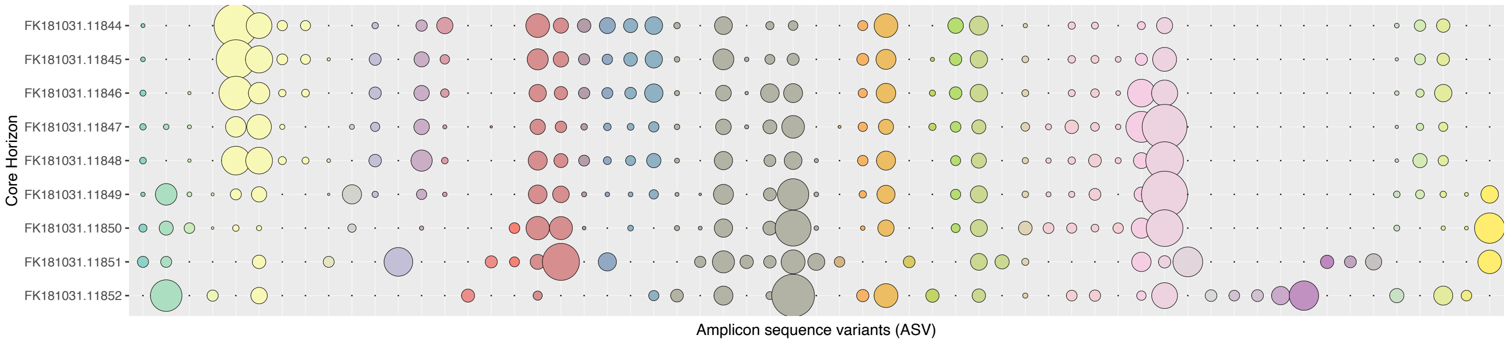


**Supplemental figure S23.**
FK181031
S0196 PC5
ASV abundance

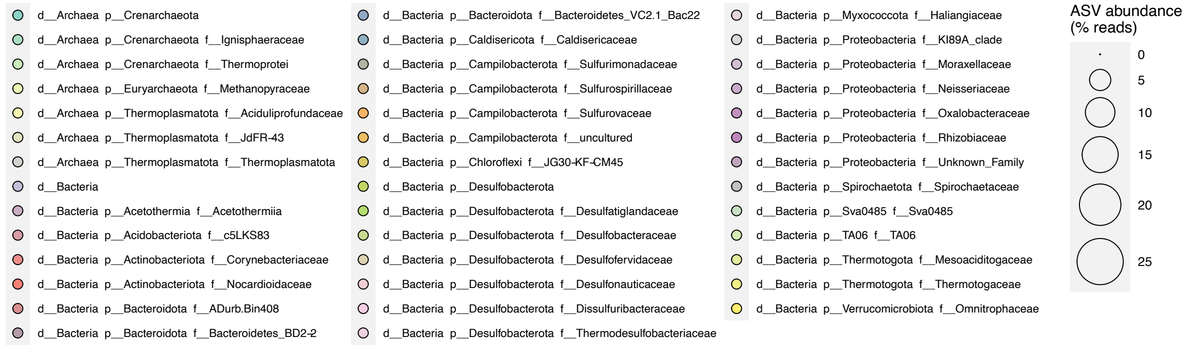


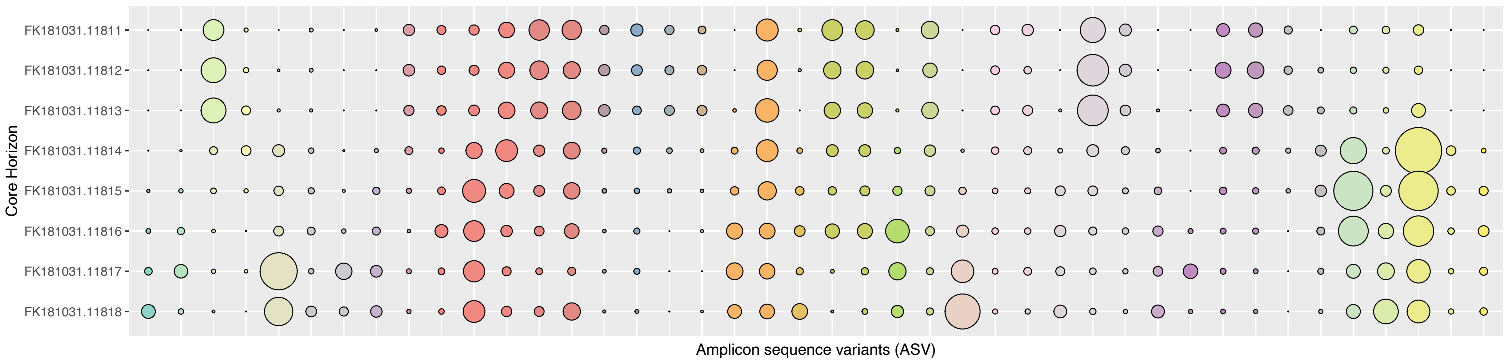


**Supplemental figure S24.**
FK181031
S0196 PC6
ASV abundance

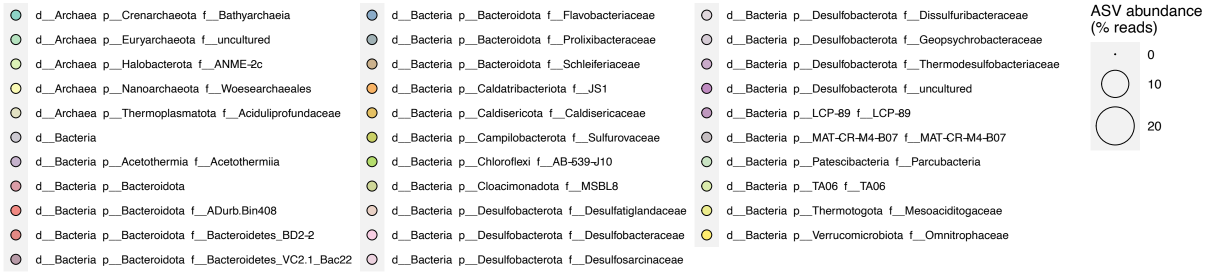


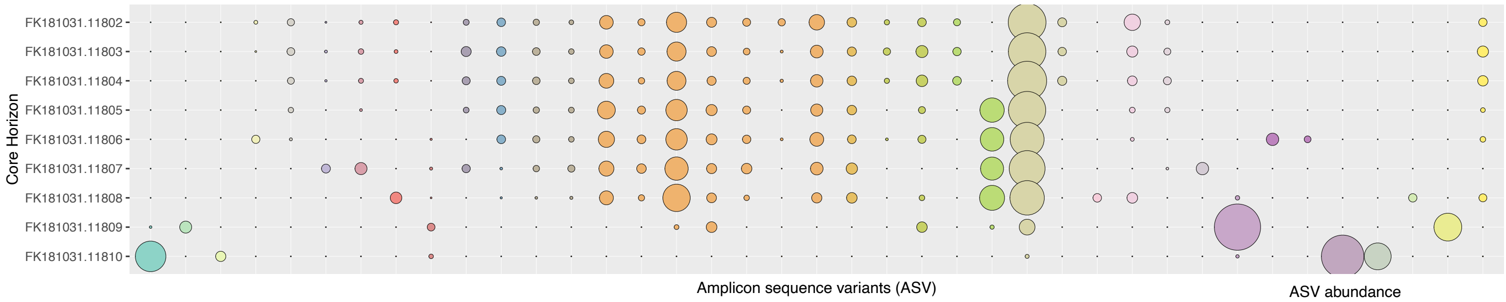


**Supplemental figure S25.**
FK181031
S0196 PC7
ASV abundance

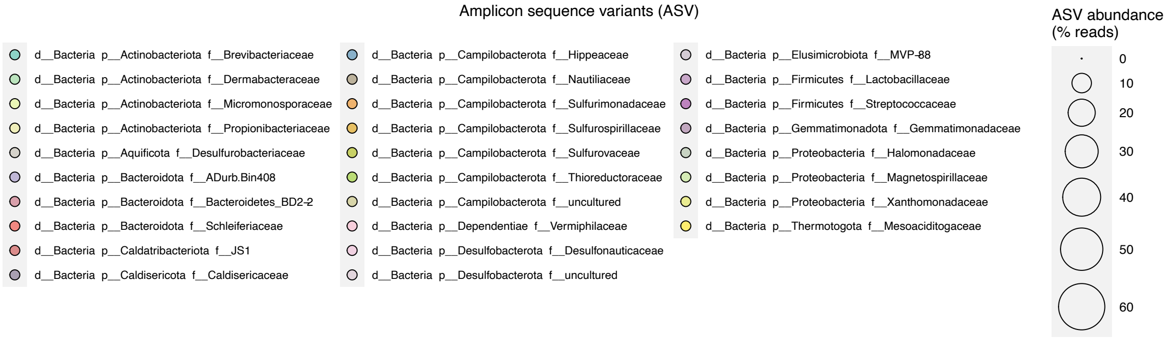

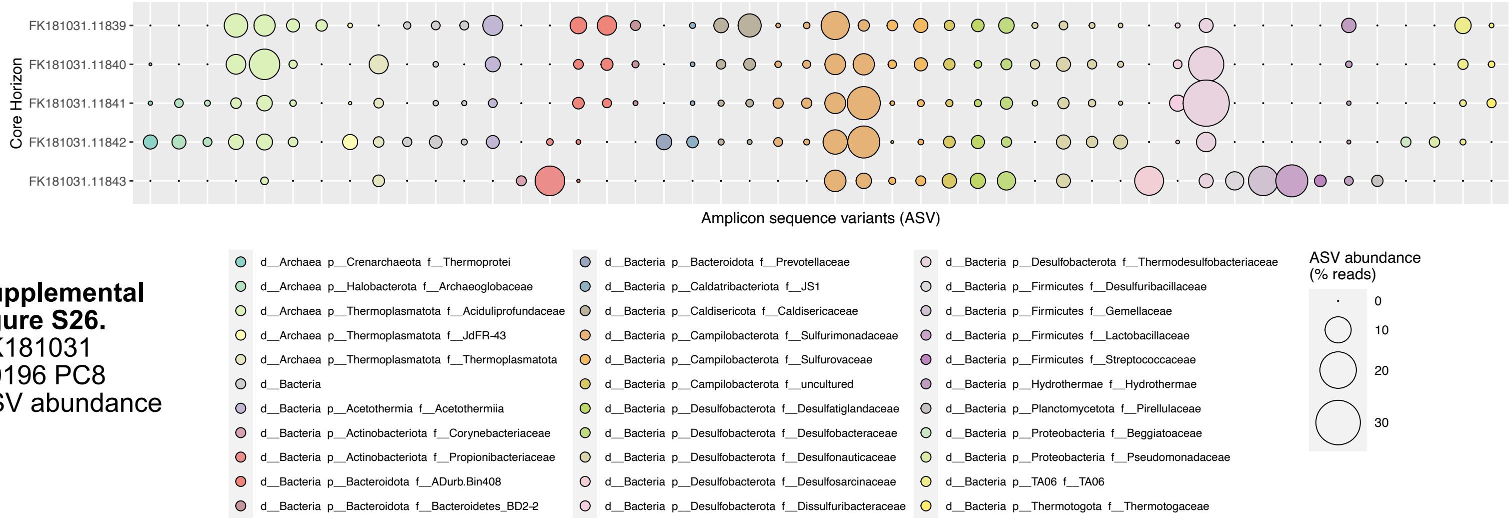


**Supplemental figure S26.** FK181031 S0196 PC8 ASV abundance

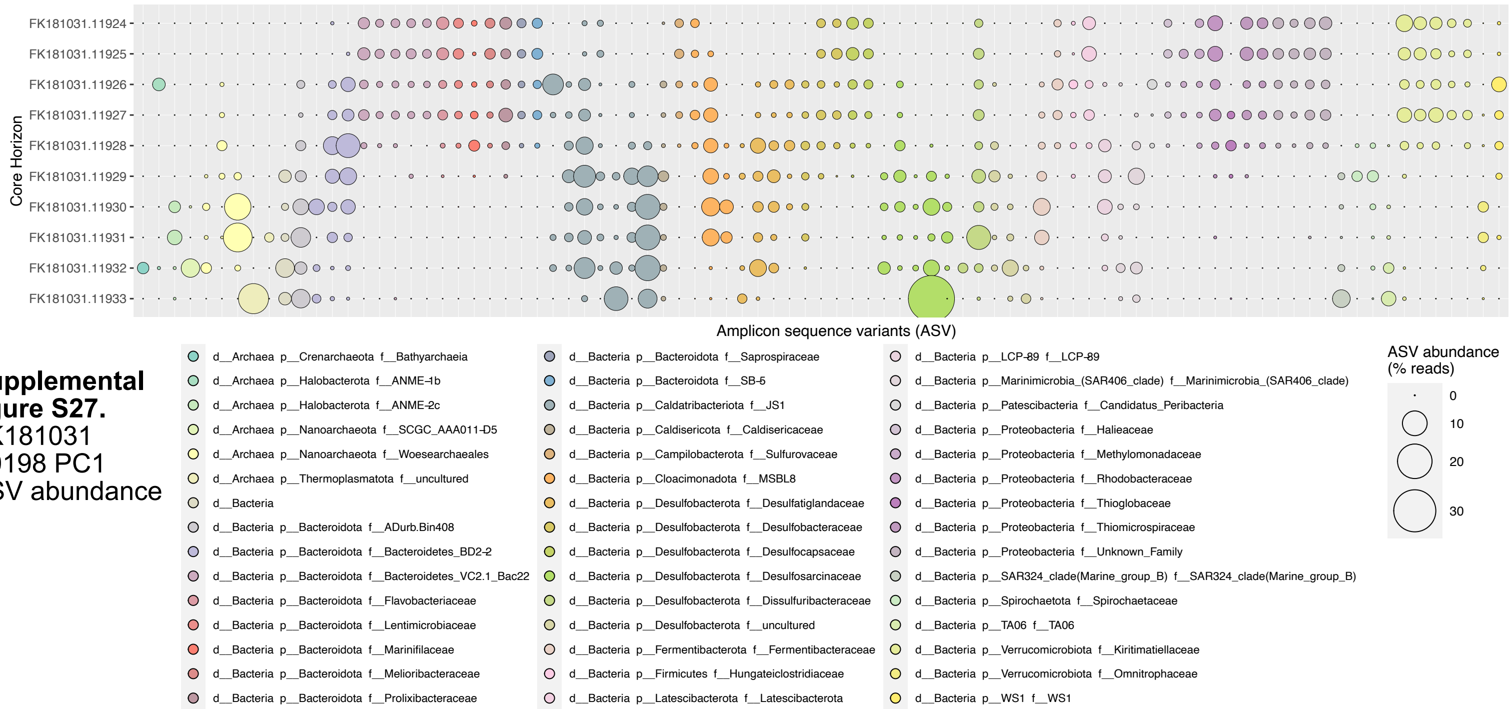


**Supplemental figure S27.** FK181031 S0198 PC1 ASV abundance

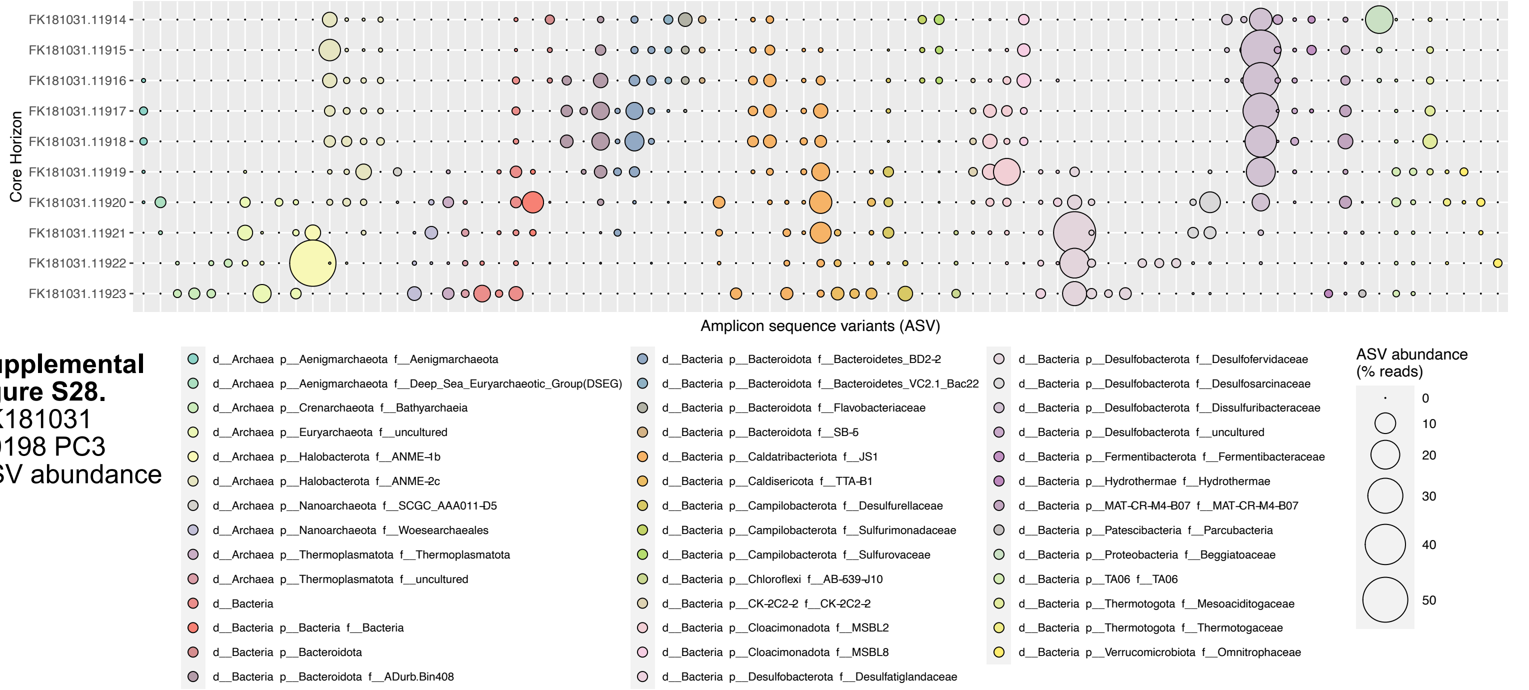


**Supplemental figure S28.** FK181031 S0198 PC3 ASV abundance

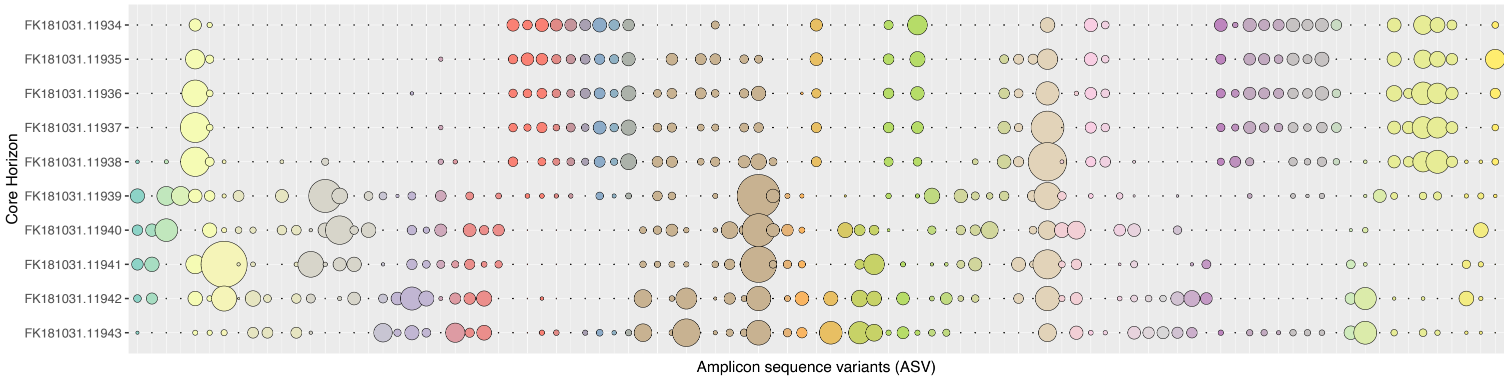


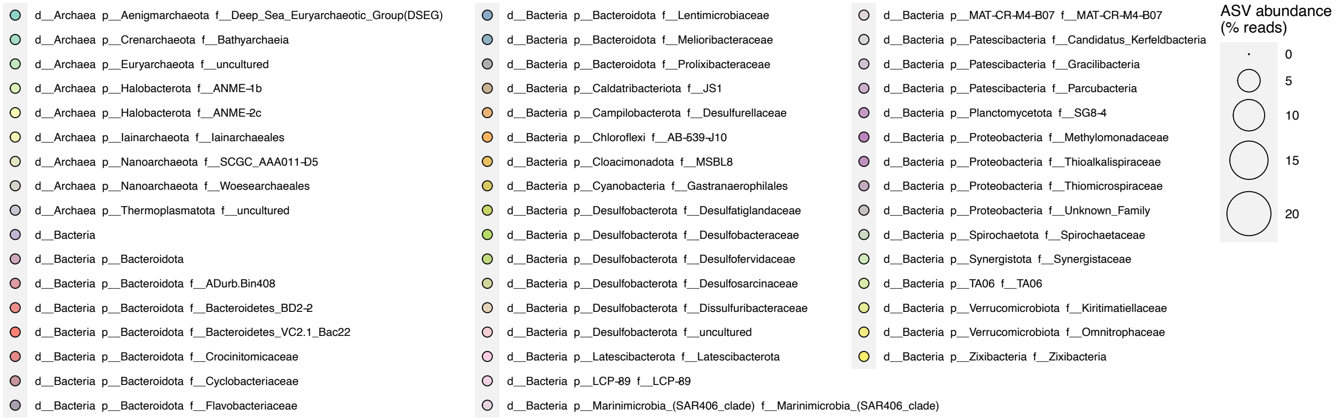


**Supplemental figure S29.**
FK181031
S0198 PC5
ASV abundance

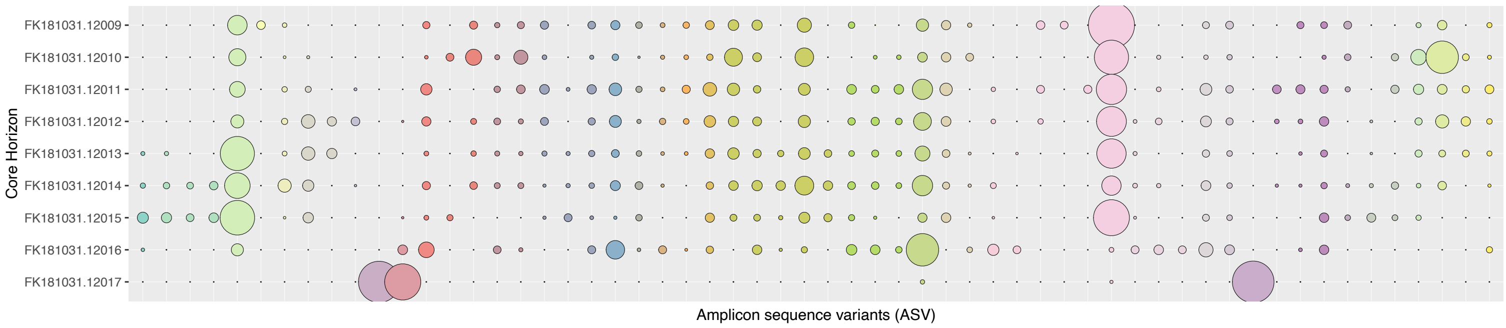


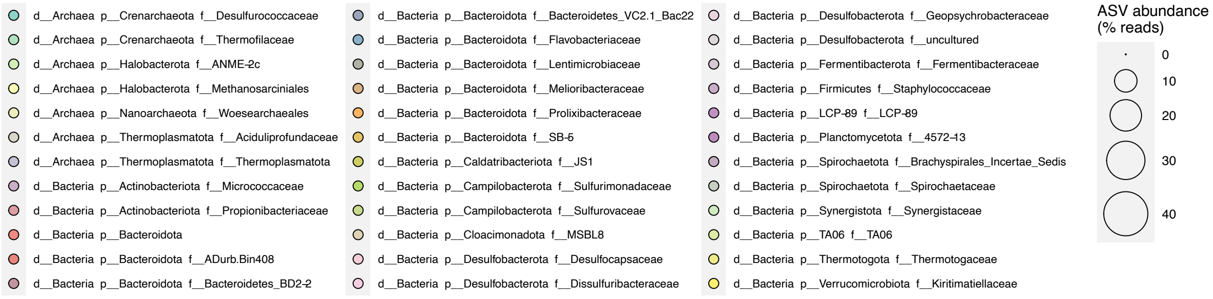


**Supplemental figure S30.**
FK181031
S0200 PC1
ASV abundance

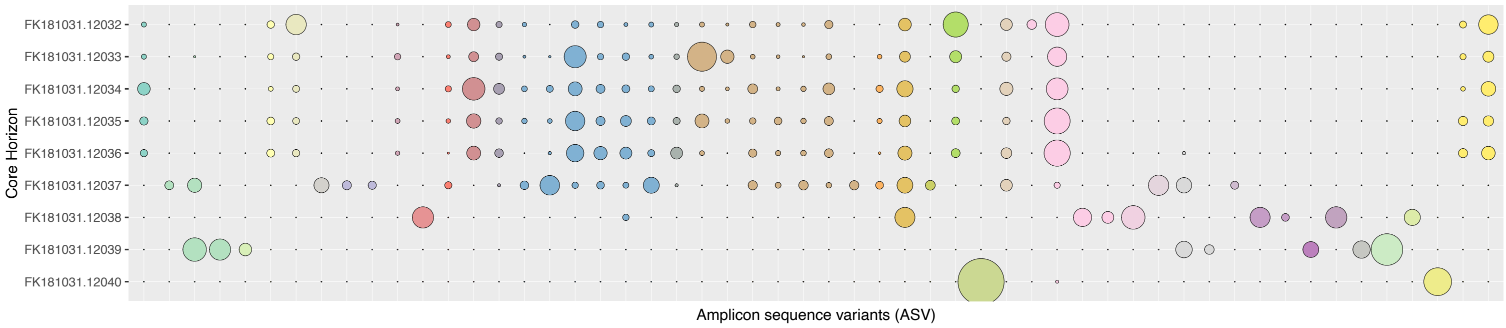


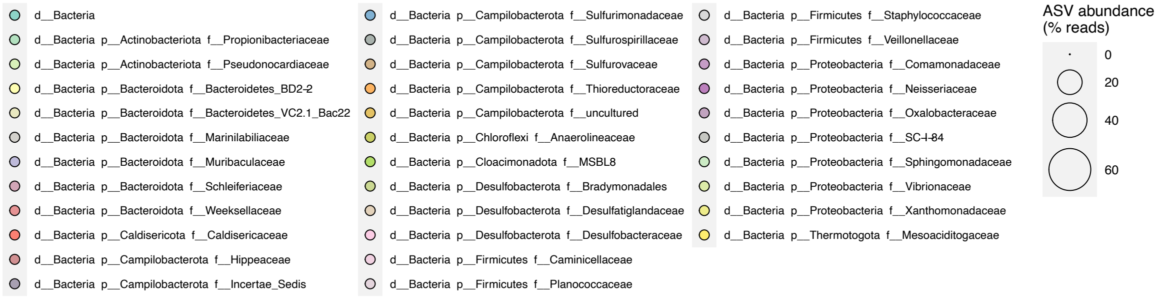


**Supplemental figure S31.**
FK181031
S0200 PC5
ASV abundance

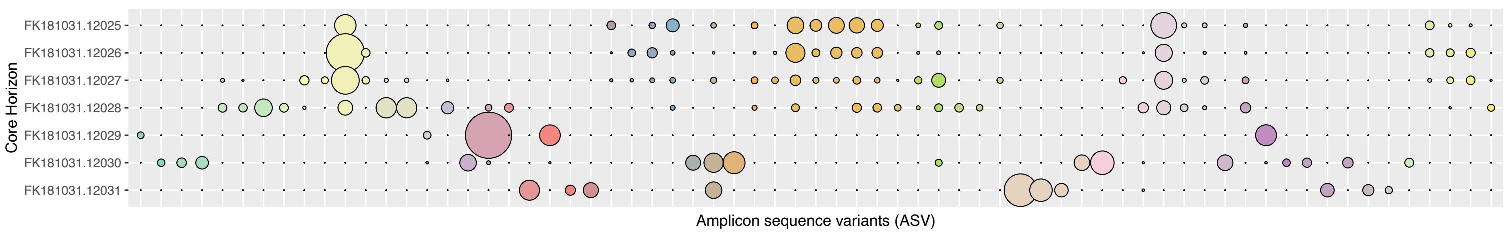


### **Supplemental figure S30.**
FK181031
S0200 PC7
ASV abundance

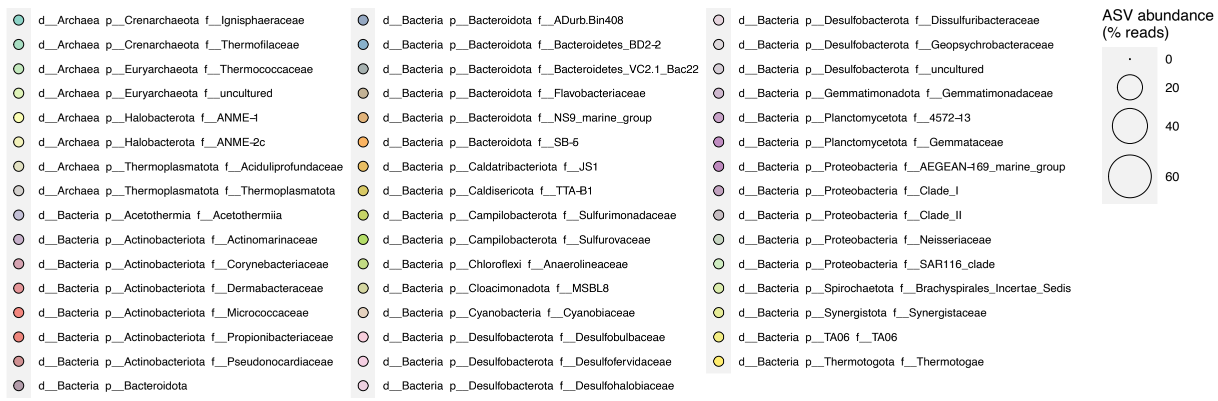

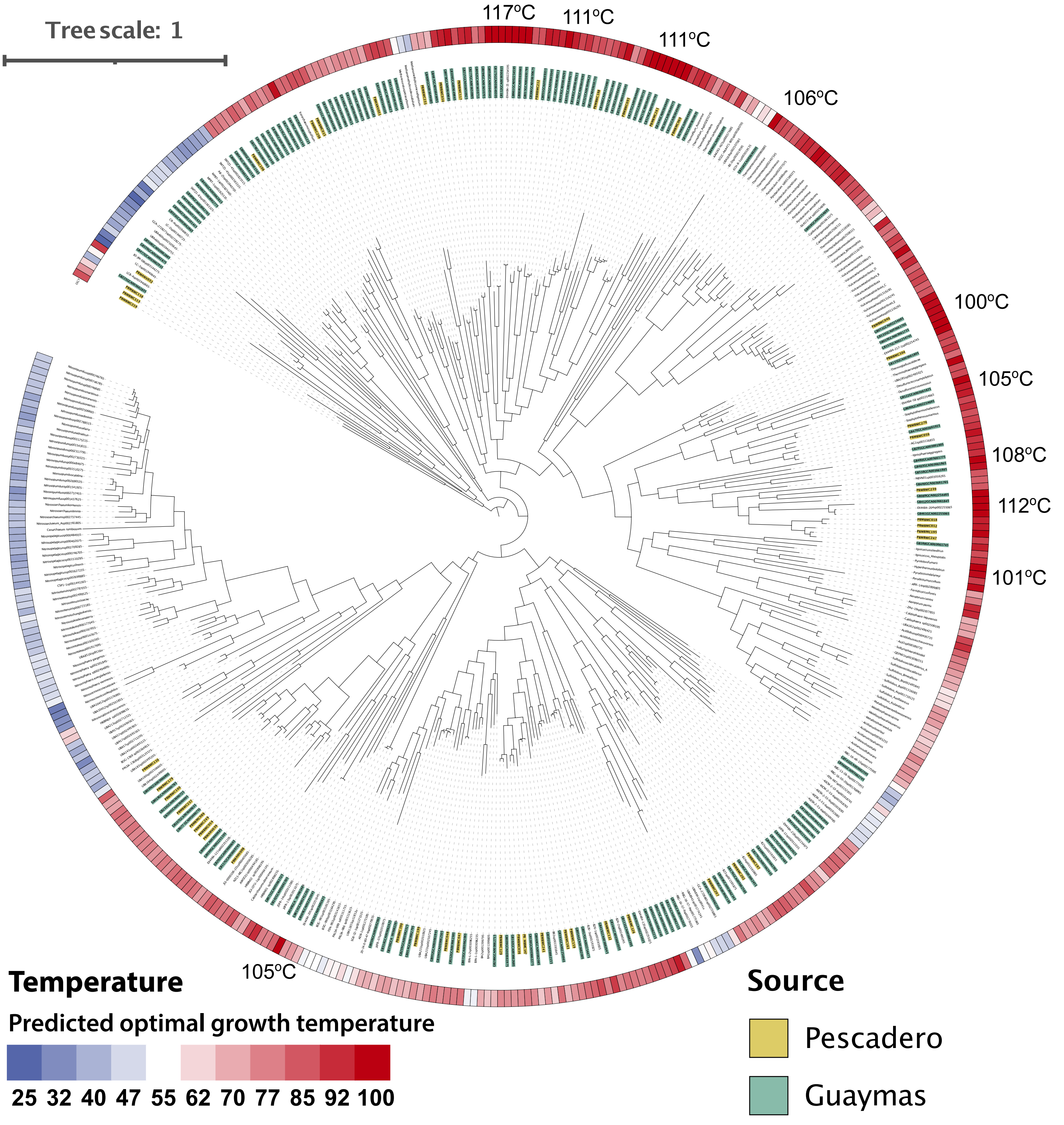


**Supplemental Figure S33. TACK phylogeny and predicted optimal growth temperature**

Concatenated marker gene phylogeny of all *Asgardarchaeota* and *Crenarchaeota* genomes from the genome taxonomy database (GTDB, v89), Guaymas basin (PRJNA362212) and Pescadero Basin (Auka, this study). The phylogeny was calculated using FastTree, on a concatenated alignment based on 76 Archaeal marker genes retrieved from the genomes using Anvi'o and aligned using Muscle. Optimal growth temperature (OGT) was predicted using the OGT prediction algorithm by Sauer and Wang (https://doi.org/10.1093/bioinformatics/btz059). Predicted OGTs above the scale maximum of 100°C are indicated.

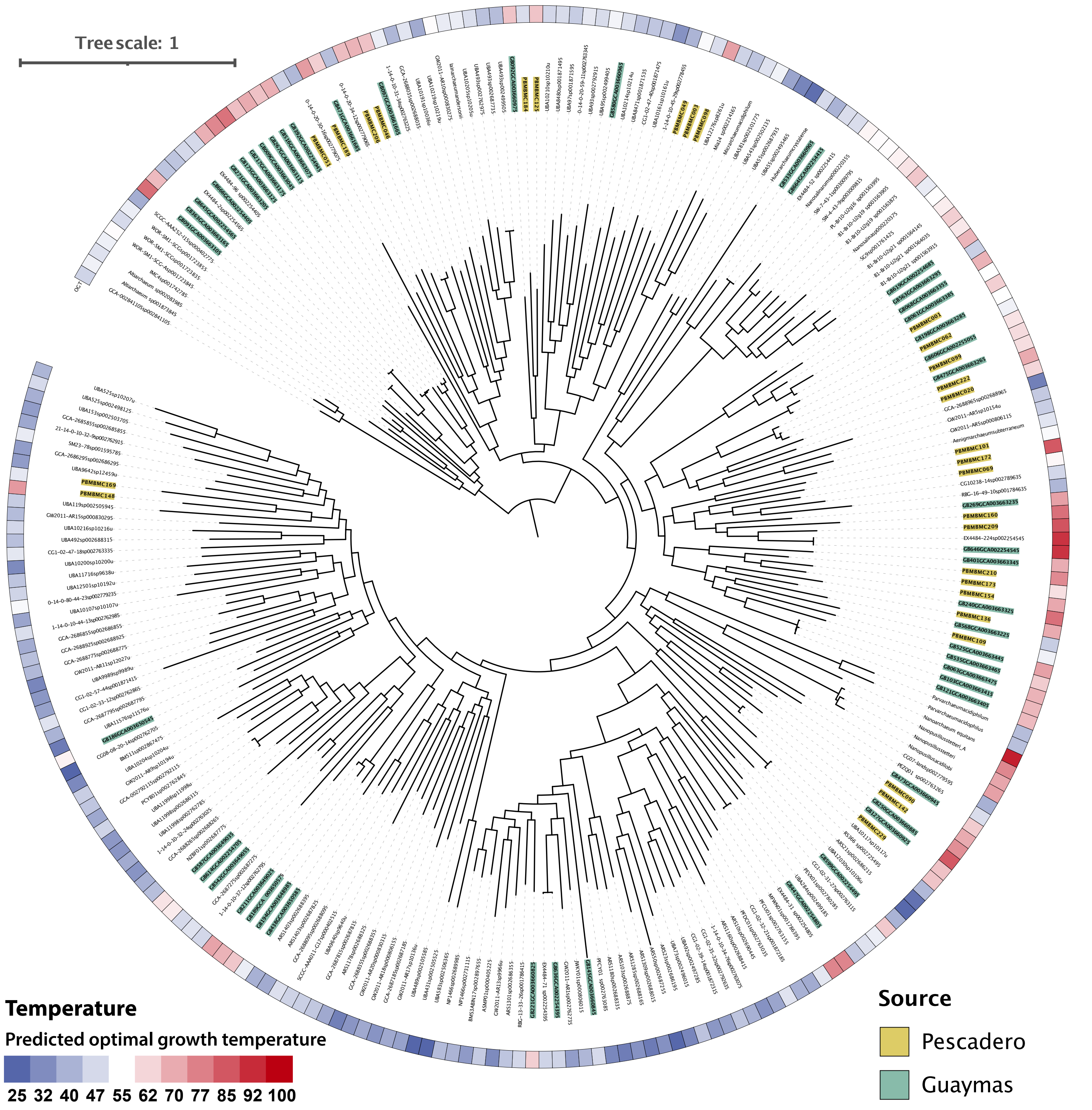


**Supplemental Figure S34. DPANN phylogeny and predicted optimal growth temperature**

Concatenated marker gene phylogeny of all *Aenigmarchaeota, Altiarchaeota,* EX4484_52, *Huberarchaeaota, Iainarchaeota, Micrarchaeota, Nanoarchaeota, Nanohaloarchaeota,* and UAP2 genomes from the genome taxonomy database (GTDB, v89), Guaymas basin (PRJNA362212) and Pescadero Basin (Auka, this study). The phylogeny was calculated using FastTree, on a concatenated alignment based on 76 Archaeal marker genes retrieved from the genomes using Anvi'o and aligned using Muscle. Optimal growth temperature was predicted using the OGT prediction by Sauer and Wang (https://doi.org/10.1093/bioinformatics/btz059).

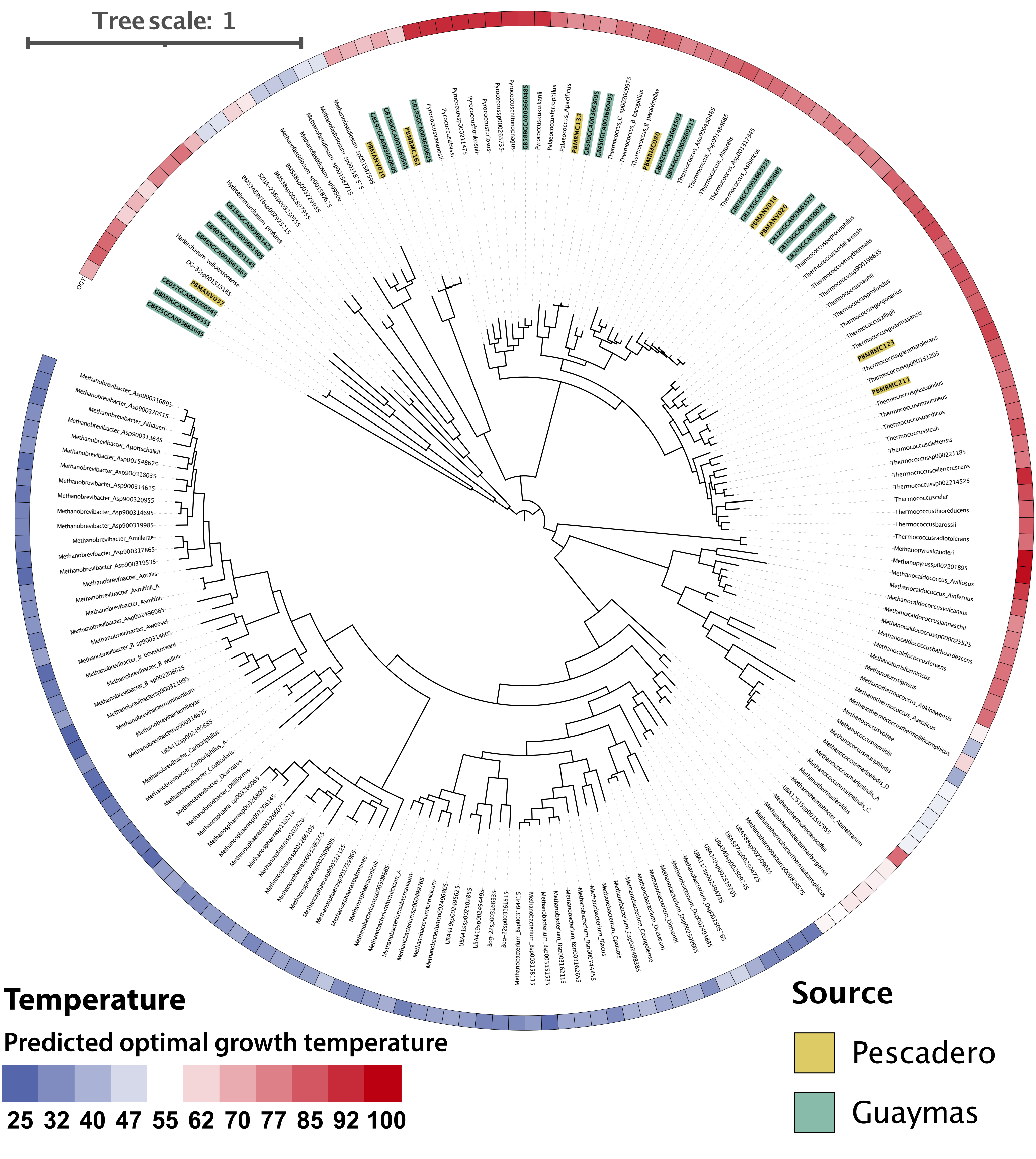


**Supplemental Figure S35. Euryarchaeota phylogeny and predicted optimal growth temperature**

Concatenated marker gene phylogeny of all *Euryarchaeota, Hadarchaeota,* and *Hydrothermarchaeota* genomes from the genome taxonomy database (GTDB, v89), Guaymas basin (PRJNA362212) and Pescadero Basin (Auka, this study). The phylogeny was calculated using FastTree, on a concatenated alignment based on 76 Archaeal marker genes retrieved from the genomes using Anvi'o and aligned using Muscle. Optimal growth temperature was predicted using the OGT prediction by Sauer and Wang (https://doi.org/10.1093/bioinformatics/btz059).

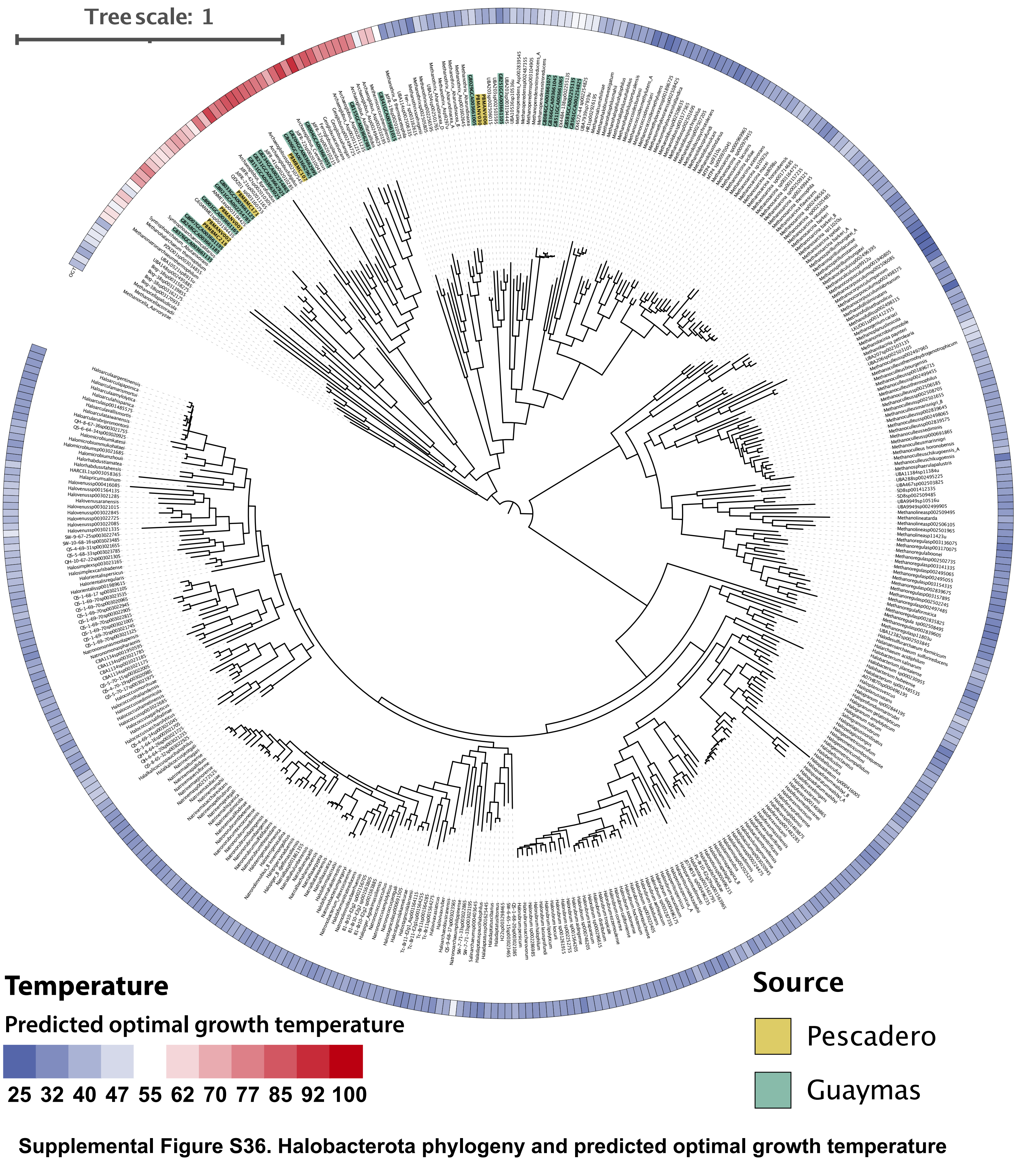


**Supplemental Figure S36. Halobacterota phylogeny and predicted optimal growth temperature**

Concatenated marker gene phylogeny of all *Halobacterota* genomes from the genome taxonomy database (GTDB, v89), Guaymas basin (PRJNA362212) and Pescadero Basin (Auka, this study). The phylogeny was calculated using FastTree, on a concatenated alignment based on 76 Archaeal marker genes retrieved from the genomes using Anvi'o and aligned using Muscle. Optimal growth temperature was predicted using the OGT prediction by Sauer and Wang (https://doi.org/10.1093/bioinformatics/btz059).

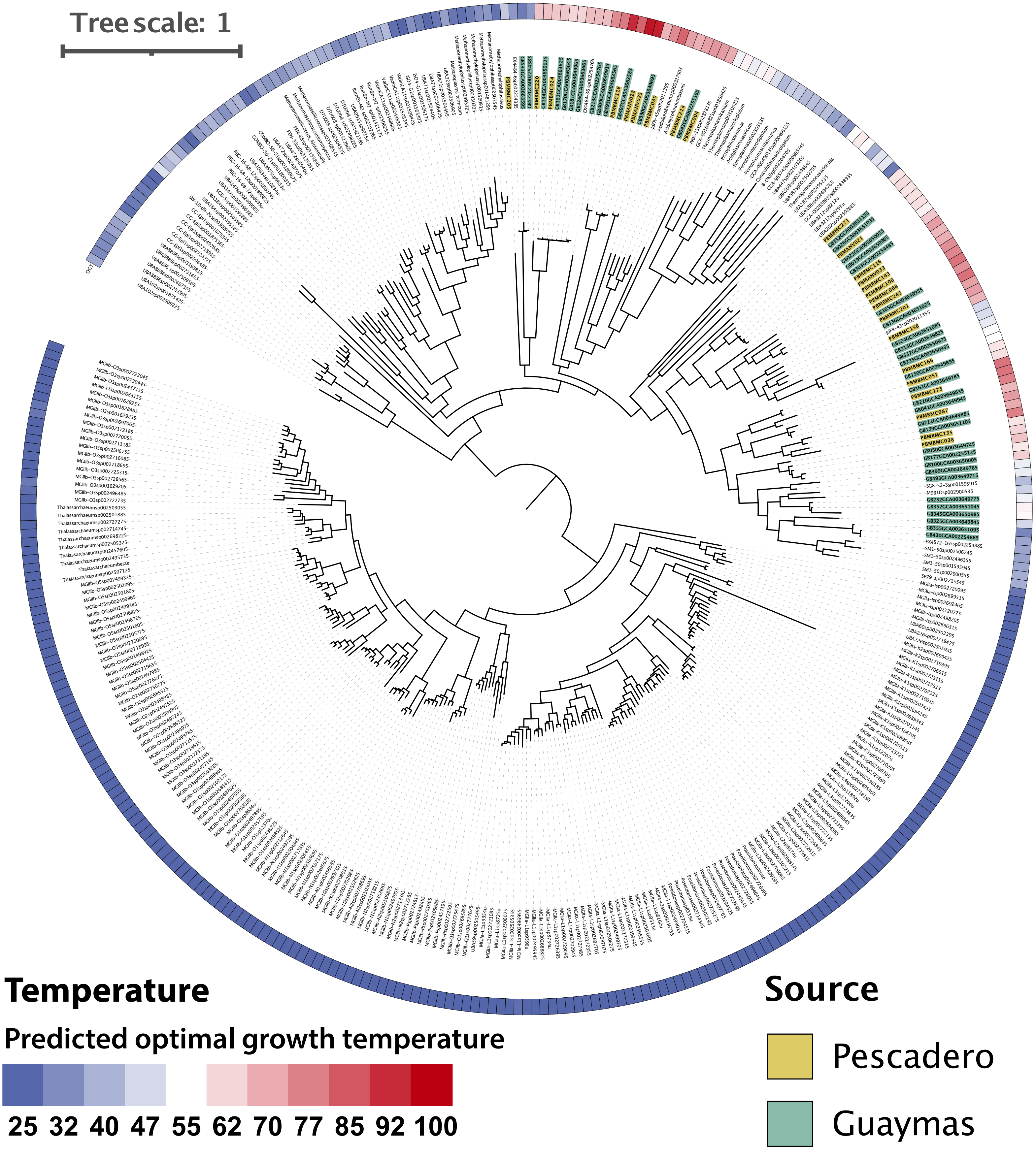


**Supplemental Figure S37. Thermoplasmatota phylogeny and predicted optimal growth temperature**

Concatenated marker gene phylogeny of all *Thermoplasmatota* genomes from the genome taxonomy database (GTDB, v89), Guaymas basin (PRJNA362212) and Pescadero Basin (Auka, this study). The phylogeny was calculated using FastTree, on a concatenated alignment based on 76 Archaeal marker genes retrieved from the genomes using Anvi'o and aligned using Muscle. Optimal growth temperature was predicted using the OGT prediction by Sauer and Wang (https://doi.org/10.1093/bioinformatics/btz059).

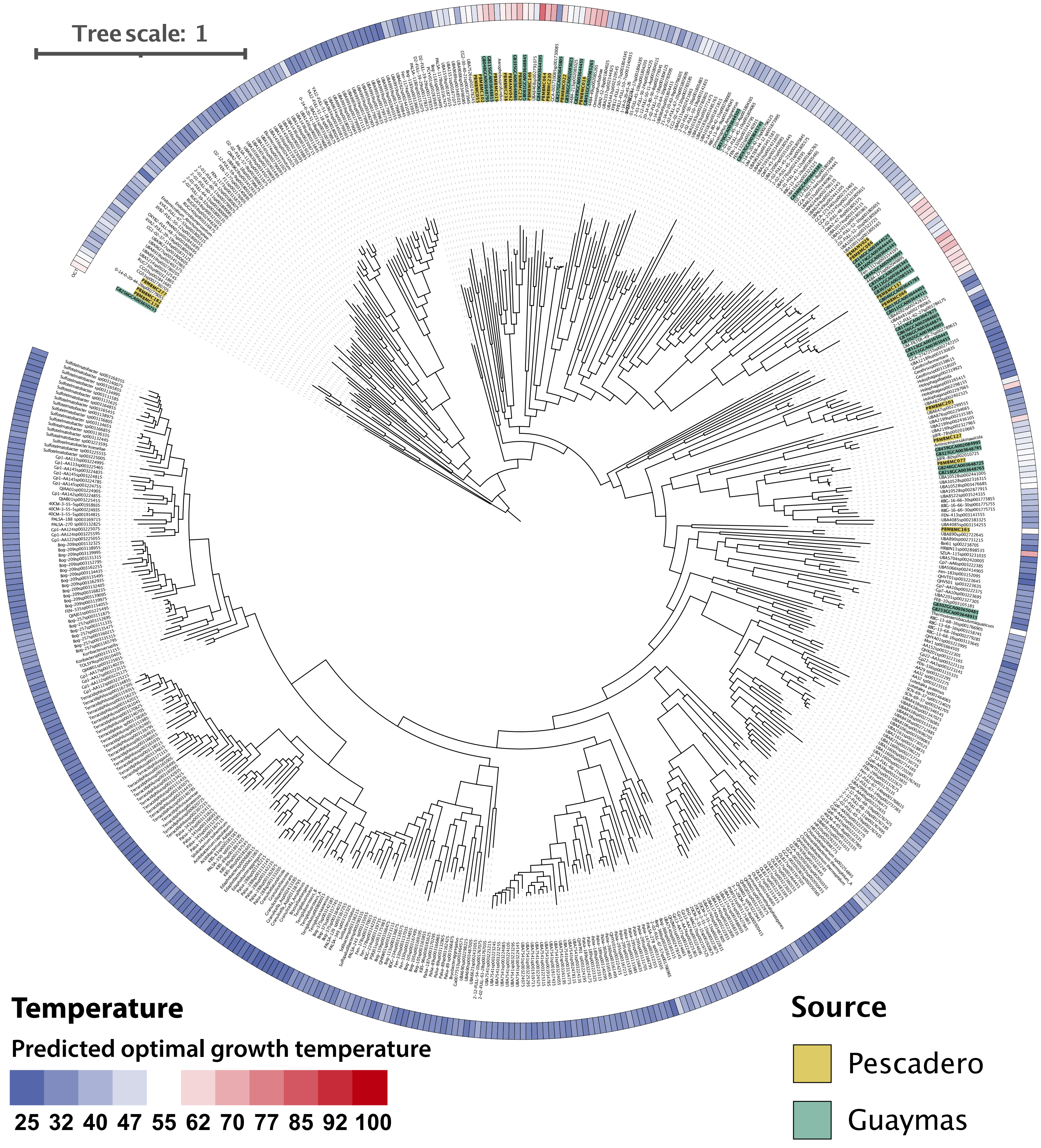


**Supplemental Figure S38. Acidobacteriota phylogeny and predicted optimal growth temperature**

Concatenated marker gene phylogeny of all *Acidobacteriota, Aerophobota,* CG03*, Desantisbacteria, Elusimicrobiota,* FCPU426*, Firestonebacteria,* GCA-001730085*, Goldbacteria, Lindowbacteria, Omnitrophota, Ratteibacteria,* RUG730, UBA6262, UBA8481, UBA9089, UBP18, and UBP4 genomes from the genome taxonomy database (GTDB, v89), Guaymas basin (PRJNA362212) and Pescadero Basin (Auka, this study). The phylogeny was calculated using FastTree, on a concatenated alignment based on 71 Bacterial marker genes retrieved from the genomes using Anvi'o and aligned using Muscle. Optimal growth temperature was predicted using the OGT prediction by Sauer and Wang (https://doi.org/10.1093/bioinformatics/btz059).

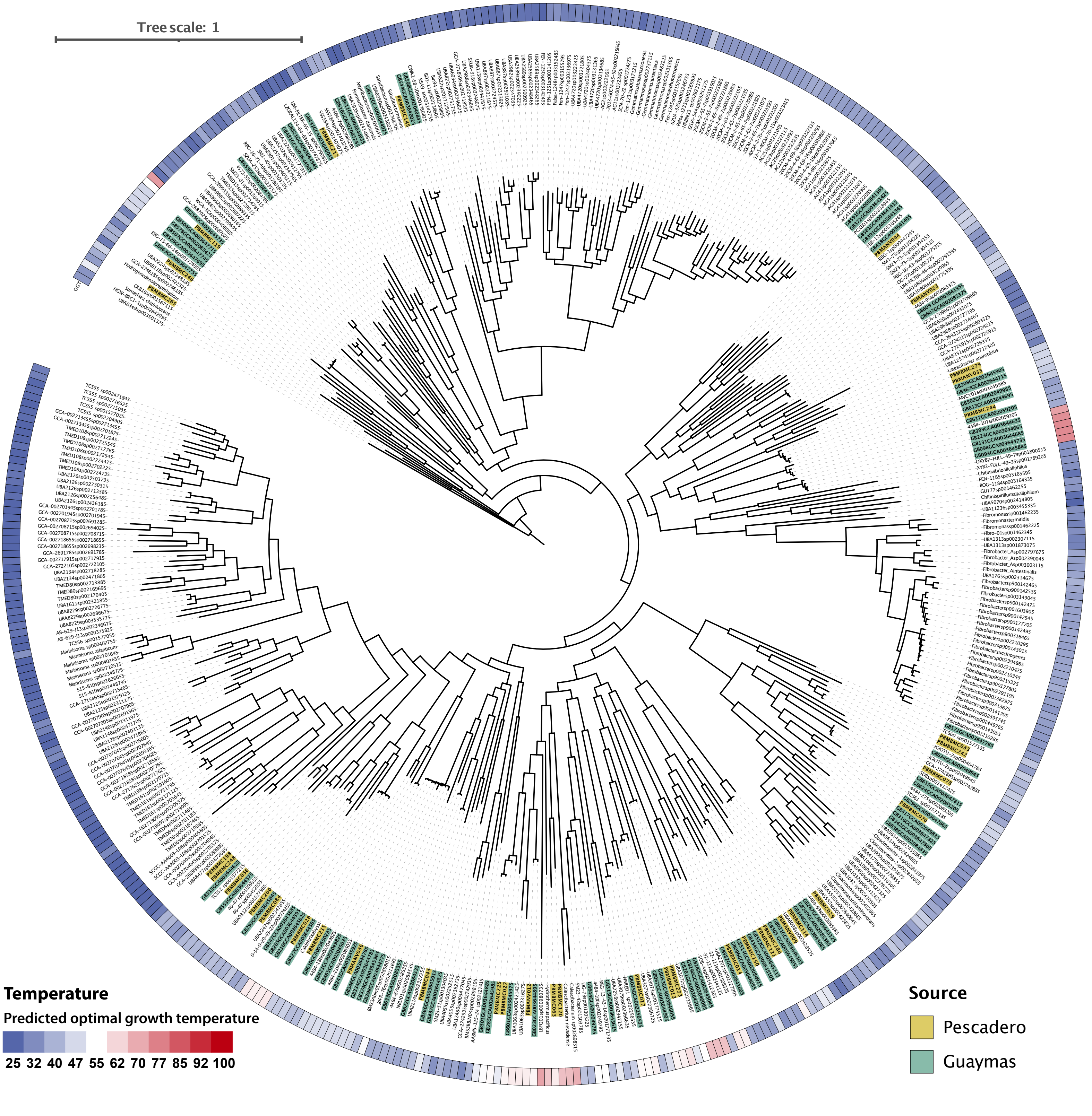


**Supplemental Figure S39. Phylogeny and predicted optimal growth temperature of phyla basal to the Bacteroidota**

Concatenated marker gene phylogeny of all 4572_22, AABM5-125-24, *Calditrichota, Calescibacterota, Cloacimonadota, Delongbacteria, Edwardsbacteria, Eisenbacteria, Fermentibacterota, Fibrobacterota, Gemmatimonadota, Hydrogenedentota, Krumholzibacteriota,* KSB1, *Latescibacterota, Marinisomatota,* OLB16, *Poribacteria, RBG-13-66-14, SM23-31, Sumerlaeota,* TA06_A, TA06, UBP14, WOR-3_A, WOR-3_B, WOR-3, and *Zixibacteria* genomes from the genome taxonomy database (GTDB, v89), Guaymas basin (PRJNA362212) and Pescadero Basin (Auka, this study). The phylogeny was calculated using FastTree, on a concatenated alignment based on 71 Bacterial marker genes retrieved from the genomes using Anvi'o and aligned using Muscle. Optimal growth temperature was predicted using the OGT prediction by Sauer and Wang (https://doi.org/10.1093/bioinformatics/btz059).

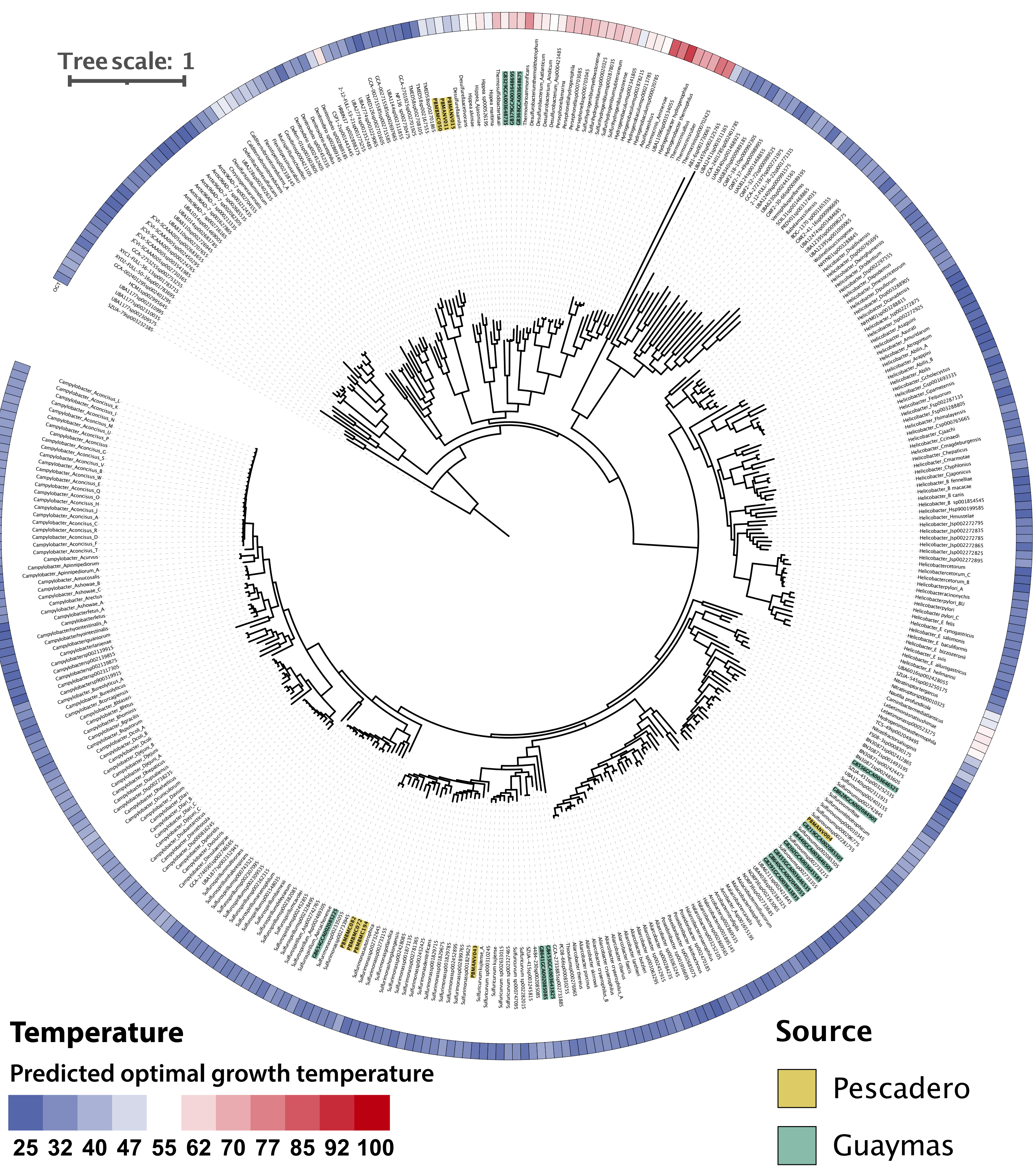


**Supplemental Figure S40. Campylobacterota phylogeny and predicted optimal growth temperature**

Concatenated marker gene phylogeny of all AB1-6, *Aquificota, Campylobacterota, Chrysiogenetota, Dadabacteria, Deferribacterota, Dependentiae,* SAR324, SZUA-79, *Thermosulfidibacterota,* and UBP6 genomes from the genome taxonomy database (GTDB, v89), Guaymas basin (PRJNA362212) and Pescadero Basin (Auka, this study). The phylogeny was calculated using FastTree, on a concatenated alignment based on 71 Bacterial marker genes retrieved from the genomes using Anvi'o and aligned using Muscle. Optimal growth temperature was predicted using the OGT prediction by Sauer and Wang (https://doi.org/10.1093/bioinformatics/btz059).

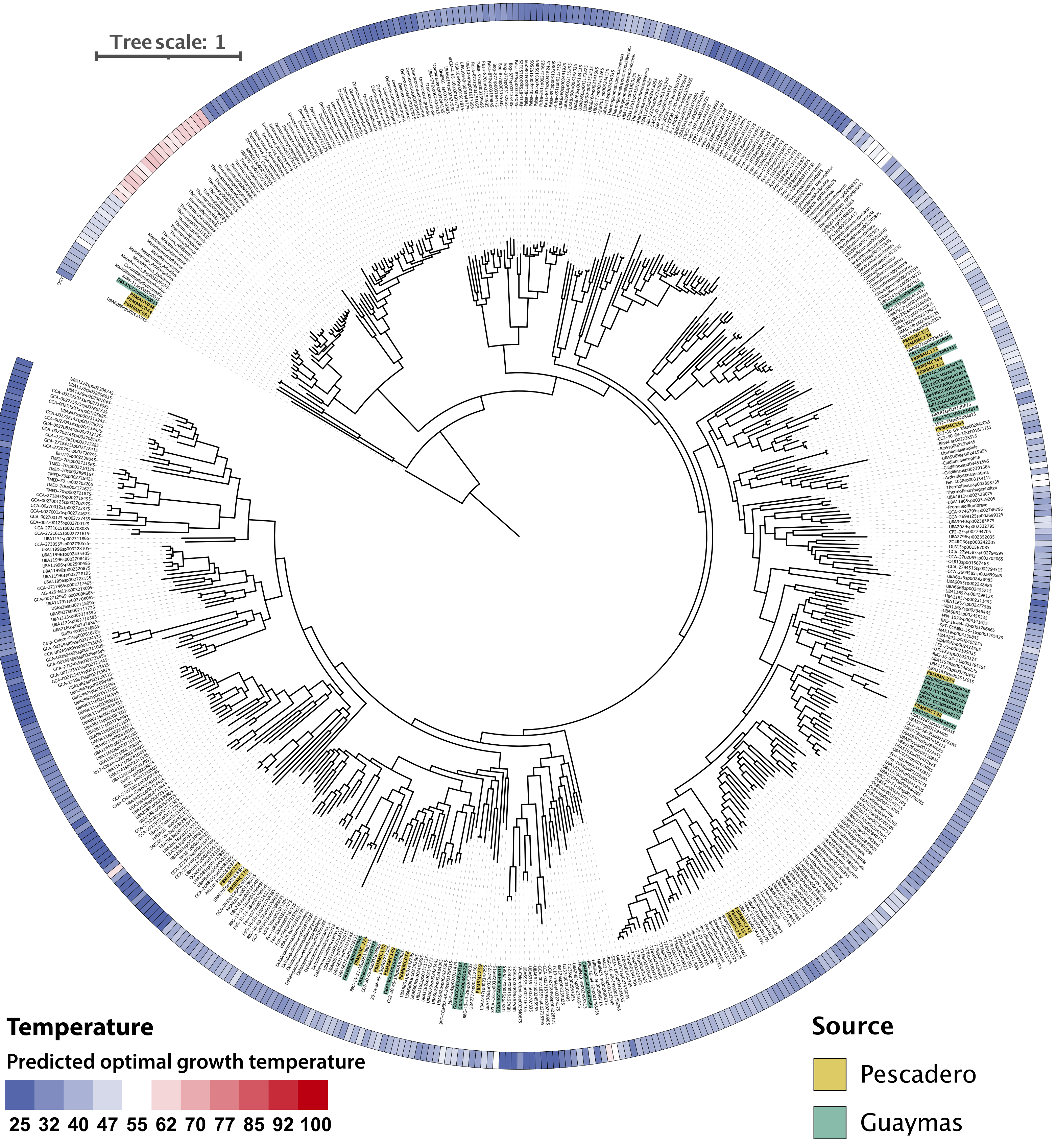


**Supplemental Figure S41. Chloroflexota phylogeny and predicted optimal growth temperature**

Concatenated marker gene phylogeny of all *Chloroflexota*_A, *Chloroflexota*_B, *Chloroflexota*, *Deinococcota*, *Dormibacterota*, UBP15, and UBP7_A genomes from the genome taxonomy database (GTDB, v89), Guaymas basin (PRJNA362212) and Pescadero Basin (Auka, this study). The phylogeny was calculated using FastTree, on a concatenated alignment based on 71 Bacterial marker genes retrieved from the genomes using Anvi'o and aligned using Muscle. Optimal growth temperature was predicted using the OGT prediction by Sauer and Wang (https://doi.org/10.1093/bioinformatics/btz059).

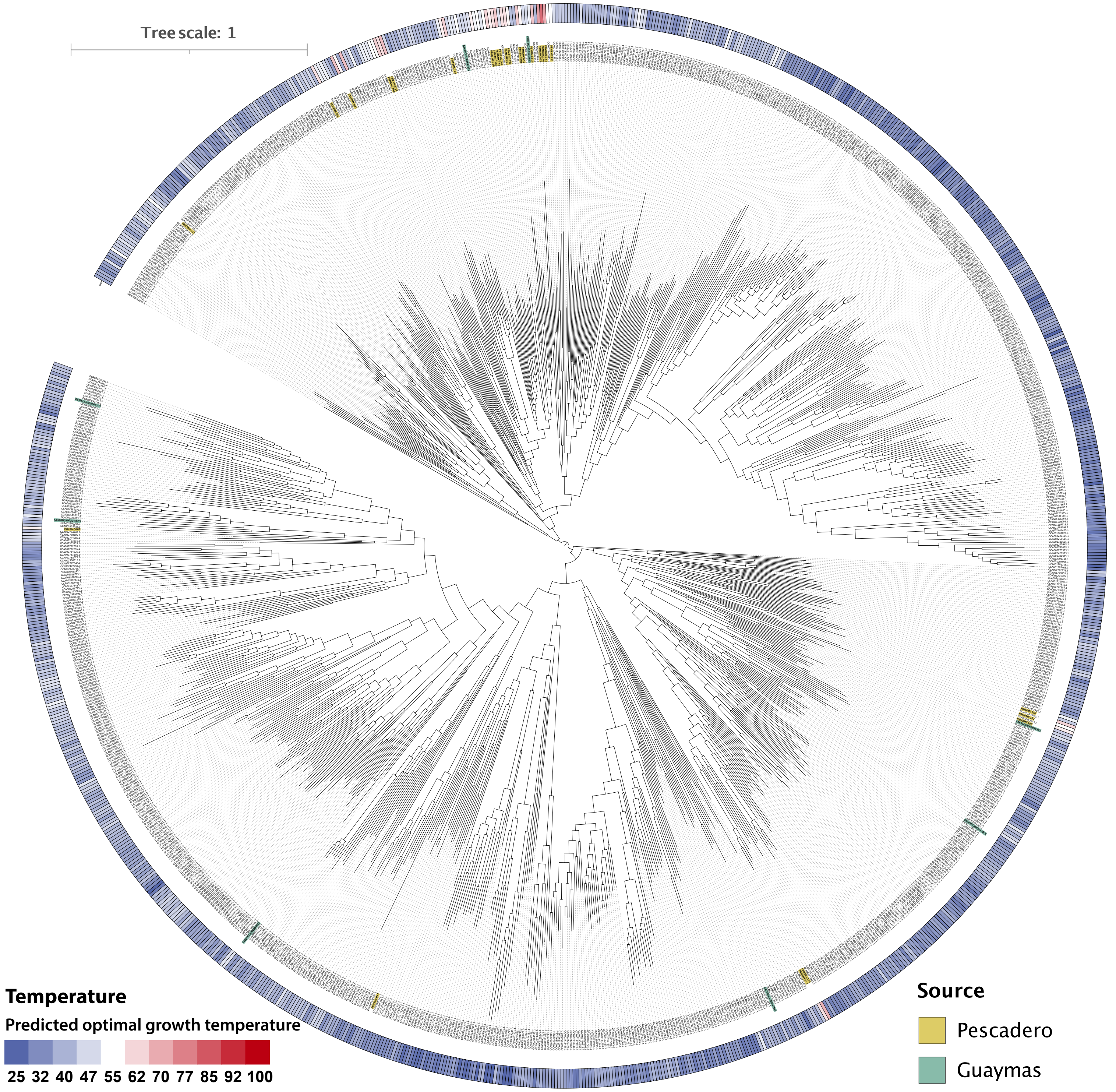


**Supplemental Figure S42. Patescibacteria (CPR) phylogeny and predicted optimal growth temperature**

Concatenated marker gene phylogeny of all *Patescibacteria* (CPR) genomes from the genome taxonomy database (GTDB, v89), Guaymas basin (PRJNA362212) and Pescadero Basin (Auka, this study). The phylogeny was calculated using FastTree, on a concatenated alignment based on 71 Bacterial marker genes retrieved from the genomes using Anvi'o and aligned using Muscle. Optimal growth temperature was predicted using the OGT prediction by Sauer and Wang (https://doi.org/10.1093/bioinformatics/btz059).

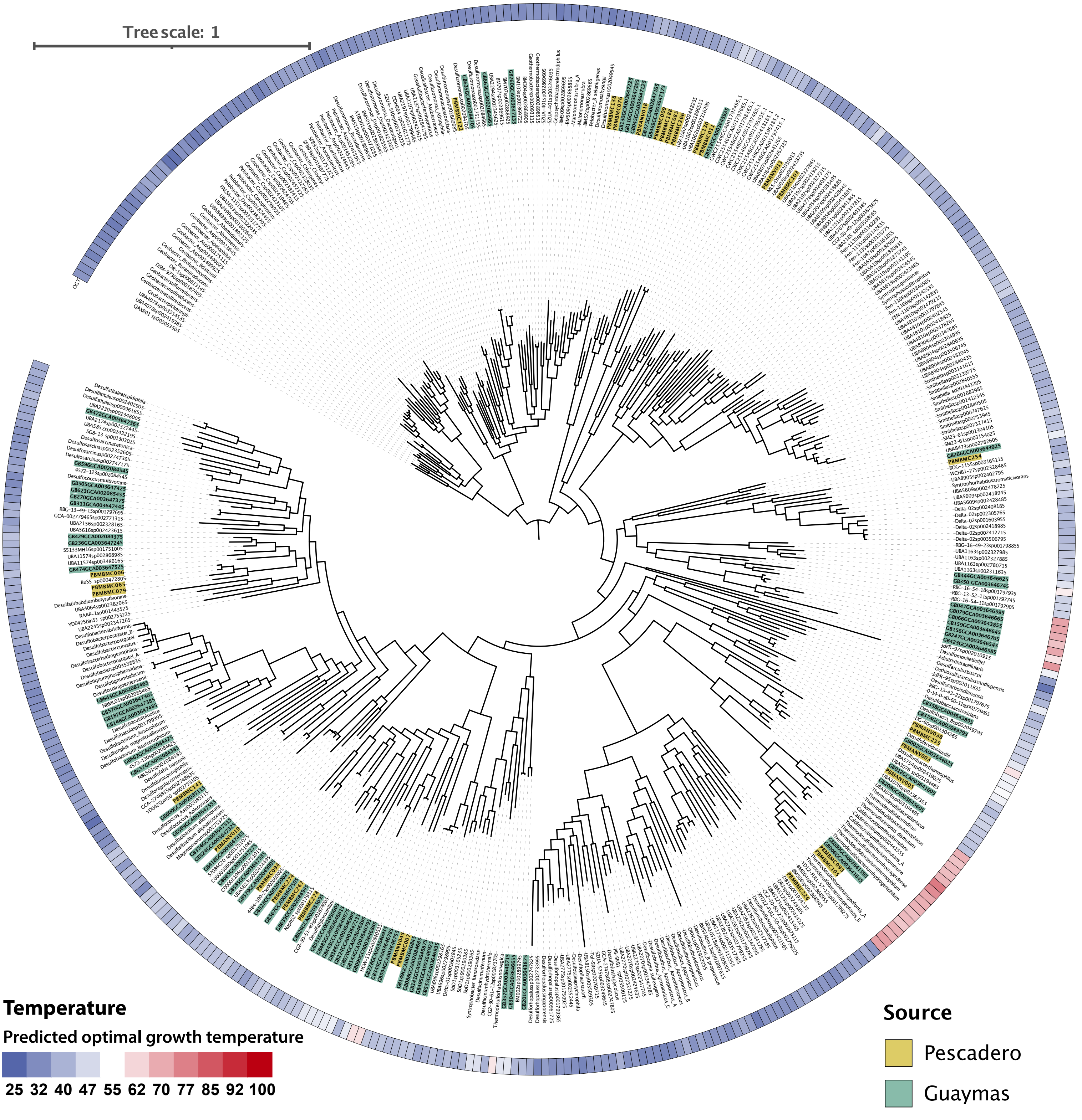


**Supplemental Figure S43. Desulfobacterota phylogeny and predicted optimal growth temperature**

Concatenated marker gene phylogeny of all *Desulfobacterota_A, Desulfobacterota, Desulfuromonadota,* and GWC2-55-46 genomes from the genome taxonomy database (GTDB, v89), Guaymas basin (PRJNA362212) and Pescadero Basin (Auka, this study). The phylogeny was calculated using FastTree, on a concatenated alignment based on 71 Bacterial marker genes retrieved from the genomes using Anvi'o and aligned using Muscle. Optimal growth temperature was predicted using the OGT prediction by Sauer and Wang (https://doi.org/10.1093/bioinformatics/btz059).

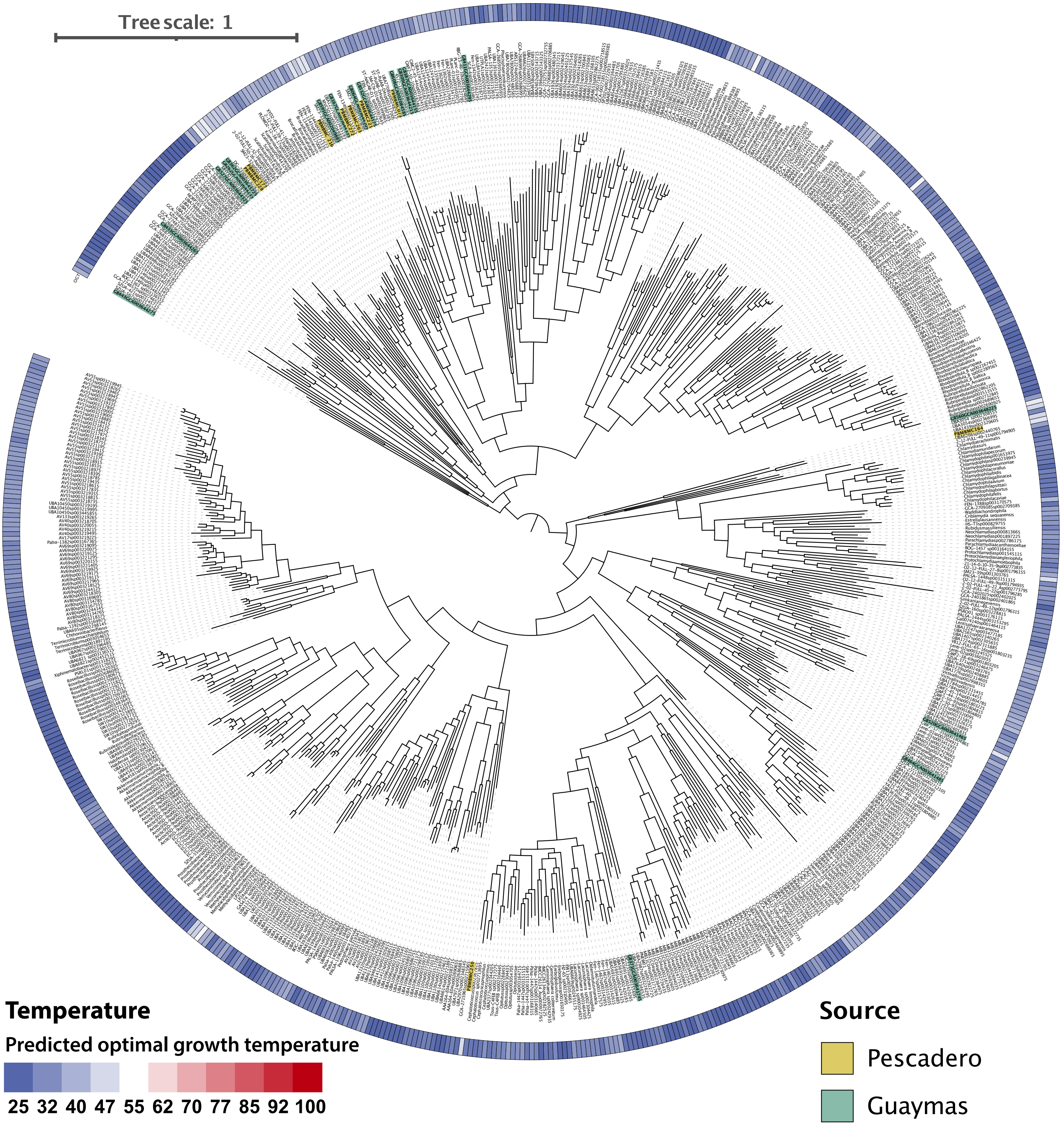


**Supplemental Figure S44. PVC phylogeny and predicted optimal growth temperature**

Concatenated marker gene phylogeny of all *Planctomycetota,* UBP3*, Verrucomicrobiota*_A, and *Verrucomicrobiota* genomes from the genome taxonomy database (GTDB, v89), Guaymas basin (PRJNA362212) and Pescadero Basin (Auka, this study). The phylogeny was calculated using FastTree, on a concatenated alignment based on 71 Bacterial marker genes retrieved from the genomes using Anvi'o and aligned using Muscle. Optimal growth temperature was predicted using the OGT prediction by Sauer and Wang (https://doi.org/10.1093/bioinformatics/btz059).

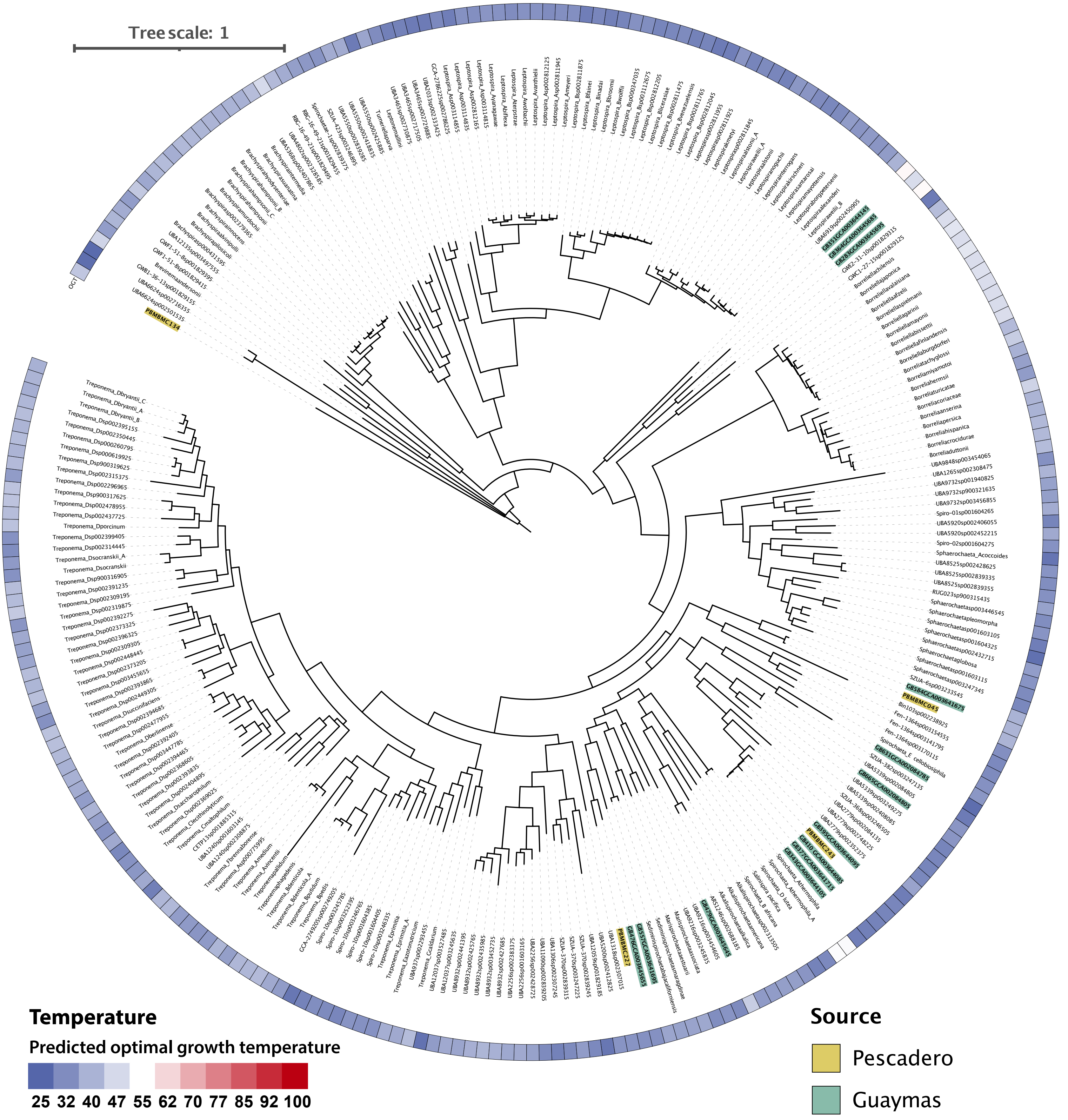


**Supplemental Figure S45. Spirochaetota phylogeny and predicted optimal growth temperature**

Concatenated marker gene phylogeny of all *Spirochaetota* and UBP7 genomes from the genome taxonomy database (GTDB, v89), Guaymas basin (PRJNA362212) and Pescadero Basin (Auka, this study). The phylogeny was calculated using FastTree, on a concatenated alignment based on 71 Bacterial marker genes retrieved from the genomes using Anvi'o and aligned using Muscle. Optimal growth temperature was predicted using the OGT prediction by Sauer and Wang (https://doi.org/10.1093/bioinformatics/btz059).

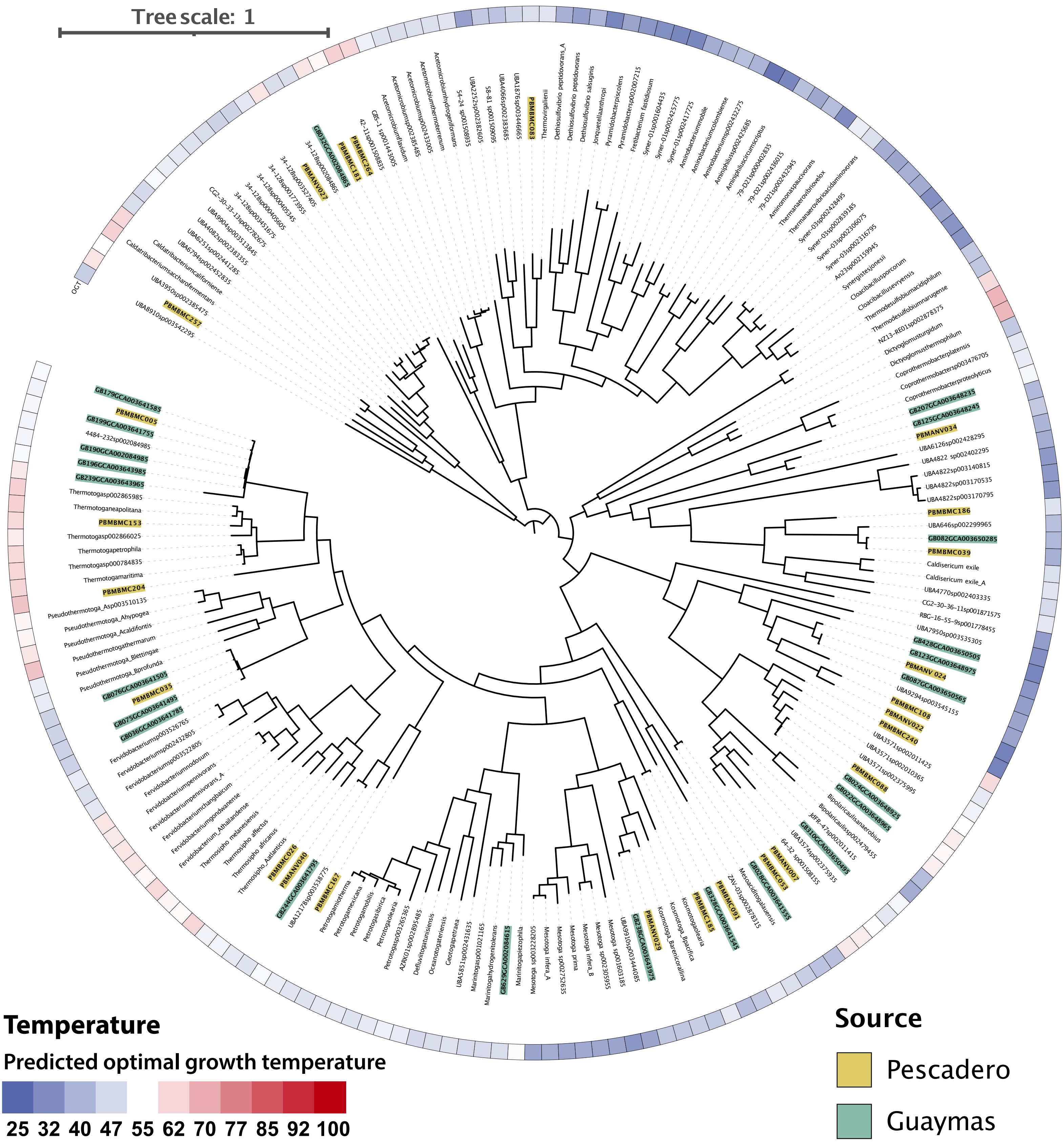


**Supplemental Figure S46. Thermotogota phylogeny and predicted optimal growth temperature**

Concatenated marker gene phylogeny of all *Bipolaricaulota, Caldatribacteriota, Caldisericota, Coprothermobacterota, Dictyoglomota, Synergistota, Thermodesulfobiota,* and *Thermotogota* genomes from the genome taxonomy database (GTDB, v89), Guaymas basin (PRJNA362212) and Pescadero Basin (Auka, this study). The phylogeny was calculated using FastTree, on a concatenated alignment based on 71 Bacterial marker genes retrieved from the genomes using Anvi'o and aligned using Muscle. Optimal growth temperature was predicted using the OGT prediction by Sauer and Wang (https://doi.org/10.1093/bioinformatics/btz059).

**Supplemental text**

Distinct sulfur oxidizing communities at Guaymas Basin and Auka

In stark contrast to the lineages shared between Auka and Guaymas Basin (see main text), a notable difference between the microbial communities of Auka vent field and Guaymas Basin are the sulfur oxidizing Bacteria (SOB) comprising the microbial mats on the sediment surface. These mats are dominated by *Beggiatoa* in Guaymas Basin (Gundersen et al. 1992; McKay et al. 2012) (~1700m depth) whereas the mat microbial community in the deeper Auka vent field (>3600m depth) is dominated by *Campylobacterota* (formerly *Epsilonproteobacteria*) related to *Sulfurimonas (Inagaki et al. 2003)* and *Sulfurovum (Inagaki et al. 2004)* (Supplemental figure S40). A similar depth trend was observed at the ultramafic hosted Mid-Cayman rise vents with *Gammaproteobacteria* (*Beggiatoa* and *Thiothrix*) dominating the shallower Von Damm community (2450m depth), and *Sulfurovum* dominant at the deeper Piccard vents (4950m depth) (Anderson et al. 2017). Yamamoto and Takai previously hypothesized *Campylobacterota* SOB have a wider ecophysiological range than *Gammaproteobacteria* SOB based on their ability to both oxidize and reduce sulfur cycle intermediates (Yamamoto and Takai 2011), suggesting differences in fluid chemistry could account for the observed depth trend. However, given the striking differences in hydrothermal fluid composition between sediment-hosted and ultramafic-hosted vents (McDermott et al. 2018), it is unclear whether fluid composition can explain the depth trend observed at both locations.

**References to the Supplemental text**


Anderson, Rika E., Julie Reveillaud, Emily Reddington, Tom O. Delmont, A. Murat Eren, Jill M. McDermott, Jeff S. Seewald, and Julie A. Huber. 2017. “Genomic Variation in Microbial Populations Inhabiting the Marine Subseafloor at Deep-Sea Hydrothermal Vents.” *Nature Communications* 8 (1): 1114.

Gundersen, Jens K., Bo Barker Jorgensen, Einer Larsen, and Holger W. Jannasch. 1992. “Mats of Giant Sulphur Bacteria on Deep-Sea Sediments due to Fluctuating Hydrothermal Flow.” *Nature* 360 (6403): 454–56.

Inagaki, Fumio, Ken Takai, Hideki Kobayashi, Kenneth H. Nealson, and Koki Horikoshi. 2003. “Sulfurimonas Autotrophica Gen. Nov., Sp. Nov., a Novel Sulfur-Oxidizing Epsilon-Proteobacterium Isolated from Hydrothermal Sediments in the Mid-Okinawa Trough.” *International Journal of Systematic and Evolutionary Microbiology* 53 (Pt 6): 1801–5.

Inagaki, Fumio, Ken Takai, Kenneth H. Nealson, and Koki Horikoshi. 2004. “Sulfurovum Lithotrophicum Gen. Nov., Sp. Nov., a Novel Sulfur-Oxidizing Chemolithoautotroph within the Epsilon-Proteobacteria Isolated from Okinawa Trough Hydrothermal Sediments.” *International Journal of Systematic and Evolutionary Microbiology* 54 (Pt 5): 1477–82.

McDermott, Jill M., Sean P. Sylva, Shuhei Ono, Christopher R. German, and Jeffrey S. Seewald. 2018. “Geochemistry of Fluids from Earth’s Deepest Ridge-Crest Hot-Springs: Piccard Hydrothermal Field, Mid-Cayman Rise.” *Geochimica et Cosmochimica Acta* 228 (May): 95–118.

McKay, Luke J., Barbara J. MacGregor, Jennifer F. Biddle, Daniel B. Albert, Howard P. Mendlovitz, Daniel R. Hoer, Julius S. Lipp, Karen G. Lloyd, and Andreas P. Teske. 2012.

“Spatial Heterogeneity and Underlying Geochemistry of Phylogenetically Diverse Orange and White Beggiatoa Mats in Guaymas Basin Hydrothermal Sediments.” *Deep Sea Research Part I: Oceanographic Research Papers* 67 (September): 21–31.

Yamamoto, Masahiro, and Ken Takai. 2011. “Sulfur Metabolisms in Epsilon- and Gamma-Proteobacteria in Deep-Sea Hydrothermal Fields.” *Frontiers in Microbiology* 2 (September): 192.
